# Genome Mining Uncovers Clustered Biosynthetic Pathways for Defense-Related Molecules in Bread Wheat

**DOI:** 10.1101/2021.11.04.467362

**Authors:** Guy Polturak, Martin Dippe, Michael J Stephenson, Rajesh Chandra Misra, Charlotte Owen, Ricardo Ramirez-Gonzalez, John Haidioulis, Henk-Jan Schoonbeek, Laetitia Chartrain, Philippa Borrill, David R Nelson, James Brown, Paul Nicholson, Cristobal Uauy, Anne Osbourn

## Abstract

Wheat is one of the most widely grown food crops in the world. However, it succumbs to numerous pests and pathogens that cause substantial yield losses. A better understanding of biotic stress responses in wheat is thus of major importance. Here we identify previously unknown pathogen-induced biosynthetic pathways that produce a diverse set of molecules, including flavonoids, diterpenes and triterpenes. These pathways are encoded by six biosynthetic gene clusters and share a common regulatory network. We further identify associations with known or novel phytoalexin clusters in other cereals and grasses. Our results significantly advance the understanding of chemical defenses in wheat and open up new avenues for enhancing disease resistance in this agriculturally important crop.

## INTRODUCTION

The allohexaploid bread wheat (*Triticum aestivum*) accounts for approximately 20% of the calories consumed by humans worldwide^1^. Around one fifth of the global annual wheat yield is lost due to pest and pathogen attack^2^, a value that is expected to sharply rise as the climate warms^3,4^. A better understanding of how wheat responds to biotic stresses could enable the development of strategies for minimizing yield losses and reducing reliance on pesticides. Significant advances have been made in identification of wheat resistance genes (R-genes) involved in pathogen recognition and the immune response^5^. However, very little is known about the chemical defenses produced by wheat in response to pathogen attack (phytoalexins).

The agronomic importance of wheat has led to extensive research into its genetics, and to the generation of a vast body of transcriptome data from numerous studies into wheat development, physiology, and interactions with the environment. However, the first bread wheat genome assembly became available only recently because of the challenges associated with its large genome size, high repetitive sequence content, and relatedness between homoeologous subgenomes^6^. The availability of these genome and transcriptomic resources now offers the opportunity to employ a genomics-driven approach to uncover novel chemical defense molecules and biosynthetic pathways in this valuable crop. Such an approach is particularly useful for uncovering metabolites that are produced in small quantities or under specific conditions (e.g. pathogen-induced), thereby eluding traditional chemical analyses^7^.

Here, by coupling gene co-expression network analysis with genome mining, we identify six defense-related candidate biosynthetic gene clusters (BGCs) in bread wheat. We show by expression of cluster genes in *Nicotiana benthamiana* that these BGCs encode pathways for the production of flavonoid, diterpene and triterpene compounds that likely serve as broad-spectrum phytoalexins in wheat. Through comparative genomics we also identify associations with known or novel phytoalexin clusters in other cereals and grasses. We further report the full characterization of the pathways for the novel defence compounds ellarinacin and brachynacin, which are respectively produced by related gene clusters in wheat and the grass purple false brome (*Brachypodium distachyon*). Our work uncovers new biosynthetic pathways for novel pathogen-induced compounds in wheat and demonstrates a powerful approach for rapid discovery of defense-related molecules and metabolic pathways in crop plants, which may have future applications in crop protection.

## RESULTS

### Gene co-expression network analysis coupled with genome analysis identifies candidate pathogen-induced biosynthetic gene clusters in wheat

In a recently published study, 850 transcriptome datasets were compiled and analyzed to produce a genome-wide view of homoeolog expression patterns in hexaploid bread wheat. Weighted Gene Co-expression Network Analysis (WGCNA) was carried out based on gene expression patterns in the compiled datasets, and an additional set of networks were built for six separate subsample sets: grain, leaf, spike, root, abiotic and disease^8^.

We hypothesized that new defense-related metabolites and metabolic pathways in wheat could be found by mining the ‘disease’ gene network. Specifically, genes that are physically clustered in the genome and are co-induced by pathogens or pathogen-associated molecules could serve as excellent candidates for biosynthesis of defense compounds^7^. WGCNA assigned 55,646 genes from the ‘disease’ network (generated from 163 RNA-seq samples) into 69 modules based on their expression, and expression values of all genes in each module were averaged to get a single ‘eigengene’ expression pattern per module^8^. To find genes that exhibit a general, non-specific induction by exposure to pathogens or pathogen-associated molecular patterns (PAMPs), we averaged for each module the difference in normalized eigengene expression between treatment and control in seven different studies and sorted the modules by the average expression delta (Fig. 1a). The top five modules (i.e., the modules represented by the most highly induced eigengenes), namely ME34, ME25, ME12, ME36 and ME8, showed consistent induction in all seven experiments used in the analysis (Fig. 1b) and were selected for further investigation.

**Fig. 1:**
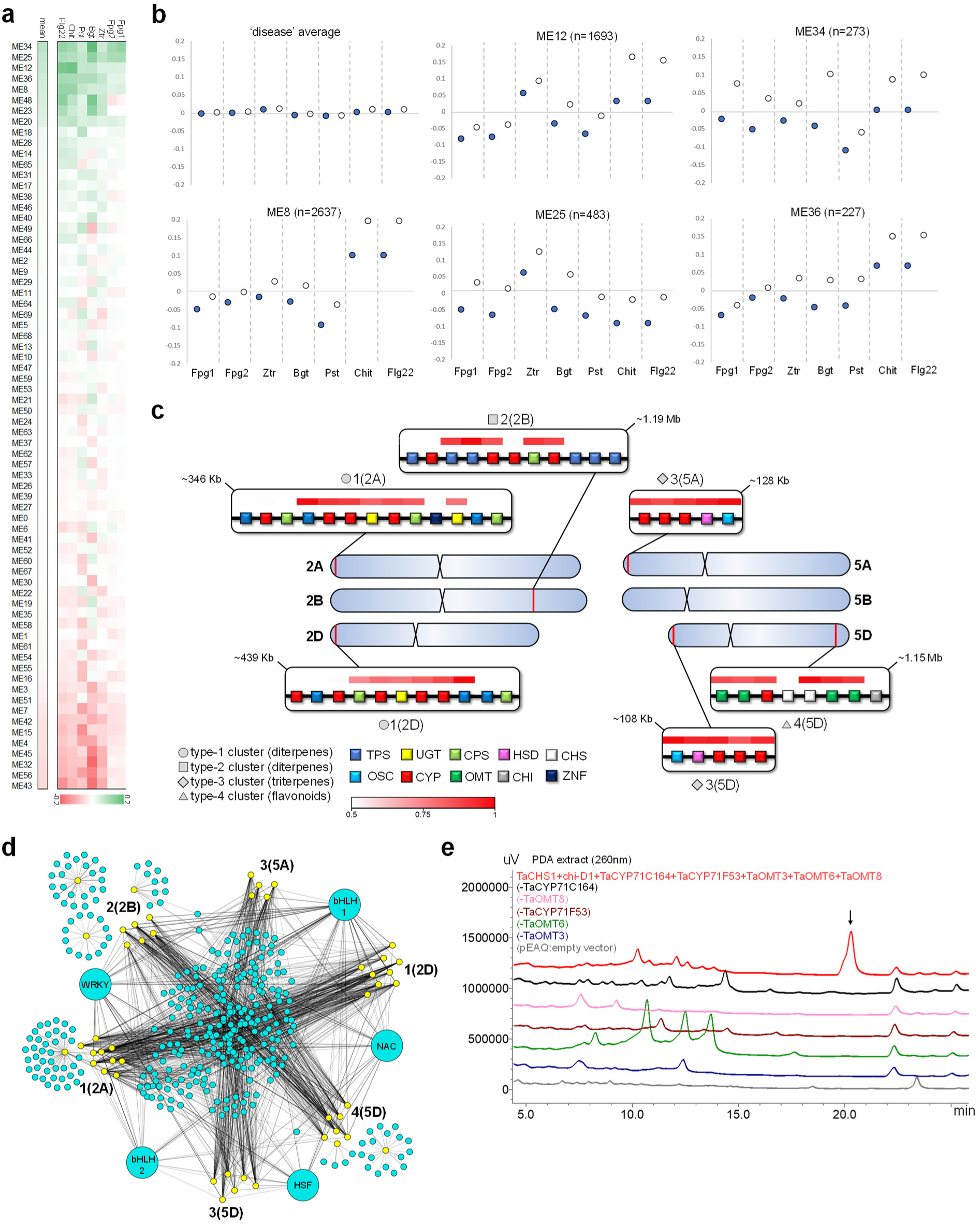
Co-expression network analysis reveals candidate defense-related BGCs in bread wheat. **a**, differential expression values of eigengenes representing 69 gene expression modules generated by Weighted Gene Co-expression Network Analysis (WGCNA) of a ‘disease’ subset of wheat genes. Eigengenes are sorted by mean of delta between treatment and control experiments with different pathogens or PAMPs: Fpg1, *Fusarium pseudograminearum*^77^; Fpg2, *Fusarium pseudograminearum*^78^; Ztr*, Zymoseptoria tritici*^79^; Bgt*, Blumeria graminis f. sp. tritici*^80^; Pst*, Puccinia striiformis f. sp. tritici*^39^; Chit, chitin^8^; Flg22, flagellin peptide^8^. **b**, normalized expression of eigengenes from modules ME34, ME25, ME12, ME36 and ME8, and average of eigengenes from all modules of the ‘disease’ network. The number of genes (n) within each module is indicated. Control and treatment experiments in each study are represented by full and empty circles, respectively. **c**, putative disease-related biosynthetic gene clusters (BGCs) identified in wheat. The red lines indicate the chromosomal positions of the BGCs on the bread wheat chromosomes. The different types of cluster genes are colour-coded according to their annotation: *TPS*, terpene synthase; *OSC*, oxidosqualene cyclase; *UGT*, UDP-glycosyltransferase; *CYP*, cytochrome P450; *CPS*, copalyl diphosphate synthase; *OMT*, O-methyl transferase; *HSD*, hydroxysteroid dehydrogenase; *CHI*, chalcone isomerase; *CHS*, chalcone synthase; *ZNF*, Zinc finger RING/FYVE/PHD-type. Clusters are named according to type (types 1-4) and chromosomal location. The white to red color-coding denotes the Pearson correlation (r) values for expression of each gene with a representative ‘bait’ gene from the cluster. **d**, target gene-transcription factor interactions derived from a GENIE3-based wheat regulatory network. Yellow nodes represent target genes from six BGCs. Light blue nodes represent transcription factors interacting with one or more target genes. Pairwise interaction weight is denoted by edge width. Representative TFs from the most highly interacting TF groups (based on sum of all interaction weights) are enlarged and annotated: bHLH1 (TraesCS3B01G122800), NAC (TraesCS5A01G411800), HSF (TraesCS1A01G350400), bHLH2 (TraesCS7D01G360600), WRKY (TraesCS2D01G011700). **e**, LC-PDA analysis of the products of cluster 4(5D) genes following transient *Agrobacterium*-mediated expression in *N. benthamiana*. The product at Rt=20.3, marked with an arrow, is formed with expression of the complete cluster, but not in the absence of any of the genes *TaCYP71C164_5D*, *TaCYP71F53_5D*, *TaOMT3*, *TaOMT*6, or *TaOMT8*. pEAQ-HT-DEST1, empty vector control.

To determine whether any of these gene expression modules contained genes that form putative biosynthetic gene clusters, we next mined the five modules by filtering for groups of three or more genes with successive accession numbers, i.e. that are physically adjacent in the genome. A total of 55 groups were found (Extended Data Table 1), which include groups of tandem duplicates as expected. Twenty contain protein kinase genes with possible roles in biotic stress responses. A further six consist of genes for different types of enzyme families associated with plant specialized metabolism, and so were identified as possible biosynthetic gene clusters (BGCs) for synthesis of defense compounds. These six putative BGCs included two pairs of homoeologous clusters and were thus defined as four cluster types (cluster types 1-4), and assigned as clusters 1(2A), 1(2D), 2(2B), 3(5A), 3(5D), and 4(5D). The bracketed numbers refer to the chromosomes that the clusters are located on (Fig. 1c, Supplementary Table 1). The two homoeologous cluster pairs are 1(2A) and 1(2D), and 3(5A) and 3(5D), respectively.

The majority of genes in the six putative BGCs were found in a single module, ME25, indicating highly similar expression patterns and suggesting possible co-regulation of the BGCs by a shared network of transcription factors (TFs) (Supplementary Table 1). Analysis of a previously generated GENIE3-based wheat regulatory network^8^ indeed revealed a highly overlapping network of TFs predicted to interact with the six BGCs. Specifically, 137 TFs predicted to interact with genes from two or more of the BGCs were found, including 21 TFs from 10 groups (i.e. groups of homoeologs or tandem duplicates) predicted to interact with genes from all six clusters. The top five most highly interacting TF groups included transcription factors from the WRKY, bHLH (two groups), NAC, and HSF families (Fig. 1d, Extended Data Table 2), all of which have been associated with regulation of phytoalexin biosynthesis or pathogen resistance in plants^9,10^. Examination of Gene Ontology (GO) term enrichment of the predicted target genes of representative TFs from each of the five groups showed that the most significantly enriched terms are related to immune response or defense from biotic stress, for all five TFs excluding the NAC transcription factor, for which the most significantly enriched GO terms were related to response to chemicals/toxins (Extended Data Table 2). Of the 21 TFs that are associated with all six BGCs, none interact with any of the characterized genes for the biosynthetic pathway of the benzoxazinoids^11^ (e.g., DIBOA, DIMBOA), a group of well-characterized defense compounds found in several cereal crops, including wheat^12^ (Extended Data Table 2). This is consistent with the definition of benzoxazinoids as phytoanticipins (constitutively produced defense compounds)^13^, also reflected by the fact that the benzoxazinoid biosynthetic genes are not found in the pathogen-induced WGCNA expression modules.

### The six predicted biosynthetic gene clusters comprise co-expressed genes potentially involved in diterpene, triterpene and flavonoid metabolism

Plant BGCs typically contain one or more genes required for generation of a natural product scaffold, along with genes encoding downstream tailoring enzymes that modify this scaffold (e.g. cytochromes P450 (CYPs), sugar transferases (UGTs), methyl transferases (MTs))^14^. The six predicted pathogen-induced wheat BGCs each contain 5-7 co-expressed biosynthetic genes (Fig.1c, Supplementary Fig. 1). Based on the gene annotations, the predicted scaffold-forming enzymes for the clusters are terpene synthases (TPSs) (clusters 1(2A), 1(2D) and 2(2B)); oxidosqualene cyclases (OSCs) (clusters 3(5A) and 3(5D)); and a chalcone synthase (CHS) (cluster 4(5D)), hallmarks of diterpene, triterpene and flavonoid biosynthesis, respectively. Notably, all three classes of compound are associated with plant defense, including in the grasses^15–18^.

Co-expression within each cluster was assessed by calculation of the Pearson correlation coefficient (r-val) between the expression of a representative scaffold-forming gene from each cluster and other cluster genes, within an RNA-seq dataset including 68 experiments from the wheat-expression.com website^8,19^. In the putative diterpene clusters 1(2A) and 1(2D), several genes were found to be highly co-expressed with the TPS bait (r-val>0.8), including a copalyl diphosphate synthase (CPS), encoding a key enzyme in diterpene biosynthesis that typically catalyzes the preceding step to TPS; one 1(2D) or two 1(2A) UGTs; and three CYPs. In cluster 2(2B), two TPSs, two CYPs and a CPS are co-expressed. In clusters 3(5A) and 3(5D) all five genes are co-expressed, while in cluster 4(5D) all genes are co-expressed with the exception of one chalcone synthase duplicate and a chalcone-flavanone isomerase (Supplementary Table 1, Fig. 1c).

### The type 4 biosynthetic gene cluster 4(5D) encodes a functional flavonoid biosynthetic pathway

To establish whether the predicted BGCs were likely to be functional, we first investigated the candidate flavonoid BGC 4(5D) (Fig. 1c, Supplementary Table 1, Supplementary Fig. 2). The genes for the predicted scaffold-generating enzyme (TaCHS1) and co-expressed tailoring enzymes (TaCYP71C164 and TaOMT3/6/8) were cloned and transiently expressed in *Nicotiana benthamiana* by agroinfiltration^20^, together with the clustered chalcone-flavanone isomerase (chi-D1), and an additional CYP71 gene (TaCYP71F53_5D), which is located 425 Kb upstream of the terminal OMT of the cluster and also belongs to the ME25 expression module. A fourth OMT in the cluster (TaOMT7) is a tandem duplicate of TaOMT6 with a single amino acid difference and was not included in the analysis. The combined expression of all genes resulted in formation of a new product exhibiting UV absorbance (λmax = 260nm) with exact mass [M+H=329.1010 (Supplementary Fig. 2) and predicted elemental composition C_18_H_17_O_6_ (−1.78 ppm). This product was not produced in combinations in which TaCHS1 or any of the two CYPs and three OMTs were omitted, indicating that the proteins encoded by all six genes are enzymatically active (Fig. 1e, Supplementary Fig. 2). Inclusion of chi-D1 was not essential for formation of this product in *N. benthamiana* (Supplementary Fig. 2). Thus, the co-expressed genes within cluster 4(5D) encode a functional pathway that, based on UV absorbance, exact mass and the calculated elemental composition of the putative end-product, is likely to produce a hydroxy-trimethoxy-flavone. Future work is needed to fully elucidate the structures of the pathway end-product and intermediates.

### The homoeologous type 1 biosynthetic gene clusters 1(2A) and 1(2D) are related to but functionally distinct from the rice momilactone cluster

Rice produces a variety of diterpene phytoalexins for which the biosynthetic pathways are well-characterized. The genes for several of the pathways for labdane-related diterpenes (e.g. momilactones, phytocassanes/oryzalides) are clustered in the rice genome^21–23^. These labdane-related diterpenes are formed from the universal diterpenoid precursor geranylgeranyl diphosphate (GGPP) (**1**) via initial cyclization reactions catalyzed by copalyl diphosphate synthases (CPSs) that produce normal, *ent*, or *syn* stereoisomers of copalyl diphosphate (CPP). The CPP intermediates are subsequently utilized by terpene synthases (TPSs) to form various diterpene backbones, which then typically undergo further tailoring reactions^24^. In wheat, diterpene metabolism is considerably less well characterized than in rice. In previous studies aimed at functional characterization of diterpene-related genes in wheat, five copalyl diphosphate synthases (CPS1-5) and six kaurene synthase-like terpene synthases (KSL1-6) were cloned and characterized by recombinant expression^25–27^. Four of the CPS enzymes catalyzed production of normal or *ent* stereoisomers of CPP, while five of the KSL enzymes were shown to convert normal-, *ent*-, or *syn*-CPP to several different diterpene products. The physical location and general expression patterns of these genes were, however, unknown.

Interestingly, our study identified two of these genes, namely *TaKSL1* and *TaCPS2*, as the co-expressed TPS and CPS genes in cluster 1(2A). A third gene, *TaKSL4* is found in the homoeologous 1(2D) cluster (Fig. 2a). *TaKSL4* is not co-expressed with other cluster genes and generally exhibits a root-specific, non-induced expression pattern (Supplementary Fig. 1). The co-localization and co-expression of *TaKSL1* and *TaCPS2* coincides with their previously ascribed enzymatic functions - TaCPS2 produces normal-CPP (**2**), while TaKSL1 acts on normal-CPP to produce isopimara-7,15-diene (**3**)^25,26^. TaKSL1 can also react with a *syn*-CPP substrate, but a *syn*-CPP producing copalyl synthase is yet to be identified in wheat^26^. Transient expression of the Chr.2D homoeologs of TaCPS2 and TaKSL1 (named TaCPS-D2 and TaKSL-D1 hereinafter) in *N. benthamiana* revealed that these enzymes are functional and produce compounds with mass spectra matching copalol and isopimara-7,15-diene respectively, confirming the activity of this pair of genes in the 1(2D) cluster (Fig 2b, Supplementary Fig. 3). The occurrence of additional co-expressed CYP genes and a UGT gene in the 1(2D) and 1(2A) clusters (Fig. 2a) suggests that these clusters form pathogen-induced pathways for production of isopimara-7,15-diene-derived diterpenes (Fig 2c).

**Fig. 2:**
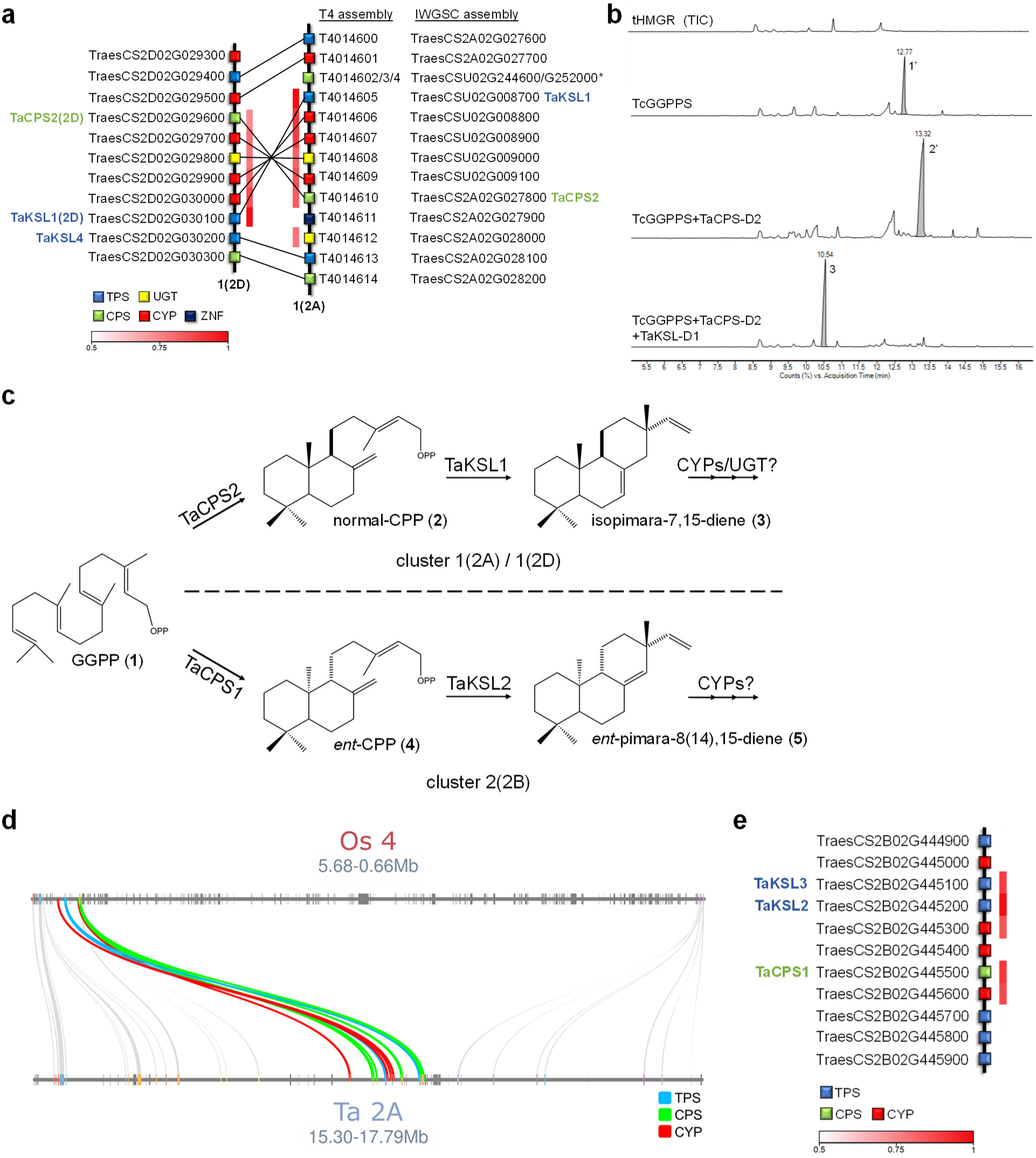
Diterpene-producing BGCs are found on group 2 chromosomes in bread wheat. **a**, assignment of homoeologous genes in the type 1 clusters 1(2A) and 1(2D), including the previously characterized genes *TaCPS2*, *TaKSL1* and *TaKSL4*. Chr.2A genes were positioned based on the T4 wheat genome assembly^59^ and homoeologs were assigned based on pairwise sequence alignments. The T4 assembly reveals the presence of five Chr.2D homoeologs in inverted positions on Chr.2A (*TraesCSU02G008700-G009100*), which were previously unmapped in the IWGSC assembly. *CPS* genes *TraesCSU02G252000* and *TraesCSU02G244600* (asterisked) have partial coding sequences. *TPS*, terpene synthase; *UGT*, UDP-dependent glycosyltransferase; *CPS*, copalyl diphosphate synthase; *CYP*, cytochrome P450; *ZNF*, Zinc finger, RING/FYVE/PHD-type. The white to red color-coding denotes Pearson correlation (r) values for expression of each gene with a representative ‘bait’ gene from the cluster. **b**, GC-MS analysis of leaf extracts following expression of the wheat TaKSL-D1 and TaCPS-D2 enzymes in *N. benthamiana.* Cytosol-targeted TaKSL-D1 and TaCPS-D2 were transiently expressed together with a *Taxus canadensis* GGPP synthase and oat tHMGR. Total ion chromatograms (TIC) are shown. Peaks were putatively identified as geranylgeraniol (**1’**), copalol (**2’**), and isopimara-7,15-diene (**3**), based on comparison of mass spectra to the NIST database and the literature (see Supplementary Fig. 3). **c**, predicted pathways for diterpene production by the type 1 BGCs 1(2A) and 1(2D), and the type 2 BGC 2(2B). The type 1 clusters 1(2A) and 1(2D) comprise co-expressed genes for TaCPS2 and TaKSL1, CYPs and UGTs, predicted to form isopimara-7,15-diene-derived diterpenoids from geranylgeranyl diphosphate (GGPP). Cluster 2(2B), includes co-expressed genes for TaCPS1, TaKSL2, TaKSL3 and two CYPs, putatively forming pimara-8(14),15-diene-derived diterpenoids from GGPP. **d**, microsynteny analysis of wheat BGC 1(2A) (T4 assembly) and the momilactone cluster in a syntenic region in rice Chr.4. **e**, structure of BGC 2(2B) and assignment of the previously characterized genes *TaCPS1*, *TaKSL2* and *TaKSL3*.

Intriguingly, microsynteny analysis between wheat and rice suggests that the type 1 clusters present on wheat chromosomes 2A and 2D (BGCs 1(2A) and 1(2D)) likely share a common evolutionary origin with the rice momilactone cluster. The *KSL* genes in clusters 1(2A) and 1(2D) are close homologs of the *OsKS4* gene from the rice momilactone BGC^21,22^. Directly adjacent to *TaKSL*1 is a *CPS* gene that is orthologous to *OsCPS4*. The wheat cluster also includes four cytochrome P450s belonging to the CYP99 family that are homologs of the CYP99A2/A3 P450 pair in the rice momilactone BGC^22^. Furthermore, the chromosomal regions harbouring wheat clusters 1(2a) and 1(2D) are syntenic to the region of the rice genome containing the momilactone cluster, which is found on rice Chr.4, the corresponding chromosome of wheat Chr.2^28^ (Fig. 2d). However, although these clusters may share a common evolutionary origin, they produce different types of diterpenes: the rice momilactones are derivatives of the *syn*-CPP-derived scaffold *syn*-pimara-7,15-diene^21^, while functional characterization and gene expression data of the wheat 1(2D) cluster and previous characterization of the *TaKSL1* and *TaCPS2* genes^25,26^, which we have shown to be in wheat cluster 1(2A), implies that these two BGCs encode pathways that yield derivatives of the normal-CPP-derived isopimara-7,15-diene-scaffold. Of note, the rice momilactone cluster also includes two short-chain dehydrogenase/reductase (SDR) genes *OsMAS* and *OsMAS2*^22,29^ that do not have apparent orthologs in the wheat type 1 clusters or elsewhere in the wheat genome.

The third predicted diterpene BGC that we found, cluster 2(2B), also includes three other previously characterized wheat genes, namely *TaCPS1*, *TaKSL2* and *TaKSL3*^25,26^, all of which are co-expressed (Fig. 2e). TaCPS1 catalyzes formation of *ent*-CPP (**4**), while TaKSL2 acts on *ent*-CPP to produce pimara-8(14),15-diene (**5**). TaKSL3, a tandem duplicate of TaKSL2, only exhibits low activity, selectively acting on *ent*-CPP to produce two unknown products^26^. The combined functions of TaCPS1 and TaKSL2, together with the presence of additional co-expressed CYPs in the cluster, suggest that BGC 2(2B) encodes a pathway for production of *ent*-pimara-8(14),15-diene derivatives (Fig. 2c).

### The homoeologous type 3 cluster 3(5D) encodes a biosynthetic pathway to ellarinacin, a novel arborinane-type triterpenoid

The type 3 cluster 3(5D) contains genes implicated in triterpenoid biosynthesis, most notably a predicted oxidosqualene cyclase gene (*TaOSC*). Flanking *TaOSC* are three cytochrome P450s (*TaCYP51H35, TaCYP51H37* and *TaCYP51H13P*) and a gene annotated as a 3β-hydroxysteroid-dehydrogenase/decarboxylase (*TaHSD*) (Fig. 3a). The genomic sequences of *TaOSC*, *TaHSD*, *TaCYP51H35* and *TaCYP51H37* predict full coding sequences for all four genes, while *TaCYP51H13P* was found by manual annotation to carry two premature stop codons (Supplementary Fig. 4) and was designated a pseudogene. The homoeologous cluster on Chr.5A is similarly structured (Fig. 3a), but with a predicted full coding sequence for *TaCYP51H13_5A*. Amino acid sequence identity between homoeologous pairs in the 3(5D) and 3(5A) clusters is >99% for TaOSC, TaHSD and TaCYP51H37, and >97% for TaCYP51H35 and TaCYP51H13. As for the type 1 diterpene cluster, homoeologs of the type 3 cluster genes are not found in the B genome. However, a homologous gene cluster is present adjacent to the 3(5D) cluster on Chr.5D, which includes paralogs of the *TaOSC*, *TaHSD*, and *TaCYP51H* genes. Similarly, one *TaOSC* and two *CYP51H* paralogs are also found on Chr.5A, adjacent to the 3(5A) cluster (Supplementary Fig. 5). These Chr.5A and Chr.5D paralogs, however, in general have low expression across all transcriptomic data available on wheat-expression.com, and so are not likely to belong to active BGCs (Supplementary Table 2).

**Fig. 3:**
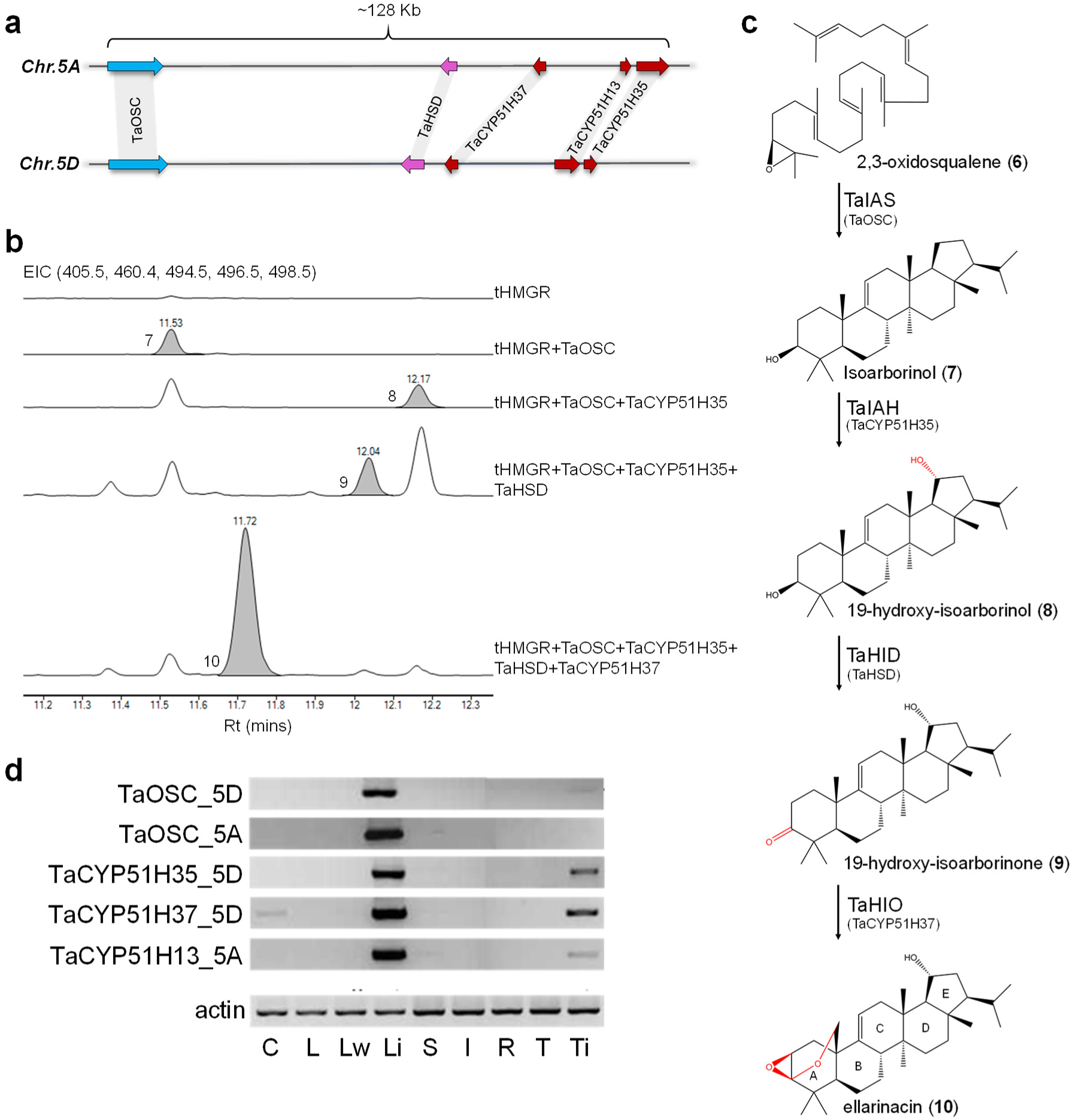
Wheat cluster 3(5D) produces an isoarborinol-derived triterpenoid. **a**, structures of homoeologous triterpene biosynthetic gene clusters identified on wheat chromosomes 5A and 5D. **b**, GC-MS traces for wheat BGC 3(5D) genes transiently expressed in *N. benthamiana*. EIC, extracted ion chromatogram for ions representing isoarborinol (**7**) (498.5), 19-hydroxy-isoarborinol (**8**) (496.5), 19-hydroxy-isoarborinone (**9**) (494.5), ellarinacin (**10**) (405.5) and internal standard 5α-cholestan-3β-ol (460.4). **c**, assigned structure of ellarinacin and predicted biosynthetic pathway in wheat. TaIAS, isoarborinol synthase; TaIAH, isoarborinol 19-hydroxylase; TaHID, 19-hydroxy-isoarborinol dehydrogenase; TaHIO, 19-hydroxy-isoarborinone oxidase. Rings A-E are annotated. **d**, semiquantitative RT-PCR of selected genes from type 3 clusters 3(5A) and 3(5D) in ‘Chinese Spring’ wheat tissues. C, coleoptile; L, leaf; L_w_, leaf after wounding; L_i_, leaf infected with *Blumeria graminis f. sp. tritici*; S, stem; I, inflorescence; R, root; T, root tip; T_i_, root tip after infection with *Gaeumannomyces graminis*.

Functional analysis of the cluster 3(5D) genes was carried out by transient expression in *N. benthamiana*. All genes were co-infiltrated with an *Agrobacterium* strain harbouring an expression construct for a feedback insensitive form of 3-hydroxy-3-methylglutaryl coenzyme A reductase (tHMGR) from oat, which enhances triterpenoid precursor supply^30^. GC- and LC-MS analyses of leaf extracts revealed that the four enzymes TaOSC, TaHSD, TaCYP51H35 and TaCYP51H37 form a sequential biosynthetic pathway (Fig. 3b, Supplementary Figs. 6-9). As the *TaCYP51H13P* pseudogene from cluster 3(5D) does not encode a complete functional protein, we tested the activity of its Chr.5A homoeolog, *TaCYP51H13_5A*, through agroinfiltration with the four 3(5D) cluster genes in different combinations. TaCYP51H13_5A exhibited the same activity as TaCYP51H35, but to a lower extent, resulting in lower levels of product compared to TaCYP51H35 (Supplementary Fig. 10). This redundant activity provides a possible explanation why *TaCYP51H13* is not conserved in the Chr.5D cluster.

The structures of the purified products of co-expression of TaOSC+TaCYP51H35, and of the combined four cluster genes (all from cluster 3(5D)) were determined by NMR following large-scale vacuum-mediated agroinfiltration and purification (Supplementary Figs. 11,12, Supplementary Tables 3, 4). The product of co-expression of TaOSC+TaCYP51H35 was identified as 19-hydroxy-isoarborinol (**8**), indicating that TaOSC (hereinafter isoarborinol synthase, TaIAS) synthesizes the triterpene scaffold isoarborinol (**7**) which is subsequently hydroxylated by TaCYP51H35 (hereinafter isoarborinol 19-hydroxylase, TaIAH). The product of co-expression of all four cluster genes was found to have an unusual triterpenoid structure, with a β-epoxy group and an ether bridge attached to the A ring (Fig. 3c). The GC/LC-MS data and NMR-assigned structure together suggest oxidation of the 3-alcohol to the ketone 19-hydroxy-isoarborinone (**9**) by TaHSD (hereinafter 19-hydroxy-isoarborinol dehydrogenase, TaHID); TaCYP51H37 (hereinafter 19-hydroxy-isoarborinone oxidase, TaHIO) likely then hydroxylates the C25-methyl carbon, leading to nucleophilic attack of the A-ring ketone, thus forming a hemiacetal intermediate which further reacts to produce the epoxide. This unusual reaction may involve two independent catalytical cycles mediated by TaHIO. This would be in line with the only other previously reported non-canonical CYP51 enzyme (AsCYP51H10*, Sad2*) which hydroxylates the C16 position of the β-amyrin scaffold and also converts an alkene to an epoxide at C12-C13 via two independent reactions^31^. However, a mechanism involving just one catalytic cycle may also be possible (Supplementary Fig. 13). The structure of the BGC 3(5D) product has not, to the best of our knowledge, been previously reported, and was named ellarinacin (**10**). The proposed biosynthetic pathway is shown in Fig. 3c.

Interestingly, the ellarinacin cluster (BGC 3(5D)) provides multiple links to sterol metabolism. Production of plant sterols from 2,3-oxidosqualene (**6**) is initiated by highly conserved OSCs known as cycloartenol synthases (CASs), while triterpene scaffolds are generated from 2,3-oxidosqualene by other diverse OSCs (triterpene synthases)^16^. TaOSC shares higher sequence similarity with characterized monocot CAS enzymes in comparison to other functionally characterized monocot triterpene synthases (Supplementary Figs. 14, 15). Plant 3β-hydroxysteroid-dehydrogenase/decarboxylases belong to the short chain dehydrogenase reductase (SDR) superfamily and are involved in biosynthesis of phytosterols and steroidal glycoalkaloids^32–35^. Phylogenetic analysis shows that TaHSD is related to *Arabidopsis thaliana* genes *3*β*HSD/D1* and *3*β*HSD/D2*, that take part in sterol biosynthesis (Supplementary Fig. 16)^32,36^. The cytochrome P450 genes found in the cluster provide further connections to sterol metabolism, as they belong to the sterol-related CYP51 family (Supplementary Figs. 17,18). CYP51 enzymes catalyze 14α-demethylation of sterols in all eukaryotes and are the only family of cytochrome P450s that are evolutionarily conserved from prokaryotes through fungi, plants, and mammals^37^. To date, only one plant CYP51 has been found to catalyze a reaction different from the canonical sterol demethylase activity- AsCYP51H10 (*Sad2)*, which is involved in biosynthesis of an antifungal triterpene glycoside known as avenacin in oat^31,38^. Several members of the ellarinacin cluster thus appear to have been recruited from sterol biosynthetic genes, most likely through gene duplication and neofunctionalization.

### The ellarinacin cluster is highly induced by biotic stress

We next sought to determine whether the type 3 clusters are likely to be involved in plant defense by further investigating the expression patterns of the clustered genes. Analysis of the wheat-expression.com dataset revealed that the expression patterns of the genes were consistent with their positioning in the ME25 and ME34 modules, i.e. that they showed induction by various fungal pathogens and by the PAMPs chitin and flg22. (Supplementary Fig. 19). Notably, the clusters were not substantially induced in response to various abiotic stresses, including drought, heat, cold, phosphate starvation and drought-simulating treatment with PEG-6000 (Supplementary Fig. 20).

The observed expression pattern of the type 3 clusters was further supported by semiquantitative RT-PCR analysis of selected genes from BGC 3(5A) and 3(5D) using homoeolog-specific primers: expression of all tested genes was strongly induced in leaves infected with powdery mildew (Li) but not by mechanical wounding (Lw), with little or no expression in the other various wheat tissues analyzed (Fig. 3d). Weak induction was also observed in roots infected with *Gaeumannomyces graminis*, a soil-borne fungus that causes ‘take-all’ disease (Ti). Induction of the entire BGC 3(5D) by infection with powdery mildew was further validated by quantitative real-time PCR (qRT-PCR). Detached wheat leaves were exposed to spores of either wheat-adapted (*Blumeria graminis f. sp. tritici*, Bgt) or non-adapted (*Blumeria graminis f. sp. hordei,* Bgh) isolates of powdery mildew, and relative transcript abundance was determined 12 and 24 hours post infection. Treatment with Bgt or Bgh resulted, in both cases, in strong induction of the four cluster genes. Interestingly, induction was more marked for Bgh (non-adapted) compared to Bgt (Fig. 4a). Our analyses of transcriptome data from previously published studies^39,40^ in which wheat plants were challenged with the fungal pathogens powdery mildew, cereal blast (*Magnaporthe* spp.), and leaf or yellow rust (*Puccinia* spp.) also revealed stronger induction of the cluster genes by non-host vs. host interactions (Supplementary Figs. 21, 22).

**Fig. 4:**
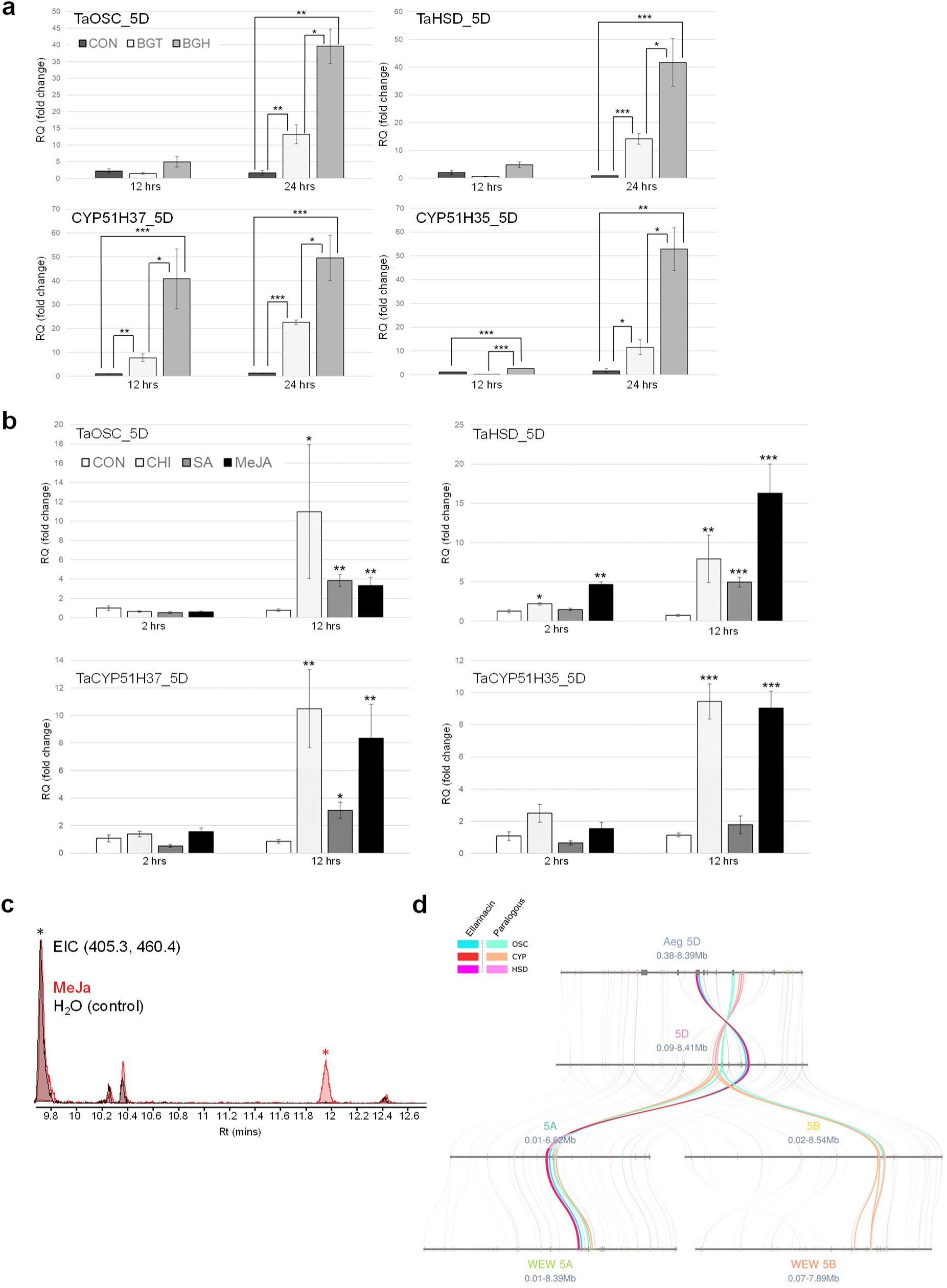
The ellarinacin BGC 3(5D) is induced by pathogens and elicitors. **a**, quantitative real-time PCR (qRT-PCR) of ellarinacin BGC genes in detached wheat leaves infected with two powdery mildew isolates, 12 and 24 hours post infection. Con, control (non-infected); Bgt and Bgh, infected with wheat-adapted isolate *Blumeria graminis f. sp. tritici* or the non-adapted isolate *Blumeria graminis f. sp. Hordei*, respectively **b**, quantitative real-time PCR (qRT-PCR) for ellarinacin BGC genes in detached wheat leaves treated with methyl jasmonate (MeJa), salicylic acid (SA), chitin (CHI) or H_2_O (CON), for 2 or 12 hours. For panels A and B, relative quantification values (in fold-change) indicate means of three biological replicates ± SEM. Asterisks denote t-test statistical significance of differential expression. *, *p*-val<0.05. **, *p*-val<0.01. ***, *p*-val<0.001. **c**, GC-MS analysis of TMS-derivatized extracts from wheat leaves treated with methyl jasmonate (MeJa), or H_2_O (control) for three days. Extracted ion chromatograms are for ions representing ellarinacin (405.3, Rt 11.94, red asterisk) and 5α-cholestan-3β-ol (460.4, Rt 9.70, black asterisk). **d**, microsynteny analysis of the region surrounding the ellarinacin BGC and its paralogous cluster in Chr.5 of the wheat A, B and D genomes, and wheat progenitors *Aegilops tauschii* (Aeg) and wild emmer wheat (WEW).

Finally, we analyzed gene expression in detached wheat leaves treated with the elicitors methyl jasmonate (MeJa) and salicylic acid (SA), as well with the PAMP, chitin. All four cluster 3(5D) genes analyzed (*TaOSC*, *TaHSD*, *TaCYP51H35*, *TaCYP51H37*) were significantly induced compared to the control 12 hrs after treatment with MeJa, SA or chitin, with the exception of *TaCYP51H35* in SA-treated leaves (Fig. 4b). Thus, the ellarinacin cluster is highly induced by biotic stress, suggesting a possible function in wheat response against pathogens. The very low basal expression in various wheat tissues, as observed in the RT-PCR and RNA-seq data analysis, and strong induction by pathogens, defense-related hormones and PAMPs, further suggests that ellarinacin serves as a phytoalexin rather than a phytoanticipin. Correspondingly, GC-MS analysis detected ellarinacin in extracts of MeJa-treated but not control detached wheat leaves (Fig. 4c). A 60% increase in isoarborinol levels was also observed in MeJa-treated leaves compared to control leaves (Supplementary Fig. 23).

### The ellarinacin cluster is conserved in wheat ancestors

We next investigated whether ellarinacin-like clusters also exist in the genomes of ancestral species of common wheat. Specifically, we looked for related clusters in two wild progenitors that have sequenced genomes; *Aegilops tauschii* (Tausch’s goatgrass; donor of the D genome of bread wheat), and *Triticum turgidum ssp. diccocoides* (wild emmer wheat, progenitor of cultivated emmer; the donor of the A and B genomes of bread wheat). Microsynteny analysis of the regions surrounding the homoeologous type 3 BGCs on Chr.5A and Chr.5D show that while these clusters appear to be conserved on chromosome 5 of the A and D genomes of *A. tauschii* and wild emmer wheat, a homoeologous cluster could not be found on chromosome 5B of bread wheat or wild emmer wheat. Chromosome 5B of both species do, however, contain homoeologs of the OSC and/or P450s of the paralogous, transcriptionally non-active cluster in the A and D genomes (Fig. 4d). The wild emmer wheat and *A. tauschii* clusters each contain an OSC, an HSD and three CYP51 genes, in the same order and orientation as in wheat (Fig. 5a). Sequence comparison of the cluster genes in wheat and its two wild progenitors revealed that the predicted protein sequences are also highly conserved (>99.4% amino acid identity for all proteins in both species; Supplementary Table 5). To assess the functionality of the *A. tauschii* cluster, we transiently expressed the first two genes of the predicted *A. tauschii* pathway, namely the orthologs of *TaIAS* and *TaIAH*, in *N. benthamiana*. Co-expression of the two genes resulted in formation of 19-hydroxy-isoarborinol (Supplementary Fig. 24), the same product obtained by *TaIAS* and *TaIAH* expression. The coding sequence of the *A. tauschii* ortholog of *TaCYP51H13P* contains one of the premature stop codons found in its wheat homolog (Supplementary Fig. 4), and so is likely to be non-functional. The remaining predicted active enzymes in the *A. tauschii* pathway, orthologs of TaHID and TaHIO, exhibit 100% amino acid identity with their wheat counterparts, suggesting that the *A. tauschii* cluster encodes a complete biosynthetic pathway for ellarinacin.

**Fig. 5:**
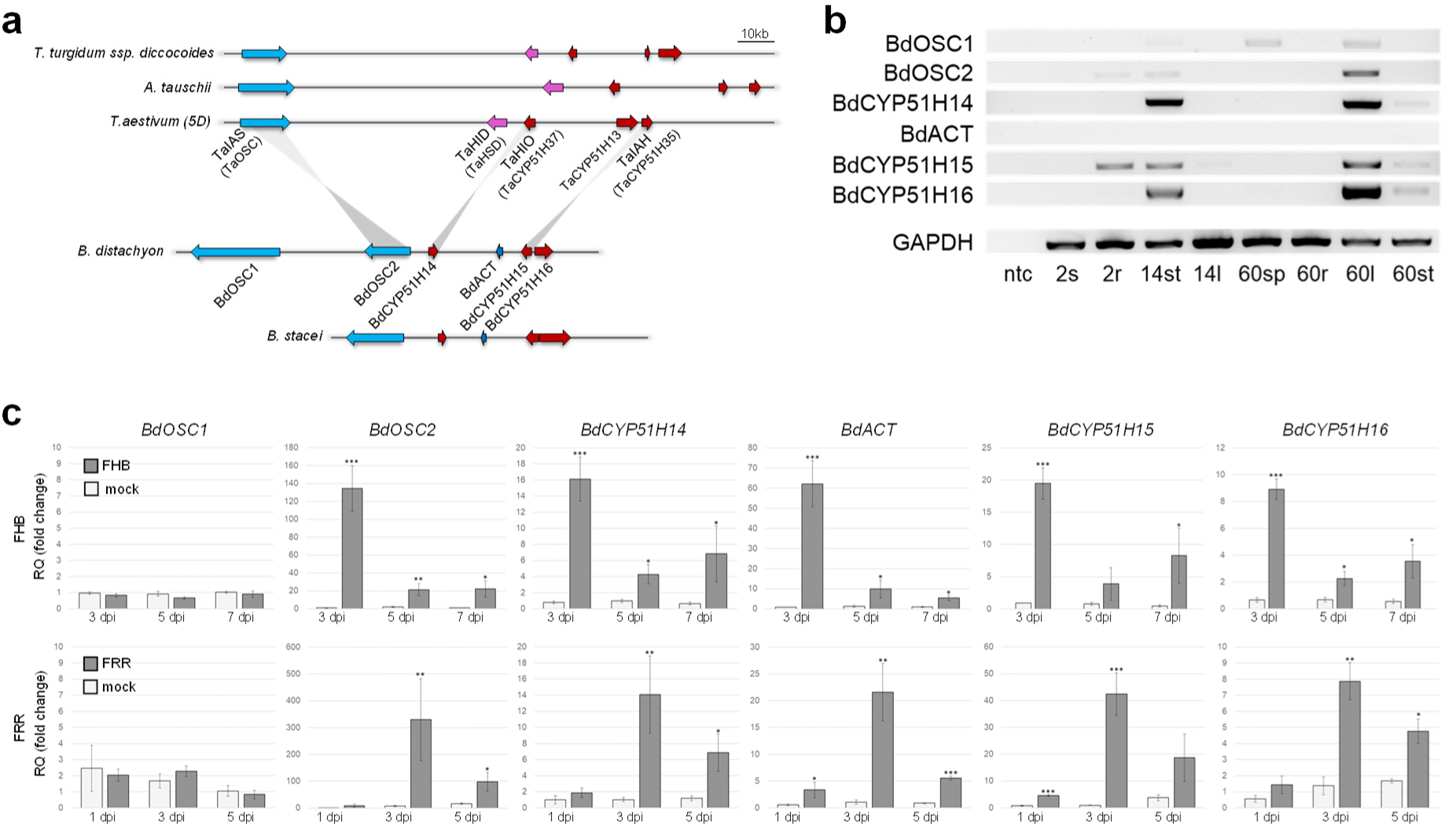
Occurrence and expression of ellarinacin-like BGCs in *Brachypodium* and wheat ancestral species. **a**, wheat ellarinacin BGC is conserved in wheat wild ancestors *Aegilops tauschii* and wild emmer wheat (*Triticum turgidum subsp. dicoccoides*), and homologous to a BGC identified in chromosome 3 of *Brachypodium distachyon*. Grey lines link between wheat and *B. distachyon* BlastP reciprocal best hits. **b**, semiquantitative RT-PCR of *B. distachyon* Chr.3 clustered genes. ntc, no template control; 2s, seedling shoot (2 day old); 2r, seedling root; 14st, young plant stem base (14 day old); 14l, young plant leaf; 60sp, mature plant spike (60 day old); 60r, mature plant root; 60l, mature plant leaf; 60st, mature plant stem base. **c**, quantitative real-time PCR (qRT-PCR) of brachynacin BGC genes in *B. distachyon* plants infected with Fusarium head blight (FHB) or Fusarium root rot (FRR). Con, control (non-infected); dpi, days post infection. Relative quantification values (in fold-change) indicate means of three biological replicates ± SEM. Asterisks denote t-test statistical significance of differential expression. *, *p*-val<0.05. **, *p*-val<0.01. ***, *p*-val<0.001.

### Arborinane-type clusters are found in other grasses

The occurrence of conserved ellarinacin-like clusters in wheat and its progenitors raised the possibility that BGCs for ellarinacin or other arborinane-type terpenoids may also occur in other grasses. The isoarborinol scaffold has been reported from other Poaceae species, including sorghum^41^ and rice^42^. We therefore searched for orthologs of *TaIAS* in additional Poaceae species, based on sequence similarity. Orthologs for *TaIAS* could not be identified in maize, sorghum, barley, and rice. The latter has a previously characterized isoarborinol synthase gene^42^, but this gene bears low similarity to *TaIAS* (56% similarity on amino acid level) and has most likely evolved independently.

A BlastP search of TaIAS against the recently published genome of the diploid oat species *Avena strigosa*^43^ found a candidate OSC gene on chromosome 1, herein named *AsOSC1*, with high predicted amino acid sequence similarity to the TaIAS protein (91.2%). This was also the reciprocal best hit (RBH) of TaIAS. Flanking *AsOSC*1 (∼25 Kb away) is a *CYP51H* gene, herein named *CYP51H73*. Transient expression of AsOSC1 in *N. benthamiana* yielded a new product, which was verified by GC-MS as isoarborinol. No additional products were detected when AsCYP51H73 was co-expressed with AsOSC1 (Supplementary Fig. 25). Since *AsCYP51H73* is orthologous with wheat *TaCYP51H37* (*TaHIO*), we also tested if AsCYP51H73 would exhibit the same or similar activity, by co-expressing AsCYP51H73 together with TaIAS, TaIAH and TaHID. However, no activity was detected.

In the genome of the grass model plant *Brachypodium distachyon* (strain *Bd21*)^44^, a *TaIAS* homolog was identified on chromosome 3, *BdOSC2*, which was the RBH of *TaIAS*. Flanking this gene were genes predicted to encode another highly similar OSC (*BdOSC1*), and three cytochrome P450s of the CYP51H subfamily (*BdCYP51H14*, *BdCYP51H15* and *BdCYP51H16*) (Fig. 5a). A predicted BAHD-type acyltransferase gene (*BdACT*) was also found between *BdCYP51H14* and *BdCYP51H15*. Thus, together these genes form a potential BGC for production of arborinane-type or similar triterpenoids in *B. distachyon*. A conserved cluster that has a similar gene structure to the *B. distachyon* BGC but with one OSC gene only was also found in the genome of the closely related species *B. stacei* (Fig. 5a).

### The *B. distachyon* Chr.3 BGC is induced by fungal pathogens

To test whether the clustered genes identified on chromosome 3 of *B. distachyon* might form an active BGC, their expression profiles were examined. Analysis of *B. distachyon* gene expression datasets in JGI Gene Atlas (https://phytozome.jgi.doe.gov/),^45^) and PlaNET (http://aranet.mpimp-golm.mpg.de/)^46,47^, showed that the three *CYP51* and two *OSC* genes are co-expressed, with highest expression in the mature leaf and stem base. The BAHD acyltransferase gene displayed a similar pattern, but with markedly lower overall expression values (Supplementary Fig. 26). Relative expression of all six genes was further assessed by semi-quantitative RT-PCR of seven *B. distachyon* tissues at different developmental stages. The cluster genes generally exhibited highest expression in the leaves and stem base (Fig. 5b). A *BdACT* amplicon could only be detected with extended exposure (Supplementary Fig. 27). Unlike the other cluster genes, *BdOSC1* was also expressed in the spikes of mature plants.

qRT-PCR analysis of *B. distachyon* plants infected with *Fusarium graminearum* causing Fusarium head blight or Fusarium root rot showed that, as for the wheat cluster, the *B. distachyon* cluster is highly induced by fungal pathogens. Significant increases in gene expression following infection were observed in both experiments for all clustered genes except *BdOSC1* (Fig. 5c).

### The *B. distachyon* Chr.3 BGC produces an arborinane-type triterpenoid

Since gene expression analysis suggested an active BGC, we next investigated the functions of the cluster genes by transient expression in *N. benthamiana* (Fig. 6a). GC-MS analysis revealed that BdOSC2 and CYP51H15 exhibit the same activities as their respective wheat orthologs, i.e., production of isoarborinol and its 19-hydroxylated derivative (Supplementary Fig. 28). Co-expression of BdOSC2 and CYP51H15 together with the two additional CYP51s and the BdACT acyltransferase resulted in formation of the putative BGC end product, with a mass signal of [M+H-H_2_O=515.3] (Supplementary Figs. 29, 30). This product was purified following large-scale transient expression of the *B. distachyon* cluster genes in *N. benthamiana* and found by ^1^H and ^13^C NMR analyses to be an isoarborinol-derived triterpenoid with hydroxyl groups on the C7,19,28 carbons and an acetoxy group on the C1 carbon. (Supplementary Fig. 31, Supplementary Table 6). The assigned structure allowed the full elucidation of the biosynthetic pathway from 2,3 oxidosqualene, in which BdOSC2 and BdCYP51H15 generate 19-OH-isoarborinol, BdCYP51H14 hydroxylates the C7 and C28 carbons to give 7,19,28-trihydroxy-isoarborinol (**11**), and BdCYP51H16 hydroxylates the C1 carbon to give 1,7,19,28-tetrahydroxy-isoarborinol (**12**), which is further acetylated by BdACT (Fig. 6b). This compound, which has not previously been reported, was named brachynacin (**13**). The occurrence of brachynacin in *B. distachyon* was verified by GC-MS analysis of leaf extracts. As for ellarinacin in wheat, the relative abundance of brachynacin, as well as of isoarborinol, were found to be significantly higher in MeJa-treated vs. non-treated detached leaves (Fig. 6c, Supplementary Fig. 32).

**Fig. 6:**
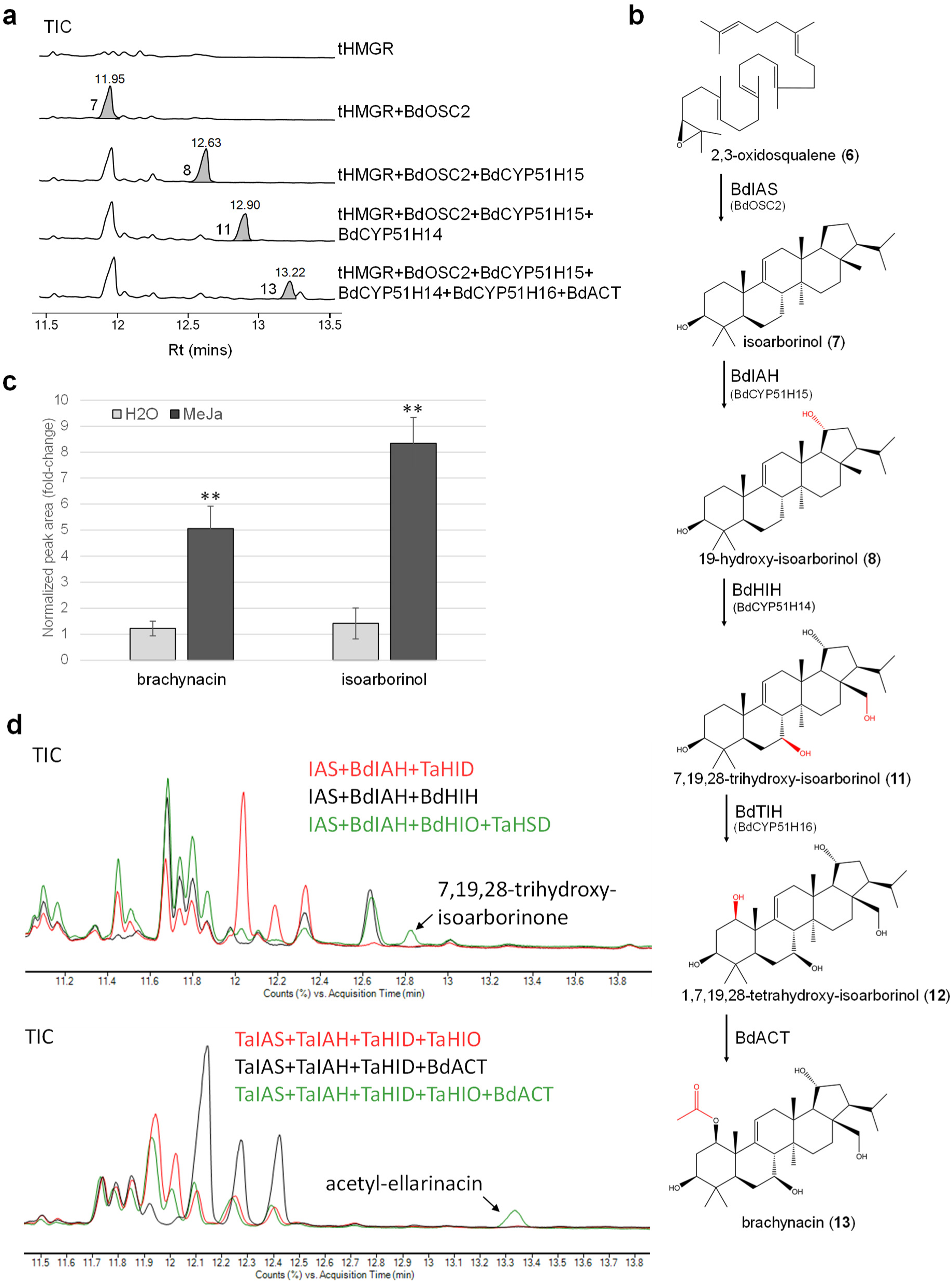
*B. distachyon* BGC produces the isoarborinol-derived triterpenoid, brachynacin. **a,** GC-MS traces for *B. distachyon* cluster genes transiently expressed in *N. benthamiana*. TIC, total ion chromatogram. Marked peaks were identified as isoarborinol (**7**), 19-hydroxy-isoarborinol (**8**), 7,19,28-trihydroxy-isoarborinol (**11**) and brachynacin (**13**) (494.5). **b**, assigned structures and predicted biosynthetic pathway of brachynacin in *B. distachyon*. BdIAS, isoarborinol synthase; BdIAH, isoarborinol hydroxylase; BdHIH, 19-hydroxy-isoarborinol hydroxylase; BdTIH, 7,19,28-trihydroxy-isoarborinol hydroxylase. BdACT 1,7,19,28-tetrahydroxy-isoarborinol acetyltransferase. **c**, relative abundance of isoarborinol and brachynacin in TMS-derivatized extracts of *B. distachyon* leaves treated with MeJa or H_2_O for 12 hours. Relative quantification is based on normalized peak areas in GC-MS analysis of four biological replicates. **d**, GC-MS total ion chromatograms (TIC) of *N. benthamiana* leaves transiently expressing combinations of wheat and *B. distachyon* genes.

### Combination of ellarinacin and brachynacin biosynthetic genes yields novel compounds

The similarities between the ellarinacin and brachynacin BGCs indicates that they possibly originate from a common ancestral cluster but have evolved to produce different end products, through duplication and neofunctionalization of CYP51H enzymes and recruitment of additional modifying enzymes (TaHSD and BdACT, respectively). The evolution of these two pathways may have been facilitated by a degree of promiscuity that enabled the pathway enzymes to accept different substrates. Indeed, co-expression of different combinations of genes from the two BGCs in *N. benthamiana* did yield new products. Expression of TaHSD with BdCYP51H14 and BdCYP51H15 led to production of a new compound with a molecular mass [M+H-H_2_O=455.3], matching the expected product, 7,19,28-trihydroxy-isoarborinone (Fig. 6d). Likewise, expression of BdACT with the ellarinacin cluster resulted in formation of a new compound identified as acetyl-ellarinacin, based on its molecular mass signal [M+H=497.3] and the fact that its formation required expression of the entire wheat BGC (Fig. 6d, Supplementary Figs. 33,34). The formation of novel compounds through combining genes from the wheat and *B. distachyon* BGCs demonstrates the promiscuous nature of enzymes encoded by genes within these two clusters.

## DISCUSSION

Despite the importance of wheat as a food and feed crop, our understanding of the molecules that it produces in response to biotic stress remains limited. Conversely, various phytoalexins and their biosynthetic pathways have been well-characterized in other major cereal crops such as rice, maize, oat and sorghum^48,49^, and serve as potential targets for crop improvement^48,50^. Research into specialized metabolism in wheat has until recently been hindered by the lack of a fully assembled genome. The availability of a newly assembled genome coupled with the vast amount of available transcriptomic data now opens up opportunities to deploy genomics-driven approaches for discovery of novel metabolic pathways in wheat, including those implicated in plant defense. Here, utilization of wheat genomic and transcriptomic resources has enabled us to identify previously unknown pathogen-induced biosynthetic pathways for flavonoids, diterpenes and triterpenes. These pathways are driven by sets of genes that are co-localized in the wheat genome, forming six BGCs, including two pairs of homoeologous clusters.

We identified a cluster on wheat Chr.5D, BGC 4(5D), encoding a biosynthetic pathway for O-methylated flavonoids. Although flavonoids form a ubiquitous and highly diverse class of compounds in plants, BGC 4(5D) is the only identified and functionally validated flavonoid BGC to date. Further research will be needed for full structural assignment of the pathway product. Additionally, two different types of diterpene-producing clusters (BGC 1(2A/2D) and BGC 2(2B)), were identified on group-2 chromosomes, one of which is syntenic to the rice momilactone cluster. However, the pathways encoded by these syntenic BGCs diverge in their early steps due to differential CPS activities (i.e., producing *syn*-CPP or normal CPP). Notably, momilactone BGCs were also found in genomes of barnyard grass, *Echinochloa crus-galli*^51^ and the bryophyte *Calohypnum plumiforme* ^52^, the latter having evolved independently from the momilactone cluster in the grasses ^53^. We did not identify a homologous cluster in the other grass genomes that we analyzed, which included *B. distachyon*, oat, barley and maize. A putative terpene cluster homologous to cluster 2(2B) was however found in *B. distachyon* (Fig. 7a, Supplementary Table 7).

**Fig. 7:**
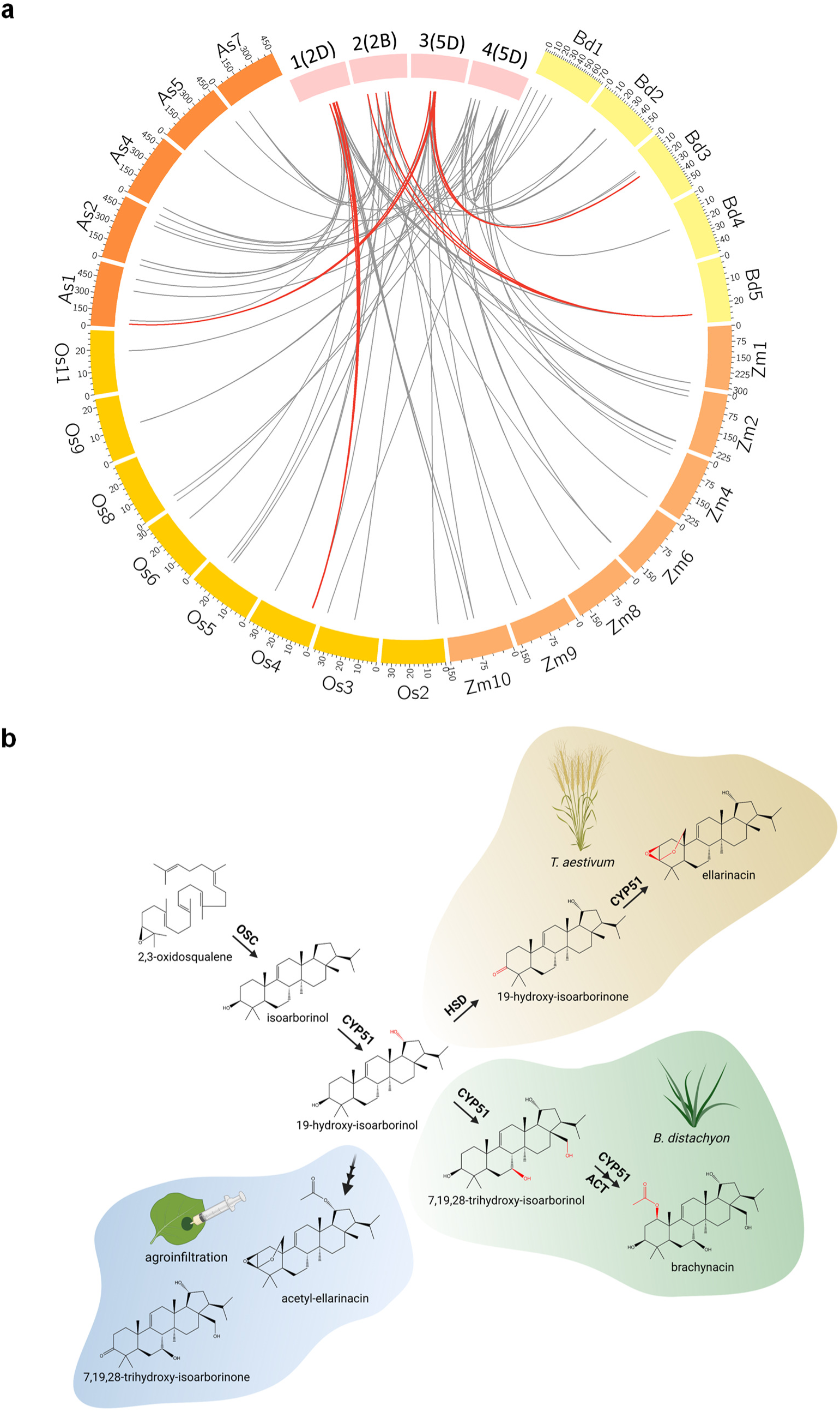
Phylogenetic and chemical divergence of arborinane-type biosynthetic gene clusters. **a,** Circos plot depicting genomic locations of closest matching homologs of co-expressed genes from wheat BGCs 1(2D), 2(2B), 3(5D) and 4(5D) on chromosomes of *B. distachyon* (Bd), diploid oat *Avena strigosa* (As), maize (Zm) or rice (Os). In grey: links to homologs dispersed across the analyzed genomes. In red: links where two or more matching homologs from different gene families co-localize in the analyzed genomes. Cluster 1(2D) is linked to momilactone BGC on rice Chr.4; cluster 2(2B) is linked to a putative terpene BGC in *B. distachyon* Chr.5; cluster 3(5D) is linked to the brachynacin BGC in *B. distachyon* Chr.3 and OSC-CYP51H pair in oat Chr.1; cluster 4(5D) homologs are dispersed in all grass genomes included in the analysis. **b**, clustered biosynthetic pathways for arborinane-type triterpenoids in wheat and *B. distachyon* diverge from a common precursor, 19-hydroxy-isoarborinol, due to neofunctionalization of CYP51 enzymes and recruitment of other gene families. These pathways can be further artificially ‘diverged’ by recombinant expression of combined genes from the two clusters. Image created with Biorender.

Finally, a pathogen-induced cluster (BGC 3(5A/5D)) for a novel isoarborinol-derived triterpenoid, ellarinacin, was found in Chr.5 of the A and D genomes, which is conserved in its wild ancestral species, wild emmer wheat and *A. tauschii*, and is comprised of genes co-opted from sterol primary metabolism. Interestingly, we found the ellarinacin BGC to be more highly induced by non-adapted strains of several fungal pathogens. This cluster may thus form part of a wider set of defense responses found to be actively suppressed in wheat by adapted fungal pathogens, presumably via suppression of plant immune response regulators by pathogen-secreted effector proteins^39^.

Microsynteny and homology searches in other grasses revealed the occurrence of a pathogen-induced cluster in *B. distachyon,* homologous to the ellarinacin cluster in wheat. The pathways encoded by these two clusters diverge from the shared intermediate 19-hydroxy-isoarborinol by neofunctionalization of duplicated CYP51 genes, together with recruitment of additional genes for other enzyme families. Recombinant expression experiments showed that at least some components of these clusters are interchangeable, enabling production of molecules that are not produced by either cluster alone (Fig. 7b), and pointing to the importance of enzyme promiscuity in facilitating chemical diversification.

In summary, a genomics-driven approach has enabled us to rapidly identify and characterize novel compounds and biosynthetic pathways in bread wheat. These clusters are highly induced in response to infection by various fungal pathogens and PAMPs, suggesting a broad-spectrum role for these clusters in chemical defense against biotic stresses. Correspondingly, co-expressed genes within these clusters were found to be part of a shared regulatory network that includes various transcription factors predicted to be associated with biotic stress responses. Future work is needed to further understand the interactions and potential contribution of each of these pathways to protection from pathogens in wheat and other grasses, as well to elucidate the regulatory network which governs the expression of these pathways.

## MATERIALS AND METHODS

### Regulatory network analysis

Target gene-transcription factor interactions and GO term enrichment tables were extracted from a GENIE3-generated wheat regulatory network^8^, available at https://doi.org/10.5447/ipk/2018/7. Network visualization was done with Cytoscape v3.8^54^. Genbank accessions for benzoxazinoid pathway genes were retrieved from^55^ and matched with IWGSC gene IDs by BlastN on EnsemblPlants (http://plants.ensembl.org). WGCNA and GENIE3-generated regulatory network^8^ were generated using IWGSC RefSeq v1.0 gene models. Other analyses described in this manuscript are based on RefSeq v1.1 gene models.

### Co-expression analysis

Co-expression within each cluster was assessed by calculation of the Pearson correlation coefficient (r-val) between the expression of a representative scaffold-forming gene from each cluster (*i.e*., TPS in clusters 1(2A), 1(2D) and 2(2B), OSC in clusters 3(5A) and 3(5D), and chalcone synthase in cluster 4(5D)), and other genes in the cluster, within an RNA-seq dataset including 68 experiments from the wheat-expression.com website^8,19^.

### Pairwise alignment with orthologous clusters in wheat ancestral species

Peptide sequences were extracted from EnsemblPlants (http://plants.ensembl.org): *Aegilops tauschii* (Aet_v4.0)^56^, *Triticum turgidum subsp. diccocoides* (WEWSeq_v1.0)^57^, *Triticum aestivum* (IWGSC)^6^. Gene models were manually corrected to obtain full coding sequences, and putative protein sequences were aligned using LALIGN (http://www.ebi.ac.uk).

### Microsynteny analyses

To perform microsynteny analysis and generate figures, a python implementation of MCScan^58^, https://github.com/tanghaibao/jcvi/wiki/MCscan-(Python-version), was used. FASTA and GFF3 files were retrieved from EnsemblPlants (http://plants.ensembl.org) for chromosomes 5A, 5B and 5D of *Triticum aestivum* (IWGSC), 5A and 5D of *Triticum turgidum subsp. diccocoides* (WEWSeq_v.1.0) and 5D of *Aegilops tauschii* (Aet_v4.0). MCScan ortholog finding and synteny assignment was run with a c-score of 0.99 and a single iteration. For wheat-rice analysis, wheat Triticum_aestivum_4.0^59^ and rice IRGSP-1.0^60^ assemblies were used.

### Semiquantitative RT-PCR in wheat tissues

Bread wheat cv. ‘Chinese Spring’ plants grown in hydroponic cultures were used for collection of coleoptile, root and root tip (5 mm terminal sections) tissues. Sterilized seeds were transferred to sterile polyacrylamide beads (Scotts) equilibrated with Hoagland’s medium #2. Tissues were harvested after 5 days of incubation under controlled conditions (16 h/ 8 h light/dark photoperiod, 23°C). Tissues of stem, inflorescence and leaf infected with mildew (*Blumeria graminis f. sp. tritici* isolate FAL92315) were harvested from 9-week-old plants grown in a greenhouse. Leaf and wounded leaf tissues were harvested from 12-day-old plants grown in a controlled environment (16 h/ 8 h light/dark photoperiod, 18°C during daytime, 13°C at night). Wounded leaf tissue was collected 3 h after wounding with forceps. For root tips infected with *Gaeumannomyces graminis* (Take-all), tissues were collected from sterile plants germinated on a fungus-containing substrate. 5 mm terminal root sections were collected 6 d after sowing. For preparation of substrate, fungus was grown in 200 ml PD medium at 22°C/130 rpm for 4 d, mycelium washed 5 times with Hoaglands medium #2 and mixed with 20 ml polyacrylate beads (prepared by addition of 5 gr of cross-linked polyacrylate (Miracle-Gro) to 250 ml of Hoagland’s medium #2, equilibration of hydrated beads with 3 x 250 ml of medium and subsequent autoclaving). RNA from all samples was extracted using TRI reagent (Sigma-Aldrich), according to manufacturer’s protocol. 5 µg total RNA of each sample was used in 20 µl reverse transcription reactions with Superscript III (Thermo Fisher Scientific), according to manufacturer’s protocol. 30-cycle PCR reactions containing 0.2 µl cDNA template in 10 µl total reaction volume were performed and analyzed on 1% agarose gels. Oligonucleotides used are specified in Supplementary Table 8.

### Semiquantitative RT-PCR in *Brachypodium distachyon* tissues

Tissues for RT-PCR were sampled from greenhouse-grown *B. distachyon* (Bd21) plants or 2-day-old seedlings grown on a Petri dish in a growth cabinet (28°C, 16 h photoperiod). RNA was extracted using RNeasy plant mini kit (Qiagen), treated with DNAse (RQ1, Promega) and used for cDNA library preparation with GoScript reverse transcriptase (Promega), using oligo(dT) primers. All PCR reactions were carried out on Eppendorf Mastercycler pro thermal cycler, for 40 cycles with 55°C annealing temp., using GoTaq G2 Green Master Mix (Promega) and oligonucleotides detailed in Supplementary Table 8. Electrophoresis of PCR products was done on EtBr-stained 1% agarose gels and photographed on a Gel Doc XR instrument (Bio-Rad).

### Inoculation of detached wheat leaves with powdery mildew

For gene expression profiling by qRT-PCR, detached leaves from 10-day-old Chinese Spring wheat plants, grown in a growth cabinet (18°C, 16 h day-length under fluorescent lights supplemented with near-UV lights and 12°C for 8 h in the dark), were inoculated with *Blumeria graminis f. sp. tritici* (isolate FAL92315, maintained on the susceptible wheat cv. Cerco), or with *Blumeria graminis f. sp. hordei* (CH4.8 isolate, maintained on the susceptible barley cv. Golden Promise). Non-inoculated detached leaves kept in same conditions were used as controls. Leaf segments of ∼4 cm length were placed in boxes containing water with 0.5% agar and 100 mg L^-1^ benzimidazole, and were inoculated by blowing fresh spores into settling towers placed over the plant material, according to the method of^61^. Following inoculation, plant material was kept in growth cabinet at constant temperature of 15°C and 16 h day-length, and samples collected 12 h and 24 h post-inoculation.

### Treatment of detached wheat leaves with elicitors

2-3 cm leaf sections were cut from 1^st^ leaf of 10-day old Chinese Spring seedlings grown in soil. Leaf sections were kept in H_2_O in a Petri dish for 24 hours in a 22°C lighted growth cabinet (16 h/ 8 h light/dark photoperiod), then transferred to Petri dishes containing different solutions and kept in same cabinet: 150 μM methyl jasmonate (Sigma-Aldrich), 500 μM salicylic acid, pH 6.0 (Sigma-Aldrich), 0.5 mg/ml chitin (NaCoSy) or H_2_O. All solutions also contained 0.02% Tween-20. Samples for qRT-PCR analysis were collected after 2 h or 12 h. Four biological replicates of MeJa-treated leaves were collected after three days of treatment for GC-MS analyses.

### Treatment of *B. distachyon* with methyl jasmonate

Sections were cut from aerial parts of *B. distachyon* Bd21 plants grown in soil for 2.5 weeks. Samples were kept in H_2_O in a Petri dish for 24 h in a 22°C lighted growth cabinet (16 h photoperiod), then transferred to Petri dishes containing 150 μM methyl jasmonate and 0.02% Tween-20, or 0.02% Tween-20 in H_2_O, and kept in same cabinet. Four biological replicates of samples were collected for GC-MS analysis after three days.

### Quantitative real-time PCR (qRT-PCR) of wheat

For qRT-PCR analysis of wheat leaves inoculated with powdery mildew or treated with elicitors, three biological replicates, each containing three leaf samples, were tested for each time point. RNA was extracted using TRI reagent (Sigma-Aldrich), according to manufacturer’s protocol. Following DNase treatment (RQ1, Promega), RNA was reverse-transcribed with M-MLV reverse-transcriptase (ThermoFisher Scientific) using a 1:1 mix of random hexamers and oligo(dT) primers. All oligonucleotides (Supplementary Table 8) were designed using Primer3 software^62^, with at least one homoeolog-specific oligo per each pair used. qRT-PCR was performed on a CFX96 Touch Real-Time PCR instrument (Bio-Rad) in the following conditions: initial step in the thermal cycler for 3 min at 95°C, followed by PCR amplification for 40 cycles of 10 s at 95°C and 30 s at 59°C, and finally dissociation analysis to confirm the specificity of PCR products with 0.5°C ramping from 55°C to 95°C. Each 10 μl reaction was comprised of 5 μl LightCycler 480 SYBR Green I Master mix (Roche Life Science), 2 μl cDNA template, 2 μl H_2_O and 1 μl primer mix (0.5 μM each primer). Relative transcript levels were calculated according to the Pfaffl method^63^, using the housekeeping gene β-tubulin (TUBB) as reference^64^.

### Quantitative real-time PCR (qRT-PCR) of *Brachypodium distachyon*

*B. distachyon* accession Bd3-1 seeds were soaked, peeled, and placed between three filter paper layers soaked in 5 ml water. The seeds were stratified for 5 days at 5°C in the dark and one day at 22°C (16h/8h - light/dark photoperiod) in a controlled environment growth cabinet. For Fusarium root rot (FRR) material, ten stratified seeds were placed on 9 cm^2^ filter square paper on 50 ml 0.8% water agar. All plates were placed in a plant propagator with water-soaked paper towels, angled 20° from the upright position, and stored for 3 days at 22°C (16 h/ 8 h - light/dark photoperiod, variable humidity). *Fusarium graminearum* isolate PH1 was maintained on potato dextrose agar (PDA) at 22°C 16 h/ 8 h - light/dark photoperiod in a controlled environment growth cabinet. One 9 cm diameter Petri-dish of seven-day old *F. graminearum* mycelia was blended to a slurry with 1 ml water and applied to three points on each root (root tip, mid root, and near seed) using a 10 ml syringe. The inoculum slurry was removed once infection was visible at 1 dpi and the roots were rinsed with sterile distilled water. Immediately after and then every two days, ten roots per plate (one biological replicate pool) were cut and flash frozen in liquid nitrogen. For Fusarium Head Blight (FHB) material, seeds were sown in 50% peat/sand and 50% John Innes mix 2 (two seeds per 8 cm^2^ pot). Plants were then maintained for six weeks at 22°C (20 h/ 4 h - light/dark photoperiod, 70% humidity) in controlled environment growth cabinet until mid-anthesis. Before the dark period, pots and matting was watered until run-off, spikes were inoculated with 1 x 10^6^ spores/cm^2^ amended with 0.05% Tween-20, and all plants were enclosed in clear plastic bags to maintain high humidity for three days. Immediately after and then every two days, three spikes from different plants were pooled and flash frozen in liquid nitrogen. For conidial suspension inoculum, Mung Bean (MB) broth^65^ with a 1 cm^2^ *F. graminearum* PDA mycelial plug was incubated at 23-25°C, 200 rpm for seven days. The inoculum was filtered with cheesecloth and quantified using a haemocytometer.

RNA from FHB, FRR, and control samples was extracted using a QIAGEN RNAeasy plant mini kit as per standard protocol. RNA was then immediately cleaned using Turbo DNA-free kits (Invitrogen) as per standard protocol except for two rounds of Turbo DNAse treatment. Subsequently cDNA was prepared using SuperScript III Reverse Transcriptase (Thermo Fisher Scientific), as per standard protocol. All oligonucleotides (Supplementary Table 8) were designed using Primer3 software^62^. Reverse transcriptase qPCR was performed in a Framestar-480/384 well plate containing 5 µl of 2x SYBR Green JumpStart Taq ReadyMix (Sigma-Aldrich), 2 µl cDNA, 0.6 µl of 10 µM per primer, and 1.8 µl water per well. The thermocycling protocol 300 s 95°C, 45x(94°C 10 s, 58°C 10 s, 72°C 10 s, 75°C 2 s (single acquisition)), followed by dissociation analysis by ramping from 65°C to 97°C, was performed on a Roche LightCycler LC480. Cq values and primer efficiency were quantified using the LinRegPCR tool (Amsterdam UMC Heart Failure Research Centre). Relative quantification was calculated according to the Pfaffl method^63^, using the housekeeping gene GAPDH as reference.

### Generation of DNA constructs

For cloning of *Brachypodium distachyon* and wheat genes, RNA was extracted from mature plant leaves of *B. distachyon* (accession Bd21) or leaves from 10-day-old *Triticum aestivum* plants (Chinese Spring), infected with powdery mildew (*Blumeria graminis f. sp. tritici*), using RNeasy plant mini kit (Qiagen). RNA was treated with RQ1 DNAse (Promega) and cDNA libraries prepared with Superscript IV or Superscript III reverse transcriptase kits (Thermo Fisher Scientific), using oligo(dT) primers, according to manufacturer’s protocols. TaOMT3, TaOMT6, TaOMT8, TaCYP71C164_5D, TaCYP71F53_5D, BdOSC2, CYP51H13P/H13_5A/14/H15/H16/H35/H37 were amplified from cDNA using Phusion DNA polymerase (Thermo Fisher Scientific) or Q5 DNA polymerase (New England Biolabs). TaOSC_5D, TaHSD, AsOSC1, AsCYP51H73 were synthesized by General Biosystems, Durham, NC, USA. BdACT, BdMeTr, TaCHS1, chi-1D, TaCPS-D2, TaKSL-D1 and *Taxus canadensis* GGPPS were synthesized by Twist Bioscience, San Francisco, CA, USA. TaCPS-D2, TaKSL-D1 and TcGGPPS lack signal sequences, to allow for cytosolic localization in *N. benthamiama* expression^66^. *A. tauschii* IAS coding sequence was derived from TaOSC_5D sequence with site directed mutagenesis^67^ to obtain a single mutation (I581S). Synthesized and cDNA-amplified genes from triterpene and diterpene BGCs were cloned into a pCAMBIA-based^68^ plant expression vector with Goldenbraid cloning^69^, using BsaI and BsmbI (New England Biolabs) and T4 DNA ligase enzymes (New England Biolabs). Gene expression in final vectors is driven by *Solanum lycopersicum* ubiquitin 10 promoter and terminator^70^. Synthesized and cDNA-amplified genes from flavonoid cluster were cloned into a pDONR207 Gateway entry vector and subcloned into a pEAQ-HT-DEST1 plasmid^71^ using BP and LR clonase enzyme mixes (Thermo Fisher Scientific), respectively. Full coding sequences of all synthesized and PCR-cloned genes used in this study are found in Supplementary Methods. Oligonucleotides used for amplification and sub-cloning are specified in Supplementary Table 8.

### Agroinfiltration-mediated transient expression in *N. benthamiana*

Plant expression vectors were transformed into *Agrobacterium tumefaciens* GV3101 via electroporation. Agrobacteria cultures were grown overnight in 28°C in LB media and resuspended in MMA buffer (10 mM MgCl_2_, 10 mM MES pH 5.6, 100 µM acetosyringone) to O.D._600_ 0.2. For co-expression of several genes, O.D._600_ 0.2 cultures of strains expressing different genes were mixed 1:1 prior to infiltration. Cultures were infiltrated by syringe into leaves of 5 weeks old greenhouse-grown *N. benthamiana* plants. The plants were further maintained in the greenhouse after infiltration. Infiltrated leaves were harvested 5 days post infection, freeze-dried and ground.

### Metabolite extraction from agroinfiltrated *N. benthamiana* leaves for GC-MS analyses

Diterpenes: for analysis of TaCPS-D2 and TaKSL-D1 transient expression, 5 mg of *N. benthamiana* leaf samples were extracted in 850 µl ethyl acetate for 1 h in room temperature, with agitation. Following removal of plant tissue by centrifugation, 750 µl from each extract was evaporated and reconstituted in 75 µl ethyl acetate. Triterpenes: for analysis of wheat BGC 3(5D) genes expression, 5 mg samples were extracted in 500 µl ethyl acetate with 5 µg/ml 5α-cholestan-3β-ol. For analysis of *B. distachyon* brachynacin cluster genes expression, 5 mg samples were extracted in 500 µl methanol with 5 µg/ml 5α-cholestan-3β-ol. For analysis of oat and *A. tauschii* OSC and CYP51 genes, 5 mg samples were extracted in 500 µl ethyl acetate. For analysis of combined expression of wheat BGC 3(5D) genes and BdACT, 5 mg samples were extracted in 300 µl ethyl acetate. For analysis of combined expression of *B. distachyon* brachynacin cluster genes and TaHSD, 5 mg samples were extracted in 500 µl ethyl acetate with 5 µg/ml 5α-cholestan-3β-ol. All triterpene extractions from *N. benthamiana* leaves were done in room temperature for 1 hour, with agitation. For all triterpene extractions, following the removal of plant tissue by centrifugation, 200 µl were evaporated and reconstituted in 70 µl TMS+pyridine (Sigma-Aldrich). Samples were derivatized for 0.5 h in 70°C.

### Metabolite extraction from MeJA-treated wheat and *B. distachyon* leaves for GC-MS analyses

MeJa-treated wheat leaf sections were freeze-dried and ground. 25 mg from each sample were extracted in 800 µl ethyl acetate containing 5 µg/ml 5α-cholestan-3β-ol, with agitation for 2 h in 40°C. Following removal of tissue by centrifugation and filtration with 0.22 µl filter mini columns (Norgen), 700 µl from each extract was evaporated and reconstituted in 70 µl TMS with pyridine (Sigma-Aldrich). Samples were derivatized for 0.5 h in 70°C. MeJa-treated *B. distachyon* leaf sections were freeze-dried and ground. 25 mg from each ground sample were extracted in 1100 µl methanol containing 2.5 µg/ml 5α-cholestan-3β-ol, with agitation for 2 h in 40°C. Following removal of tissue by centrifugation and filtration, 800 µl from each extract was evaporated and reconstituted in 70 µl TMS with pyridine (Sigma-Aldrich). Samples were derivatized for 0.5 h in 70°C.

### GC-MS analysis of diterpenes and triterpenes from *N. benthamiana* and grasses leaf extracts

GC-MS analysis was performed using an Agilent 7890B instrument with a Zebron ZB5-HT Inferno column (Phenomenex). For triterpenes analysis, a previously described method^30^ was used: injections were performed in pulsed splitless mode (30 psi pulse pressure). Inlet temperature was set to 250°C.

GC oven temperature was initially held at 170°C for 2 mins, subsequently ramped to 300°C at 20°C/min and held at 300°C for an additional 11.5 min (20 min total run time). The GC oven was coupled to an Agilent 5977B MS detector set to scan mode, from 60 to 800 mass units (solvent delay 8 min). For semi-quantification of brachynacin in *B. distachyon* leaves, Selected Ion Monitoring (SIM) mode was used, for detection of brachynacin (m/z 170.1, 340.2, 387.3, 400.3, 445.4, 475.4, 500.4 ions were monitored) and internal standard 5α-cholestan-3β-ol (m/z 215.1, 355.4, 445.5, 460.5 ions were monitored), with 100 ms dwell time for each ion. Diterpenes analysis was based on a previously described method^66^: injections were performed in splitless mode. Inlet temperature was set to 280°C. GC oven temperature was initially held at 130°C for 2 mins, subsequently ramped up to 250°C at 8°C/min, followed by ramping up to 310°C at 10°C/min and held at 310°C for an additional 5 min (28 min total run time). The MS detector was set to scan mode, from 50 to 550 mass units (solvent delay 4 min).

### Metabolite extraction from agroinfiltrated *N. benthamiana* leaves for LC-MS analyses

For LC-MS analysis of recombinantly expressed wheat BGC 3(5D) genes (including combined expression with BdACT), 25 mg of each sample were extracted in 2 ml methanol in room temperature for 1 h, with agitation. Following removal of plant tissue by centrifugation, extracts were partitioned twice with 2 ml hexane and filtered with 0.22 µl filter mini columns (Norgen). Extracts were evaporated and resuspended in 100 µl methanol. For analysis of recombinantly expressed *B. distachyon* brachynacin cluster genes (including combined expression with TaHSD), 10 mg of each sample were extracted in 400 µl 80% methanol in room temperature for 1 h, with agitation. Following removal of plant tissue by centrifugation, extracts were partitioned with 500 µl hexane and filtered. Extracts were evaporated and resuspended in 100 µl 80% methanol. For analysis of wheat flavonoid cluster genes, 250 mg freeze-dried and ground samples were extracted with 4 mL methanol at room temperature for 1 h. Extracts were fully evaporated, resuspended in 200 μL methanol, and filtrated through a mini column (pore size 0.22 µm, Geneflow). Filtered samples were transferred to glass autosampler vials and 20 μL of each sample was analyzed by UHPLC-CAD-PDA-MS.

### LC-MS analyses of triterpenes from *N. benthamiana* leaf extracts

Leaf extracts were analyzed by reverse phase HPLC on a Shimadzu LCMS-2020 single quadrupole mass spectrometer coupled with a Dionex Corona Veo RS charged aerosol detector (Thermo Scientific), using a Kinetex 2.6 µm XB-C18 100 Å, 50 x 2.1 mm LC Column (Phenomenex). MS data was collected using combined electrospray ionization (ESI) and atmospheric pressure chemical ionization (APCI) in positive mode. 10 µl samples were injected using 12 min, 14.5 min or 30 min mobile phase gradient methods using solvent A- water with 0.1% formic acid and solvent B- methanol with 0.1% formic acid, as follows. 12 min method: 50% B hold from 0 to 0.75 min, 50% to 90% B from 0.75 to 8 min, 90% B hold from 8 to 10 min, 90% to 50% B from 10 to 10.5 min, 50% B hold from 10.5 to 12 min. Flow rate, 0.4 ml/min. MS scan, m/z 250 – 1900. Column oven temperature, 40°C. 14.5 min method: 70% to 95% B from 0 to 10 min, 95% B hold from 10 to 11 min, 95% to 70% B from 11 to 11.1 min, 70% B hold from 11.1 to 14.5 min. Flow rate, 0.5 ml/min. MS scan, m/z 200 – 1200. Column oven temperature, 30°C. 30 min method: 15% B hold from 0 to 0.15 min, 15% to 60% B from 0.15 to 26 min, 60% to 100% B from 26 to 26.5 min, 100% B hold from 26.5 to 28.5 min, 100% to 15% B from 28.5 to 29 min, 15% B hold from 29 to 30 min. Flow rate, 0.3 ml/min. MS scan, m/z 100 – 1500. Column oven temperature, 30°C.

### LC-MS analyses of flavonoids from *N. benthamiana* leaf extracts

Leaf extracts were analyzed by reverse phase HPLC on a Shimadzu LCMS-2020 single quadrupole mass spectrometer coupled with a Dionex Corona Veo RS charged aerosol detector (Thermo Scientific) and a SPD-M20A HPLC Photodiode Array Detector (PDA; Shimadzu), using a Kinetex 2.6 µm XB-C18 100 Å, 50 x 2.1 mm LC Column (Phenomenex), kept at 30°C. Water containing 0.1% formic acid (FA) and acetonitrile containing 0.1% formic acid (FA) were used as mobile phases A and B, respectively, with a flow rate of 0.2 mL/min. A gradient elution program was applied as follows: 0-1.5 min linearly increased from 0% to 10% B, 1.5-26 min linearly increased from 10% to 60% B, 26-26.5 min linearly increased from 60% to 80% B, 26.5-28.5 min linearly increased from 80% to 100% B, 28.5-29 min linearly decreased from 100% to 10% B hold on for another 1 min for re-equilibration, giving a total run time 30 min. MS detection was performed in both positive and negative ESI range of *m/z* 50–1500 with the following settings: desolvation temperature was 250°C; drying gas flow, 15 L/min; detector voltage was 1.25 kV; and nebulizing gas flow, 1.5 L/min. PDA chromatograms were recorded in a 200–600 nm range using a deuterium (D2) and tungsten (W) light source.

High-resolution mass spectrometry analysis of the metabolites was carried out on a Q Exactive instrument (Thermo Scientific). Chromatography was performed using a Kinetex 2.6 μm XB-C18 100 Å, 50 mm x 2.1 mm (Phenomenex) column kept at 30°C. Water containing 0.1% formic acid (FA) and acetonitrile containing 0.1% formic acid (FA) were used as mobile phases A and B, respectively with a flow rate of 0.4 mL/min. A gradient elution program was applied as follows: 0-0.75 min linearly increased from 0% to 10% B, 0.75-13 min linearly increased from 10% to 60% B, 13-13.25 min linearly increased from 60% to 80% B, 13.25-14.25 min linearly increased from 80% to 100% B, 14.25-14.5 min linearly decreased from 100% to 10% B hold on for another 2.5 min for re-equilibration, giving a total run time 17 min. MS detection was performed in both positive and negative ESI range of 100–1500 *m/z*. The analysis of the 3D field of the Photodiode-Array Detection (PDA) was recorded in a 200–600 nm range using a vanquish detector (Thermo Scientific).

### Large-scale agroinfiltration, extraction and purification of triterpenoids

Vacuum-mediated large-scale agroinfiltrations of *N. benthamiana* plants, and downstream extraction and purification of triterpenoid products were based on a previously described method^30,72^. Specific methods for extraction and purification of the metabolites are detailed in Supplementary Methods.

### General considerations for NMR

NMR spectra were recorded in Fourier transform mode at a nominal frequency of 600 MHz for ^1^H NMR, and 150 MHz for ^13^C NMR (unless specified otherwise), using the specified deuterated solvent. Chemical shifts were recorded in ppm and referenced to the residual solvent peak or to an internal TMS standard. Multiplicities are described as, s = singlet, d = doublet, dd = doublet of doublets, dt = doublet of triplets, t = triplet, q = quartet, quint = quintet, tquin = triplet of quintets, m = multiplet, br = broad, appt = apparent; coupling constants are reported in hertz as observed and not corrected for second order effects.

### Genomic positioning of wheat BGC homologs in other grasses

Protein sequences of all co-expressed genes from wheat BGCs 1(2D), 2(2B), 3(5D) and 4(5D) were used as BlastP queries against the following genome assemblies: *Zea mays* B73 RefGen_v4^73^, *Hordeum vulgare* cv. Morex r1^74^, *Brachypodium distachyon* Bd21 v3.1^44^, *Oryza sativa ssp. japonica* cv. Nipponbare v7.0^75^ and *Avena strigosa* S75 v2.0^43^. BlastP searches in all assemblies except *Avena strigosa* were performed in Phytozome13 (https://phytozome-next.jgi.doe.gov/)^45^, using default parameters. Genomic locations of top BlastP hits in each species were visualized using Circos software v0.69-9^76^.

## ACKNOWLEDGEMENTS

We would like to thank Paul Brett, Lionel Hill, Amr El-Demerdash, James Reed, Hannah Hodgson, Rebecca Casson and Sergey Nepogodiev for assistance and advice on analytical chemistry analyses, Andrew Steed and Rachel Burns for assistance with wheat and *B. distachyon* pathogen infections, Nikolai Adamski for helpful discussions, JIC horticultural services staff for assistance with plant cultivation, Noam Chayut and Simon Orford (JIC Germplasm Resource Unit) for providing ‘Chinese Spring’ seeds, and Christine Faulkner for providing chitin. *Blumeria graminis* isolates CH4.8 and FAL92315 were kindly provided by Lesley Boyd (NIAB). G.P. is supported by a Royal Society Kohn International Fellowship (NIF\R1\180677) and a Marie Skłodowska-Curie Individual Fellowship (838242). A.O.’s lab is supported by the Biological Sciences Research Council (BBSRC)-funded Institute Strategic Programme Grant ‘Molecules from Nature’ (BB/P012523/1) and the John Innes Foundation.

## AUTHOR CONTRIBUTIONS

G.P, M.D and A.O conceived and designed the experiments. G.P, M.D, R.C.M, J.H and L.C performed the experiments. G.P, M.J.S and C.O analyzed the data. R.R.G, H.S, P.B and D.R.N advised, analyzed and contributed data. J.B, P.N, C.U and A.O jointly supervised research. G.P and A.O wrote the manuscript, with contributions from all authors.

