## Supplementary for "Genome Mining Uncovers Clustered Biosynthetic Pathways for Defense-Related Molecules in Bread Wheat"

**Supplementary Methods.** Coding sequences of genes used in this study. Synthesized genes are asterisked.

>TaCHS1* (TraesCS5D02G488700)

ATGGCCGCCGCTGTTACCGTAGACGAAGTAATGAAGATGCAGAGAGCGCAGGGCCCCGCAACTGTGTTAGCAATA
GGTACGGCCACTCCACATAACTGCGTTTACCAAGCAGACTACGCTGACTACTACTTTAGGATCACCAATAGTGAA
CATCTGACAGATCTAAAAGACAAATTCCAAAGAATGTGTGACCGAAGTATGATCAGAAAGCGTTACATGCATGTG
ACCGAGGAGATTCTCAAAGAGAACCCCAACGTTTGCGCTTACATGGCGCCTAGCCTAGATGCTCGCCAAGACATG
GTGCTTGGTGAGGTTCCCAAGCTGGGGAAAGAGGCTGCTCATAAGGCAATCAAGGAATGGGGACAGCCTCTCTCC
AAGATCACTCACCTCGTTTTCTGCACAACGAGTGGAGTGGACATGCCAGGTGCCGACTATCAGCTGACTAAGATG
CTTGGTCTGCACCCATCTGTAAGAAGGGTGATGTTATACCAGCAAGGTTGCTTTGCTGGTGGTACAGTGCTACGT
GTGGCTAAGGACTTAGCCGAGAACAACAGAGGAGCTCGTGTGCTTGTAGTGTGCTCCGAGATAACAGCAGTTACT
TTTCGAGGCCCGTGTGACACCCAGTTGGATTCCATGGTCGGACAGGCTTTGTTTGGCGATGGCGCAGCTGCAGTC
GTTGTGGGCGCTGACCCCGACGTATTGGTGGAACGGCCCTTATTCCAACTTGTATCTGCTTCCCAGACCATATTA
CCAGACACAGATGGTTTCATCAAGGGCCACCTTCGAGAGGTTGGACTTACATTCCACCTTCACAAGGATGTACCT
GTTGCGATATCAAAGAACATACAACAAGCACTTGAGGACGCGTTCGCACCACTCGGCATCGATGACTGGAACTCT
ATATTTTGGGTAGCCCACCCGGGTGGCCCAGCCATTTTGGACATGGTCGAGGCTGAGGCTAAGCTTGACAAACGA
AGAATGAGGGCAACTAGGCACATATTGTCTGAGTACGGGAACATGTCCAGCGCTTGTGTGTTGTTCATCTTAGAC
GAGATGAGGAAGAGGTCTGCAGAGGACGGCCACGCTACCACAGGCGAGGGTCTAGACTGGGGCGTCCTGTTTGGA
TTCGGACCCGGTCTTACGGTAGAGACAGTTGTCCTACACAGTGTGAGGACAACAGCCGCTTGA

>chi-D1* (TraesCS5D02G489000; Genbank JN039039)

ATGGCAGTTTCCGAACTAGAAGTTGATGGTGTTGTATTTCCACCCTTAGCAAGACCACCAGGAAGCGAACATGCG
CATTTCTTGGCTGGTGCCGGTGTTAGAGGCATGGAAATTGGTGGGAATTTTATTAAATTTACAGCTATAGGAGTT
TATTTGCAAGCTGATGCTGCTGTTAGTGCTCTTGCGGCTAAATGGGCTGGAAAACCTGCAGCGGATTTAGCATCA
GATGCAGCTTTCTTTCGTGATGTAGTTACTGGAGAATTTGAAAAGTTTACTCGGGTTACTATGATTTTGCCACTT
ACTGGTGCTCAATATTCTGATAAAGTCACCGAAAATTGTGTTGCGTATTGGAAAGCAACAGGGGTCTATACTGAT
GCTGAAGCAGCTGCTGTTGATAAATTTAAAGAAGCATTTGGCCCTCATAGTTTTGCTCCTGGGGCTTCTATTTTA
TTTACTCATTCACCTGCTGGACTTCTGACAGTGGCTTTTAGCAAAGATTCCTCAGTTCCAGAATCTGGTGGTGTT
GCTATTGATAATGCTAGGTTGTGTGAAGCAGTACTTGAATCTATTATAGGTGAACATGGTGTTTCTCCCGCAGCT
AAACTTTCCTTAGCAACACGTGTTGCAGAACTTCTTGAAGGTGCTGCACTGGCAGGTGGTGAACCTGCTGCTGAA
GCAGTCAGTGTCTCAGTATGA

>TaCYP71C164_5D (TraesCS5D02G488500)

ATGGAAGATCTCGTGAAGAAACCCATGCCAGAGACACCTCCGCGGGCATTGTTTCTGTTCCTATTCTTCCTCTTC
CTCTTGTTTGTAAGCTGGTTCACTGGAAAGACAAGAAAAATGCAGCATCAGCAGCGAGAAAACAACCACCTCCCG
CTCCCGCCTTCGCCGCCGGCACTGCCCATCATTGGTCACCTGCACCTCGTCGGCTCCCTCCCGCACGTCTCTCTC
CGCAGCCTTGCCAGGAAGTATGGTCCCGACATGATGCTTCTGCGCCTGGGCGCCGCGCGGACCCTCGTGGTGTCG
TCGCTGCGTGGCGCAGAGGCAGTGCTGCGCACCCACGACCACATCTTGGCGTCGCGGCCCAGCTCCGTAGTCTCC
GACATCCTCACGTACGGCTCGTCTGATATGGCCTTCGCGCCATACGGCGAGTACTGGCGGCAGGTGAGGAAGCTG
GTCACCACCCACATGTTGAGCGTTAAGAAGGTGCAGTCTTTCCGCAGCGCTGCCATGGAGGAGGTGAGCTTGGTG
ATGGCCAAGATTAATGAGGTGGCCACAACCGATGGTACGGTGGACATGAGAGAGCTTCTCAGCTTATTCGCGAAT
GACATGGGGTGCCGCATCGTATCGGGAAAGTTCTTCCTAAAAGATGGACAGAGCAAGTTGTTCCGGGACCTCACC
AACGATACCTCACGGCTGTTAGGAGGGTTCAACGTGGAGGAGTACTTCCCAGGATTGGCAAGGGTAGGAGTGTTT
AAGAGGGCAGTTTGTGCCCAAGCCGAGAGAGTGAGGAATAGATGGGCTGATCTGCTAGACAAGGTGATCGACGAT
CATGTGAGCAAGGATAAGTCAACGTTTGATCACGAGGATGGTGATTTTGTAGATATTCTGTTGACTTTTCAGCAC
GAGTATGATCTCACAAGAGAGCACATGAAAGCTATCCTGATAGATGTATTTGCCGGTGCAACAGACACATCAGCT
AACGCCCTCGAATTCACCTTGGCCGAGCTCATGAGGAAGCCACGTGTGATGGGGAAGCTACAAGCTGAGGTGAGG
AGTAACGTACCCCAGGGACAAGAAATTGTCAGTGAAATCAACGTGAACAATATGGCATACCTAAGAGCAGTCATA
AAGGAGTCGCTTCGACTGTATCCTATTGCGCCTCTCCTTGCTCCACACCTTGCCATGGATGACTGCAATATTGAT
GGATATATGGTTCCTGCTGGGACGCGTGTCATGGTCAATGCATGGGCCATTGCTAGGGACTCGAGCTTGTGGGAG
GACGCAGAAGAGTTCATACCTGAAAGATTTACAGATGAAGGCAGCGCTATGCATGTCAATTTCAAGGGGAATGAT
TTCCAGTTCTTGCCATTCGGGGCAGGACGAAGGATGTGCCCCGGTATGAACCTCGGAATCGCTAATGTTGAGCTT
ATGTTGGCAAACCTCATGTACCATTTTGACTGGGAACTTCCGCTTGGAGTTGAGAGAGAAGACATTGATATGACA
GCGGTGTTTGGGCTAACCATGCGCCGGAAGGAGAAACTATTGTTAATTCCAAAATCCTGGATGTAG

>TaCYP71F53_5D (TraesCS5D02G487900)

ATGGAGGGTTGGTTAACCTTATGTTTCATAGCCGTATCGACGTTAGTGGCCGTTTGGTTTTCCGGTGGCAAGAGC
AAGCCCAAGAAGCATCTGCCTCCTGGGCCATGGACTCTCCCGGTCATCGGCAGCCTCCACCACGTCCTCAGTGTC
CTCCCACACCGCACCATCACGGAGCTGTGTCGCCGGCATGGGCCACTGATGCTCCTCAAGCTAGGTGAGGTTCCA
ACCGTGGTAGTCTCGAGCGCCGAGGCGGTGGAGCAGGTGATGAAGACCAATGACATCGCCTTCTCGTACCGGCGG
ACCACCGAGTTGCAGGACATCGTCGGCTTCGGCGGCAAGGGCATCATCTTCGCCCCCTACGACAACCGCTGGCGC
CAGATGCGCAAGGTCTGCATCATGGAGCTCCTCAATTCCAAGCAGGTGAAGCGCATGGAAGACATCAGGGCCGAG
GAGGTGGGCCGCCTCCTCCGCTCAATCATCACGGCCACAGCTGTGGGCGCTACCGTCAACATCAGCCAGAAGGCT
GCGGCGCTCAGCAATGCCGTGGTGACGCGGGCGGTGTTCGGCGGCAAGTTCGCACGGCAGGAGGAGTACCTTCGC
GAGATCAACCAACTCCTAGAGCTGTTGGGAGGATTCTGCCTTGTCGACCTCTTCCCGTCGTCGCGGCTGGTGCGG
TGGTTCAGCACTTGCGAGCGCCGCACAAAGAAGAGCTGCGACCTCATCCAGCACATCATCACCGGGGTGCTTGAC
GAGCGCAAGGTGGTGCGAGCTGCCGGTGACGGCGCCTGCAGCACCGACGATGAGGACCTACTGGACGTGCTGCTC
AGGCTGCAGGAAGAGGACTCGTTGGCATACCCTCTAACGACAGAGAATATAACTACCGTCTTGTTTGACATCTTT
GCAGCTGGCACGGATACTACAGGAACCGCTTTGGAGTGGGCTATGTCAGAACTCATATGTCATCCTGAAGCTATG
GCTAAGGCACAATTAGAGGTTCGAGAGGTACTGGGTCATGGCCGAGCTATCATTGGCAATAGCGATCTTGCAAAA
CTCCACTACTTGCGGATGGTCATCAAGGAGGTTCTTAGATTGCATCCACCTGGTGCTCTACTTCCCCGCAAGACT
AGAGAGGACTGCAAAATTATGGGTTATGACATGCTTAAAGATACAAATATATACATTAATGTCTTCGCAATTTCC
CGAGATCCTCGATATTGGAACAATCCTGAAGAGTTTAATCCAGGAAGGTTTGAGAACAATAACGTGGATTATAAT
GGGAATTCTTTTGAATTCACTCCTTTCGGAGGTGGGCGACGGCAATGCCCTGGGATAGCATTCGCCACGTCACTT
TTGGAGATCACTTTAGCAAACTTTTTATATCACTTCAACTGGATGCTTCCTGGCGAAGCCAGCTCAGCGTCACTG
GATATGTCTGAGAAATTTGGGTTCACCATAGGTAGAACATCTAATCTGCACCTCAAGGCTATTCCATATGTACGC
TCCACTATATAG

>TaOMT3 (TraesCS5D02G488800)

ATGGGCTCCACTGCCGTGGAGAAGGTCGCTGTCGCCACTGGCGACGAGGAGGCGTGCATGTACGCGGTGAAGCTT
GCAGCGGCATCTATCCTTCCAATGACCCTCAAGAACGCCATCGAGCTGGGCATGCTCGAGATCCTCGTGGGTGCC
GGCGGGAAGATGTTGTCACCTTCAGAGGTGGCAGCGCAGCTTCCGTCGAAGGCCAACCCGGAGGCACCGGTTATG
GTGGACCGCATGCTGCGGCTGCTGGCATCGAACAACGTCGTGTCATGCGAGGTGGAGGAAGGTAAGGACGGCCTC
CTCGCCCGTCGATACGGCCCCGCGCCCGTGTGTAAGTGGCTCACACCCAACGAGGACGGCGCATCCATGGCTGGG
CTGCTCCTCATGACCCACGACAAGGTCACTATGGAGAGCTGGTATTATTTGAAGGACGTGGCCCTTGAAGGCGGC
CAACCATTCCACAGGGCGCACGGGATGACGGCGTACGAGTACAACAGCACAGACCCACGCGCTAACTGCTTGTTC
AACGAGGCCATGCTTAACCACTCCACCATCATCACCAAGAAGCTCCTCGAGTTCTACAGGGGCTTCGACAACGTC
GAGACCCTCGTGGATGTCGCCGGTGGCGTTGGTGCCACAGCCCACGCCATCACCTCAAAGTACCCGCACATCAAG
GGGGTAAACTTCGATCTCCCGCATGTCATATCCGAGGCGCCGCCCTACCCTGGCGTGCAGCACATCGCCGGTGAC
ATGTTCAAGAAGGTGCCCTCCGGCGATGCTATCCTCCTGAAGTGGATCCTCCACAACTGGACCGACGATTACTGT
ATGACTCTTCTGAGGAACTGCTACGATGCGTTGCCCATGAATGGCAAGGTGGTCATCGTGGAGGGCATCCTGCCG
GTGAAACCAGATGCAATGCCCAGCACGCAGACGATGTTCCAGGTCGACATGATGATGCTGCTGCACACCGCAGGC
GGCAAGGAGAGGGAACTGAGCGAATTTGAAGAGCTAGCGAAGGGCGCTGGGTTCAGCACAGTCAAGACCAGCTAC
ATCTACAGCACCGCATGGGTCATTGAGTTCGTCAAATAG

>TaOMT6 (TraesCS5D02G488300)

ATGGCGCCCACCCAAGGCAAGCAGAGTTCTCAGGATTTGCTCGAGGCTCAAGTTGACCTTTGGCACCATTCATTG
GGATTTGTCAAGTCCATGGCACTCAAATGTGCAATGGAACTGCAAATCCCTAACACCATCCAACACCATGGTGGG
GCTATGACTCTTTCTGAGCTGGCCACAAAGACAGGGATCCATCCGTCTAAGCTTCCCCCCTTAGGGCGACTCATG
CGTGTACTCACCGTATCAGGCATCTTTGTTGTCCATGACTCAGCCTCGGGAGACAACAAGGAGGCTGTGTATGGA
CTTACCCCAAGCACGTGCCTCCTCGTCAGCGATGAAGTTAAATCAAACCTATTTCCCATTCTGTCTTTGTTCCTT
GATTCAACTGCCATTACACCCTTTTTCGGCATGCACTCATGGTTCCTAGATGAGCATTCCACGTCCCTGTTCAAA
AAGGCTCACGGCCTTAACGTTTGGGAGATGGCTGACCAGAACAATGCTTGCAACGAGCAAATCAATAACGCGATG
GTTTCTGATAGTAACTTTTTCATCAATATCCTCTTGAGGGAGTGTGGTGATGTATTTCTTGGCATAAACTCACTT
ATTGATGTTGCGGGAGGACACGGTGGAGCTGCTAGGGCAATTGCTATGTCGTTCCCCCAGATGAAATGCACTGTG
TTGGATCTCCCTCATGTGGTTGCTGGAGCTCCCAGTGATGCCCATGTGTCATTTATTTCTGGCGATATGTTTAAG
TACATTCCACCGGCGGATGCTCTTTTCCTGAAGTGCATTTTTCACGATTGGTGTGATGAAGACTGTGTCAAGCTA
CTGAAGAATTGCAAGGAAGCTATCCCTCCCAGAGATGCTGGTGGAAAGGTGATAATCGTTGATATGGTGGTTGGA
TCTGGGCCAGATGACATTGTAACAAGAGAGACACAGGTTTTCTATGATCTCTTTATCATGGGTGCTGAGGGGATT
GAGCGAGAGGAATTTGAGTGGAAGAAGATATTTATGGAAGCTGGGTTCAGCAATTACAAAATTCTATCAGTGCTG
GGAGTTAGATCTGTTATTGAGCTCTACCCTTGA

>TaOMT8 (TraesCS5D02G488900)

ATGGGTTCTATCTCCGACGACGAGGCGGCTTGCATCTACGCGATGCAGCTGGCAGCGGCAGCTGTCTTGCCGATG
ACGCTCAAGAACGCCATCGAGCTGGGCATGATCGAGATCCTCGTTGGCGCTGGCGGGAAGATGCTTTCGCCGTCT
GAGGTGGCCGCCCAGCTGCCGTCGACGGCCAACGCGGAAGCACCGGCCATCGTCGACCGCATGCTGCGGCTGCTG
GCATCGCACAACGTTGTGTCGTGCGAGGTGGAGGAAGGTAAGGACGGCCGCCTCGCACGCCGGTACGGCACCGCG
CCGGTGTGCAAGTGGCTCAGCCCCGACGAGGACGGCGTGTCCTTGGCCCCGATGGTCCTCTTGAACGACAAGGTC
ATGCTGGAGAGCTGGTACCATTTGAAGGACACTGTCCTTAACGGTGGCCTGCCATTCGAAAAAGCGTACGGAATG
ACAGCGTTTGATTACCAAGGAACGGACCCGCGCTTTAATCGCGTCTTCAATGAGGCCATGAAGACCCACTCCATG
ATCATCACCAAGAAGCTCCTGGAGTTCTACAAGGGCTTCGATGGTGTGGGCACCCTCGTGGACGTCGGTGGCGGT
GTGGGCGCCACCATCCACACCATCCTTTGCAGGTACCCAAGCATCGAGGGGGTCAACTTCGACCTCCCTCACGTC
ATCTCCGAGGCACCATCCTTTGCAGGTGTGCACCACATAGGTGGTGACATGTTCAAGAAGGTTCCCTCTGGGGAC
GCAATCCTCATGAAGTGGATACTCCACGACTGGAACGATGAGCATTGCACAACGCTGCTGAGCAACTGCTACGAT
GCACTACCCGCGCATGGCAAGGTGGTCATAGTGGAATACATCCTGCCAGTGAAGCCCGACGAAACGGCCACAGCA
CAGAGGTCCTTCGAAGCCGACATGATTATGCTCACACACACACCCGGCGGCAAGGAGAGGTACCTGAGGGAGTAC
GAAGAGCTCGCCAGGAGCGCCGGGTTTGCCAGCGTCAAGGCCACCTACGTCTACAGCAACATATGGGTCATTGAG
TTGAACAAATAG

>TaCPS-D2 (truncated)* (TraesCS2D02G029600)

ATGATTAGCAAATCCCCATCCTATCCGGAGGTAGATGTGGGTGAATGGAAAGCTGATGAGTACAGACAACGCACG
GATGAACCGAGCGAGATGCGACAGATGATCGATGCCATAAGAACGGCCCTGGCGTCATTGGGTGATGATGAGACT
TCAATGAGTGTTTCAGCCTACGACACAGCGTTGGTCGCCCTTGTCAAGAACCTTGACGGAGGCGATGGACCGCAA
TTTCCCTCATGCATCGACTGGATTGTTCAAAATCAACTACCAGATGGTTCGTGGGGTGATCCGGCTTTCTTCATG
GTCCAAGATCGGATGATCAGCACCCTGGCATGTGTCGTGGCAGTGAAGTCTTGGAACATTGACAGCGACAACTTG
TGTGACAGGGGTGTGTTATTTATCAAAGAAAATATGAGCAGGTTGGTAGAAGAGGAACAGGACTGGATGCCATGC
GGCTTTGAGATTAACTTCCCAGCACTCCTAGAGAAGGCCAAGGACCTGGACTTGGACATCCCTTATGATGACCCT
GTTTTGGAAGAGATATATGCCAAGAGGAATCTGAAGCTCTCTAAGATACCTCTGGATGTGCTGCACGCCATACCA
ACGACCCTACTCTTCAGCGTCGAGGGAATGGTAGATTTACCATTGGACTGGGAAACGCTACTCAGATTGCGGTGT
CCGGATGGCTCCTTCCATTCCTCACCTGCTGCCACAGCTGCTGCCCTCAGTCACACCGGTGACAAGGAATGCCAT
GCGTTCCTGGATAGGCTCATCCAAAAGTTCGAGGGGGGAGTGCCATGTAGCCACTCAATGGACACTTTTGAGCAA
TTATGGGTGGTCGATCGGCTGATGCGTCTGGGGATATCAAGGCACTTCACAAGTGAAATTCAGCAGTGCTTAGAG
TTTATTTACAGGCGCTGGACTCAAAAGGGACTGGCTCACAACATGCACTGCCCAATCCCGGATATTGATGACACA
GCCATGGGTTTCCGTCTTCTCCGTCAGCATGGCTACGACGTCACTCCATCTGTGTTTAAGCACTTTGAGAAGGAT
GGCAAGTTCGTCTGCTTCCCAATGGAGACTAACCACGCTTCTGTTACCCCAATGCACAACACTTACCGTGCATCT
CAGTTCATGTTTCCTGGTGACGACGACGTCCTGGCTAGGGCCGGCCGCTATTGCCGTGAATTCCTTCAAGAGAGA
CAATCCTCCAACAAGCTGTACGACAAGTGGATCATCACCAAGGACCTGCCGGGCGAGGTCGGGTACACGCTAAAC
TTCCCATGGAAAGCAAGTTTGCCGCGAATTGAAACAAGGATGTATCTGGATCAGTACGGTGGCAACAACGATGTT
TGGATTGCCAAGGTCCTTTACAGAATGAATCTTGTGAGCAACGACCTGTACCTTAAAATGGCTAAGGCTGATTTC
AGAGAATATCAAAGACTCTCCCGAATTGAATGGAATGGCCTCAGAAAGTGGTATTTCAGGAACCATCTGCAGAGG
TATGGTGCGACTCCAAAGAGCGCGCTGAAGGCTTACTTCTTAGCTTCAGCAAATATCTTCGAACCCGGCCGAGCA
GCAGAACGTCTAGCATGGGCTCGCATGGCGGTGCTTGCTGAGGCGGTTACAACTCACTTCCGACACATTGGGGGT
CCATGCTATTCAACAGAGAATCTTGAAGAGCTTATCGACCTTGTTTCGTTTGATGATGTTTCCGGCGGCCTTCGT
GAAGCGTGGAAGCAGTGGCTTATGGCATGGACTGCAAAGGAGAGCCATGGCTCCGTTGATGGAGATACAGCATTG
TTGTTCGTCCGCACCATCGAGATCTGCTCAGGAAGGATTGTTTCAGCCGAGCAGAAACCGAACCTTTGGGATTAT
TCTCAGCTCGAGCAGCTCACCTCTTCCATCTGCCACAAGCTTGCCACAATAGGTCTTTCTCAGAATGAGGCAAGT
ATGGAGAACACAGAGGACTTACACCGGCAAGTGGATTTGGAGATGCAAGAACTATCTGGGCGTGTTCATCAGGGT
TGCCATGGCATCAGTAGAGAGACTAGACAGACATTTCTCAACGTGGTGAAAAGCTTCTATTACTCGGCTCATTGC
TCACATGAAACAGTTGATAGCCACATCGCAAAGGTCATATTTCAGGATGTCATTTAG

>TaKSL-D1 (truncated)* (TraesCS2D02G030100)

ATGGCAATGTTGGCAGAAGAAACCATAGGACCGCGCAGTGATGTGGAACGGGATGCTAGAATACGAAAGCATCTC
AAAAACCCTGAACTCTCGCCGTCTGCGTATGACACGGCATGGGTGGCTATGGTGCCATTGCCGGACTTCGATCCG
CAGGCTCCATGCTTCCCTCAGTGTGTTGAATGGATATTGCAAAATCAACACTCTAGTGGGTCTTGGGGAATCAAT
GAATTTGGCTTATTAGCCAACAAGGATATTATGTTATCCACATTGGCCTGTATCATTGCACTTCATAAGTGGAAC
GTTGGCTCCGACCACATAAGGAGAGGATTAGAATTTATTGGAAGGAATTTCTCCACTGTCATGGATGATCAAATT
GTTTCTCCAGTAGGCTTCAATCTCATTTTCCCTGGTATGCTTAACCATGCTTTCGGGATGGGTTTGATAATTCCA
GTCACAGAAGCTGATATCAATGGGATACTTCACCTCCGTGAGATGGAGCTGACAAGATTAAGTGAGGAGAAATCG
TGTGGGAAAGATGCATATTTGGCCTATGTTGCTGAAGAAGGGTTAGTAAACCTGCTGGACTACAATCAAGTGATG
AAGTTCCAGCGAAAGAATGGGTCGTTGTTCAACTCTCCTGCCGCAACTGCTGCTGCATTAGTGCACTACTATGAT
AATAAAGCTCTCCAGTACCTCGACTCCATTGTCAGTATATTTGGTGGTGCAGTACCAACAGCGTACCCACAGAAT
ATATATTATCAGCTCTCAATGGTGGATATGCTCGAAAAGATCGGAATATCTCGCCATTTTTCGAGTGACACAAAC
AGCATCCTGGACAAGGCATACATTTCCTGGTTACAGAGAGACGAGGAGATCATGCAAGATGTAGAAACATGTGCA

ATGGCGTTTCGCCTTTTACGGATGAATGGTTATGATGTGTCGTCAGATGACTTGTCCCATGTTGCTGAAGCCTCC
ACTTTCCATAGCTCACTTGAAGGATATTTAAATCATACAAAATCTTTATTGGAGTTATACAAGGCTTCAAAAGTA
TGTTTGTCAGAAAATGAATTGATCCTGGAGAACATAAGCAACTGGTCAGGCCACTTATTGGCAGAGAAATTGCGC
TGTGATGGGACACAAAGAATGCCAATTTTTGGAGAGGTAGAATATACTCTTAAATTTCCCTTTTATGCGACAGTA
GAACCTCTAGACCATAAGAGGAACATTGAACATTTTGATTCTAGGGTTACTCAGCAGCTAAAGAGAAAAAACATG
CCATGTCATGTCAATCAAGATCTTCTAGATTTTGCCGTTGAAGATTTCAGTTTTTCTCAATCTATATACCAGGAT
GAACTCTGCCACCTCGAGAGTTGGGAGAAAGAAAACAGGCTGGAACAGCTTAAATTTCTACGCAAGGGGAGTCTG
ATAAATTGTTATCTCTCTGCTGCTGCCACCCTATCCACTCATGAACTCTCTGATGCTCGCATTGCATGTGCGAAA
ACTATTGCGCTCGTACTTGTTACTGATGACTTCTTTGATGTTGGAGCATCGAAAGAAGAACAAGAAAACCTCATA
GCATTAGTAGAGAAGTGGGATCACCATGACGAAGTTGAGTTCTGCTCTGAGCAAGTAGAAATAGTATTTTCTGCT
TTTTATAGTACAGTTAAGCACATTGGAGAAATGGCTTCCGCAGTGCAAAAGCGCGATGTTACAAAACACCTGAAT
GAAACATGGCTACATTACTTGAGGTCTGCAGCGACTGAGGCGGAATGGCAACGGAATCAATATGTGCCAACAGTT
GAGGAATACATGATAGAAGCGGTTAACTCATTCGCAGAGGGGCCCATTATGCTAACATCACTATATTTTGTCCAA
CAAAAACTCGAGGAGTACATAATCAAAGACCCGGAGTACCATGAGTTGCTTAGAATAAAGGGCAACTGTGGCCGT
CTCCTGAATGATACTAGGGGCTTCGAGAGGGAGTCCAGTGAGGGAAAACTGAACATCATCTCACTGCTTGTTCTT
CAGAGTGGAGGTTCCATGTCCATAGAAGCTGCTCAAGAGGCGGTACAGGAGTCTATAGCCTCATGTCGGAGAGAC
CTGCTAAGGATGGTTGTTAGAGAAGACCGTGTAGTTCCTAGGGTATGCAAGGAGGTGTTCTGGAGGTTTTGCAGG
ACAGTTCACTTGTTCTACTGTCAGACTGACGGATTTTCCTCGCCCAAGGAAATGCTCCGCACGATGAACGCAATA
TTCCGAGAGCCACTCAAACTCCAACATACCAGTCCTTTGGATGCTCAGTCAGAACAATAA

>TcGGPPS(truncated)*

ATGGCTTCCTATCAAGAATGCAATAGTATGAGGAGTTGTTTTAAATTGACACCTTTTAAAAGTTTTCATGGAGTG
AATTTCAATGTTCCCTCACTGGGTGCTGCTAATTGTGAGATTATGGGTCACCTGAAACTTGGGTCATTGCCATAT
AAACAATGTTCGGTGTCATCTAAATCCACAAAAACAATGGCCCAGTTGGTTGATTTGGCTGAAACAGAGAAGGCG
GAGGGAAAGGATATTGAATTTGATTTCAACGAGTATATGAAGTCCAAGGCTGTGGCAGTGGATGCGGCACTGGAT
AAGGCAATCCCACTTGAATATCCTGAAAAAATACATGAATCAATGAGGTATTCACTTCTAGCAGGAGGTAAGCGC
GTCAGGCCTGCTCTGTGCATTGCAGCATGTGAGCTTGTAGGAGGGAGTCAGGACCTTGCCATGCCAACTGCCTGT
GCAATGGAGATGATTCATACCATGTCTCTGATTCATGATGACTTGCCGTGCATGGATAATGATGATTTCAGAAGA
GGGAAGCCAACAAATCACAAGGTCTTTGGAGAGGACACTGCTGTTCTTGCAGGGGACGCCCTGCTTTCATTTGCA
TTTGAGCATATTGCTGTGGCTACAAGCAAGACTGTGCCTAGTGATAGGACTTTAAGGGTGATATCTGAATTGGGT
AAGACAATAGGCTCTCAAGGGCTTGTAGGGGGGCAGGTGGTTGATATTACATCCGAGGGGGATGCTAATGTGGAC
CTGAAAACCCTGGAATGGATTCATATACACAAGACTGCTGTGCTCTTGGAATGTTCAGTTGTGAGTGGAGGGATC
CTTGGTGGTGCTACAGAGGACGAGATTGCGAGAATTCGGCGGTACGCCCGGTGTGTGGGGCTTCTGTTTCAGGTT
GTGGATGACATACTTGATGTCACTAAATCTTCTGAAGAATTGGGAAAGACTGCAGGAAAGGATTTGCTTACTGAT
AAGGCTACTTATCCCAAGTTGATGGGCCTGGAGAAAGCAAAAGAATTTGCCGCTGAATTGGCGACGAGAGCCAAG
GAAGAGCTGTCATCCTTTGATCAGATAAAGGCTGCACCTTTGTTGGGTCTTGCAGATTACATTGCATTCAGGCAA
AACTGA

>TaOSC(5D)* (TraesCS5D02G011800)

ATGTGGAGGCTAAAGATCGCGGAGGGCAGCGGCAACCCACTGCTGCGCACCACCAACGGCCACGTCGGCCGGCAG
GTATGGGAGTTCGATCCTGCCGCGGATGACCCCGACGAGGTCGCCGCCGTTGATGCCGCCCGCCGCGACTTCACC
AGCCGCCGGCACCAGATGAAGAACAGCTCCGACCTCCTCATGCGCATGCAGCTCTCTAAGTCAAACAATCTCAAG
ATGAACCTTCCTCCCATCAAGCTGGAACAACATGGGTATCACACAGAAGAAGACGATGTCTTGATATCTTTGAGG
AGGGCAATCAACCGATGTTCCACTCTCCAGGCACATGATGGCCATTGGCCGGGGGATTACGCTGGCACTTTATTT
CTCTTGCCCAGCTTGATTGTAGCACTGCATGTGACTGGAGCACTAACTACTGTGTTGACATCTGAACATCAGAAA
GAGATTCGGCGGTATCTCTACAATCATCAGAATGAAGACGGGGGTTGGGGACTGCATATCGAGGGAACAAGCACA
ATGTTCTGTACTGTCATGAACTATGTTGCTTTGAGATTGCTTGGGGAGGGGCTGGATTCATGCGGAGCCATGCTA
CAAGCTCGGAGTTGGATCTTGGATCATGGAGGAGCAACTTTAACACCATCGTGGGGAAAATTCTTTCTTTCGGTC
CTTGGGGTATATGACTGGTCCGGGAACAACCCATTGCCACCAGAACTATGGATGATGCCTTATTTCTTGCCTATT
CATCCAGGGCGCATGTGGTGTAATTGTCGGTTGGTATATTTGCCCATGTCATACCTATACGGGAAGAGGTTTGTG
GGTGCTACTACATCGATTGTACTAGACTTGAGGAAAGAGCTCTACAATGTCCCGTATAATGAGATTGACTGGGAC
AAGGCTCGCAATGGGTGTGCAAAGGAAGATCTCTATTATCCCCATTCACCACTACAGAATATTGTATGGGCCACT
CTCAAGAAAATTGGTGAACCGGTTCTTATGCACTGGCCTGGCAGCAATTTGCGAAAGAAAGCTCTGGACATTGTC
ATGAAACATATAAAGTATGAGGATGAAACAACTCAATATATATGCATTGCTCCTCTCAATAAGATGTTGAATATG
ATTTGTTGCTGGGTAGAAGATCCAAAGTCAGAGGCATTCAAACTCCATATTGAAAGAGTATATGATTTCTTATGG
GTTGCAGAAGACGGCATGAAGATGAAGTCTTACAATGGCAGCCAGCTATGGGATACAGCTTTGACAGTTCAGGCT
ATATTTGCTACCGGCCTCACAAAAGAGTTTGGCCATACAATTAAAGTCGCTCATGACTATATCAAACATTCCCAG
ATTCGTGCGGACTGCCCTGGAGACCAAAGCAAGTGGTATCGCCATCTATCAAAAGGTGGATGGACACATTCAACT
GCTGACCAAGGGTGGCCTGTATCAGATTGTACTGCAGAAGCATTTAAGGTTTTGTTGTTGTTAACTAAGGTTCCT
GAACTGGTTGATGAGCCTATAGAAACAAGCAGACTTGATGATGCAGTCAACATTCTACTGTCTTTGATGAATGAC

GATGGCAGTTTTGGTGCATATGAGCTAACAAGATCTTATGAATGGCTTGAGTTGTTGAATCCTTCTGAGAGCTTT
GGCGGCATAATGATCGAGTATCCGTATGTTGAGTGTACATCATCAGTAATTCAGGGTCTGGTGTTATTTAGAGAA
ATGTACCCCAAACACATTCGCTTGGAAGAGATAGACAGTTGTATCCAAAAGGCTGCAGACTACATTGAGAGCATC
CAATGGAGTGATGGATCATGGTATGGTTGTTGGGCTATCTGTTTCACATATGGTACTTGGTTTGGGGTGAGAGGA
TTGGTTGCTGCAGGGAGGACGTATGTAAATAGCCTAGCAATCAGAAAAGCGTGTGAATTTCTTTTGTCCAAGGAG
CTTCTCCCATCTGGAGGGTGGGGAGAAAGTTACCTTTCCAGTCAAGACAAGGTTTACACCAATCTTGAAGGAAAC
AGGGCTCATGCAGTTAACACTAGTTGGGCCATGTTGGCCCTAATTGATGCCGGGCAGGGTCAAAGAGATCCTGCA
TGTCTACATAGAGCAGCCAAGATTCTGATGAACTTCCAATCAGAGGATGGAGAGTTCCCTCAGCAAGACATTATT
GGAGCTACCAACCACAACCTTATGCTCACTTATGCCCAGTTCAGGAACATATTCCCTATCTGGGCTCTTGGAGAG
TACTACCAGAGAGTGCTTCCGGTAGCTTAA

>TaHSD(5D)* (TraesCS5D02G011900)

ATGGATATGTGCCCGGTTGGCGGCTGCGACCGCCGGGAACCACGTCGTTTGTGCGCGGTGACCGGTGGGCGGGGA
TTCATGGCGAGTCACCTGGTGACGGCGCTGCTCGGCTCGGGAGATTGGTGTGTGCGGATCACCGACCTTGGCCCC
CAGGCTGCCCTGTCTCCTGCCGAGAGTGATGGACTCCTGGGCGCTGCCCTCCGCGATGGCCGCGCCGCCTACTTC
TCTGTCGACGTCTGCGAATTAGCCCAGCTTACAAAAGCTTTGGAAGGGGTAGATACTGTTTTCCACACTGCTGCG
GCGGATCATACCAACAACAACTTCCAACTTCATTACAAGGTTAATGTCGAGGGTACAAGGAATGTCATCGAGGCT
TGTAACACATGCAAGGTTAAAACACTCATATATACTAGTTCCAGTGGAGTTGTATTCGATGGAGTTCATGGCCTC
TTTGGCGTAGACGAATCTACACCGTATCCAGATAAGTTTCCCGACGCATACACAGAGACAAAGGCAGAAGCAGAA
AAGATGGTGATAAAGTCCAACGGAAGAAATGAGCTTCTCACTTGCTGTATACGTCCTGGCAGCATTTTTGGTCCT
GGAGACACGATAGTGCCAATTTTAGTATCTTATGGAGGAATGATGATCATTGTTGGTGATGGCAAGAATTGTGAT
GATTTTGTATATGTTGAGAATGTAGCGCATGGTCATGTATGTGCTGAGAAAACTCTTTCTACCATAGACGGTGCA
AAGAGAAGTGGAGGCAAAGCCTATTTTATAACAAATATGGAGCCTGTAAATATGTGGGACTTTGTTTATATGATT
TTGGAAGAACTTGGATACAAAAGCCGATTCAGATTAAGAATACCTTCATATTTTCTCAAGCCAATAACATATCTG
GTAGATTGGAGCTACCATAATATATTCTCTCACTATGGAATGCGTCAACCTGGCATGCTAACTTCCGCAAGCATT
AAGTATGCGACGCTGAATAGAACATTCAACTGCAACAATGCTGCTCAACAACTTGGTTACAAACCAATAGTGTCA
CTCAAGGAGGGAGTAAAGATGACTACTGATTTCTACAGGCGATTAAGAGCATGA

>TaCYP51H35 (TraesCS5D02G012300)

ATGGACTTAGCAAGTCTCACCGCAGTGTGGTGGGCTGTGGCTCTTCTTTTCATCACCGTGTTAGTCACCAAGATT
TCAAGAGCAAGAGTCACCACCGTTGATCTACATCGTACAACAGGTCAACTTCCTCCCCTGGTGAATGGAGTTGCT
CTCCTAAAACTATTACCTACCCTTTTGAACAAGGGTCTACCGGCAATGGTGAATGATCTATATGTCAAATATGGC
AGTGTATTCATGGTAAGTTCATTTGGAGTGAAGGTAACACTCTTGATCGGGCCAGAGGTGACGGCTCATTTCTTC
CAAGGTCTGGAGTCGGAGATTAGCCACGGTAATCTGTTTGAGTTCACTGTGCCCATGTTTGGTGAAGCAGTGGGT
TATGGCAGAGATACCGCCACCCGAACTGAACAAATGCGCTTCCATATCGAAGCACTGAGACCATCAAGGTTGAGG
AGCCATGTTTATCCCATGCTTCAAGAAGTGGAGGGTTACTTTGCAAAATGGGGAGAGGAAGGCATTGTTGATCTA
AAGCTTGAGTTCGAGAAGTTACTCATGTTGATCTCAAGCAGATGTTTGCTCGGAAAAGAGGTTCGGGAGAACATG
TTTGATGAAGTCTACACACTTTTTCACGAGATTGAAGATAATGGTGTGACCTTGATTAGCTTCTTGTTCCCATAT
CTCCCAACTCCAGCAAATCGGAAGCGAGATAAAGCGCGCATCAGGCTAACACAAATCCTTTCTGATGTCGTCGAT
TCCCGTAAGAACTCCGGCAGAGTTGAGGATGATACATTGCAGAAATTGATCGACTCTAAGTACAAAGATGGTCGC
CCTACAACGGTAGAAGAGGTAGTCGGGCTAATCATTGGCCTGTTGTTTGCTGGAAAACACACCAGCTCTCACACT
AGTACTTGGACTGCAGCTTGTTTACTCAGCCATCCAACTTTCTTGAGAGCTGCCATTGAGGAGCAACAACAAATC
AGTAGCAAATACAAGGACATGGGGCTAGACTACAATGCATTTATAGAGATGGATACACTACATAGTTGCATCAAG
GAGGCGCTAAGGAAGCACCCTCCAACACCAATGCTGGTCCGCAGGGCACATAAGCAATTCATGGTGAAGACGAAA
GAGGGCAAAGAATATGACATCCCGCAAGACCACATCGTAGCAACTCCTACTATAGTGACTAATAACATCTCTTAT
ATCTATAAGGACCCTCAGGTATATGATCCATGCCGGTTTGGCCCCGAAAGAAGAGAGGACAAAGTTGGTGGCAAG
TTCTCTTACACGTCATTTAGTGGTGGAAGACACATTTGCACCGGGGAGGCCTATGCTTACATGCAACTTAAAGTG
ATATGGAGCCATTTGCTGAGGAACTTTGAGCTCGAATTGATCTCCCCTTTCCCCAAGACAGACTGGGGCAAGTTC
TTGCCAGAGCCACAAGGAAAACTACTCGTAAAATATAAGAGGAATGGCATTTTGTAG

>TaCYP51H37 (TraesCS5D02G012000)

ATGGAGATGGCAAGTAGCGCCAAGTGGTTTGCTGTGGCTCTTGTTTTCATCACTGTGATCCTACTCAGGGTCATA
AGAGGAAGGAGGATGGCCGCTGCTCCAGCCAGTGAGAAACCACCTCCACCTGTGGTGAATGGTTTTGCGTTACTA
GGACTTCTACCTAGGCTTCTTACAATGGACCTTCGAACTAAGATAAATTGCCTGCACGATAAGTATGGCAGTGTG
TTCACAGTTAGTCCTTTTGGACTAACTAATGTAACCTTTCTGATCGGCCCGGAGGTCCAATCCCATTTCTTTCAA
GGGTTGGAGTCAGAGATTAACCATGGCAATCTTCTTGAGTTCCTTGTGCCGATGTTTGGACAAGAGATAGGTCAA
GGTGTCGATGCAGCCACTCGGACTGAGCAGTCCCGCTTTTATCTCGATGCATTAAAGCAATCCAAGTTGAGGAGA
CATCTTGATCCGATGCTTCAAGAGGTCGAGAGCTACTTCGGCAAATGGGGCCTGGAAGGAGTAGTTGATTTAAAA

CATGAGTTTGAAGAGTTGCTCATGTTGATATCAAGCCGGTGTCTACTAGGAAAAGAGGTTCGGGAGAAGATGTTT
GATGAGTTCTATAAACTTTTTCGTGATGTCGAAAATGGAGTGAACATGATCAGTGTCTTCTTCCCATATATTCCA
ATTCCTGCAAACCGTAGGCGTGACAAAGCACGTCTTAAGCTCATCGAATTACTTTCTGGAACTGTGAGGTCACGT
AAGAGCTCCCCCTCAGTGGAGGAAGATGTGCTACAAAGATTGATAGATTCCAAGTATAAAGACGGCCGCTCCACA
ACGGAAACAGAGGTAACCGGGATGATCATTGCATTGATCTTTGGTGGAAAGCACACAAGTTCCCTCGCTAGTACC
TGGACCGGAGCTTGCCTACTCACTCATCCAAAGTTCTTAGCGGCTGCATCCCAGGAGCAAAAGGAAATCATGATG
AAATACAAGGACAAAATAGAATATGATGCCTTGTTAGAGATGAATACCCTTCATAATTGTATCAAAGAGGCACTT
AGGCTGAACCCACCAACAACAATGTTGGTTCGCAAGGCACTTAAACATTTCACAGTGCGGACAAGACAAGGCCAA
GAGTATGGCATTCCCAAGGGGCATACTCTTGCAAGTCCCATAATACAAAACAATTACTTGCCTTACATTTATAAG
GACCCCGAGGTATATGATCCAGATCGGTTTGGTCCCGTAAGGCAGGAGGACATAGTTGGCGGCAAGTTTTCTTAT
ACATCCTTTGGTGGTGGAAGGCATTTTTGCAGTGGGGATGCCTATGCTTACATGCAAGTTAAAGTTATATGGAGC
CACTTGCTCAACAACTTTGATCTCAAATTATTATCTCCTTATCCCAAGACTGACTGGAGTAAGTTAATCCCAGAG
CCTCAAGGGAATATGATGGTCAGCTACAAGAGACGCCAGCTGCTAGGCTAG

>TaCYP51H13_5A (TraesCS5A02G004600)

ATGGACTTGACAAGTCTCACTACGATGTGGTGGGCCATAGCTCTTCTTTCCATTACTGTATTGGCCACCAAGATC
ACAAGAGCAAGAATCAAAAACATTGATCCACAGCGTACAACAGGCCAACTTCCACCCATGGTGAATGGACTTGCT
CTCCTAGGATTATTACCTACCCTTCTGAAGAAGGGTCTACCACCTATGTTCAATTATCTATATGTTAACTATGGC
AGTGTCTTCACTGTAAGTTGCTTTGGAGTGATCAAGGTAACACTCTTGATCGGGCCAGAGGCGACGACTCATTTC
TTCCAAGGTTTGGAGTCAGAGATTAGCCATGGTAATCTGCTTGAGTTCACAGTACCCATGTTTGGCAAGGCAGTT
GGTTATGGTAGAGATACCGCCACCCGAATGGAACAAATGCGCTTCCACAGTGAAGCACTGAGGGCATCCAGGTTA
AGGAGCCATGTTTCTCCAATGCTTCAAGAAGTGGAGGATTTCTTTGCAAAGTGGGGAGAGGAAGGCGTAGTTGAT
CTAAAGCTTGAGTTCGAGCAGCTACTTATGTTGATATCAGGCCGATGTTTGCTTGGAAAAGAGGTCCGGGAGAAT
ATGTTTGATGAAGTCTACACACTTTTTCGCGAGATCGAAAAGGGTGTGACCTTGATCAGCTTCTTGTTCCCATAT
CTCCCAACTCCAGCAAACCGGCAGCGTGATAGGGCACGTATAAGGCTAACAGAGATCCTCTCTAATGTTGTCGAG
TCCCGTAAGAGCTCCGGGCGAGTGGAGGAGGACACGCTGCAGAAACTGATTGACTCCAAGTACAAAGATGGTCGC
CCTACAACAGTAGAAGAGGTAGCCGGGTTGATCATTGGCTTGCTGTTTGCTGGAAAACACACCAGCTCTCATACT
AGTACTTGGACTGGAGCTTGTCTTCTCAGCCATCCAGCCTTCTTAAGAGCTGCCATTGAGGAGCAACATCAAGTC
GCCAAGAAATACAAGGACGGACCAGAATACAATGCCTTCTTAGAGATGGATACGCTGCATAATTGCATCAAGGAG
GCGCTGCGGAAGAACCCTGCAACACCACTGCTGGTTCGCAGGGTCCATAAGCGCTTCACGGTGCAGACGAAAGAA
GGCCAACAATATGACATCCCGCAGGACTACATCGTAGCAACTCCTACTATAGTAAACAATTATATCCCTTACATT
TATACGGACCCTCAGGTGTATGATCCAAATCGGTTTGGCCCAGAAAGAAGAGAGGACAAAGTGGGTGGCAAGTTC
TCTTACACATCATTTGGTGGTGGAAGGCACGTTTGCACTGGAGAGGCTTATGCTTACATGCAACTCAAAGTGATA
TGGAGCCATTTGCTGAGGAACTTTGATCTCGAATTGGTCTCTCCCTTCCCTAACACAGACTGGAGCAAGTTCTTG
CCAGATCCAAAGGAAAAATTACTCGTGAGATATACGAGAAATGGAATTTAA

>TaCYP51H13P (TraesCS5D02G012100/200)

ATGGACTTGACAAGTCTCACTGCCATGTGGTGGGCCATAGCTCTTCTTTTCATTACTGTATTGGCCACCAAGATC
ACAAGAGCAAGAATCACCAACATTGATCCACAGCGTACAACAGGCCAACTTCCACCCATGGTGAATGGACTTGCT
CTCCTAGGATTATTACCTACCCTTCTGAAGAAGGGTCTACCACCTATGGTCAATTATCTATATGTTAACTATGGC
AGTGTCTTCACTGTAAGTTGCTTTGGAGTGATCAAGGTAACACTCTTGATCGGGCCAGAGGCGACGACTCATTTC
TTCCAAGGTTTGGAGTCAGAGATTAGCCATGGTAATCTGCTTGAGTTCACAGTGCCCATGTTTGGCAAGGCAGTT
GGTTATGGTAGAGATACCGCCACCCGAATGGAACAAATGCGCTTCCACAGTGAAGCACTGAGGGCATCCAGGTTA
AGGAGCCATGTTTCTCCAATGCTTCAAGAAGTGGAGGATTTCTTTGCAAAGTGGGGAGAGGAAGGCGTAGTTGAT
CTAAAGCTTGAGTTCTAGCAGCTACTTATGTTGATATCAGGCTGATGTTTGCTTGGAAAAGAGGTCCGGGAGAAT
ATGTTTGATGAAGTCTACACACTTTTTCGCGAGATCGAAAAGGGTGTGACCTTGATCAGCTTCTTGTTCCCATAT
CTCCCAACTCCAGCAAACTGGCAGCGTGATAGGGCGCGTATAAGGCTAACAGAGATCCTCTCTAATGTTGTCGAA
TCCCGTAAGAGCTCCGGGCGAGTGGAGGAGGACACGCTGCAGAAACTGATTGACTCCAAGTACAAAGATGGTCGC
CCTACAACAGTAGAAGAGGTAGTCGGGTTGATCATTGGCTTGCTGTTTGCTGGAAAACACACCAGCTCTCATACT
AGTACTTGGACTGGAGCTTGTCTTCTCAGCCATCCAGCCTTCTTAAGAGCTGCCATTGAGGAGCAACATCAAGTC
GCCAAGAAATACAAGGACGGACTAGAATACAATGCCTTCTTAGAGATGGATACGCTGCATAATTGCATCAAGGAG
GCGCTGCGGAAGAACCCTGCGACACCACTGCTGGTTCGCAGGGTCCATAAGCGCTTCACGGTGCAGACGAAAGAA
GGCCAACAATATGACATCCAGCAGGACTACATCGTAGCAACTCCTACTATAGTAAACAACAATATCCCTTACATT
TATACGGACCCTCAGGTGTATGATCCAAATCGGTTTGGCCCCGAAAGAAGAGAGGACAAAGTGGGTGGCAAGTTC
TCTTACACATCATTTGGTGGTGGAAGGCACGTTTGCACCGGAGAGGCTTATGCTTACATGCAACTCAAAGTGATA
TGGAGCCATTTGCTGAGGAACTTTGATCTCGAATTGGTCTCTCCCTTCCCTAACACAGACTGGAGCAAGTTCTTG
CCAGATCCACAGGGAAAATTACTCGTGAGATATACGAGAAATGGAATTTAA

>AtCYP51H35 (AET5Gv20012700)

ATGGACTTAGCAAGTCTCACCGCAGTGTGGTGGGCTGTGGCTCTTCTTTTCATCACCGTGTTAGTCACCAAGATT
TCAAGAGCAAGAGTCACCACCGTTGATCTACATCGTACAACAGGTCAACTTCCTCCCCTGGTGAATGGAGTTGCT
CTCCTAAAACTATTACCTACCCTTTTGAACAAGGGTCTACCGGCAATGGTGAATGATCTATATGTCAAATATGGC
AGTGTATTCATGGTAAGTTCATTTGGAGTGAAGGTAACACTCTTGATCGGGCCAGAGGTGACGGCTCATTTCTTC
CAAGGTCTGGAGTCGGAGATTAGCCACGGTAATCTGTTTGAGTTCACTGTGCCCATGTTTGGTGAAGCAGTGGGT
TATGGCAGAGATACCGCCACCCGAACTGAACAAATGCGCTTCCATATCGAAGCACTGAGACCATCAAGGTTGAGG
AGCCATGTTTATCCCATGCTTCAAGAAGTGGAGGGTTACTTTGCAAAATGGGGAGAGGAAGGCATTGTTGATCTA
AAGCTTGAGTTCGAGAAGTTACTCATGTTGATCTCAAGCAGATGTTTGCTCGGAAAAGAGGTTCGGGAGAACATG
TTTGATGAAGTCTACACACTTTTTCACGAGATTGAAGATAATGGTGTGACCTTGATTAGCTTCTTGTTCCCATAT
CTCCCAACTCCAGCAAACCGGAAGCGAGATAAAGCACGCATCAGGCTAACACAAATCCTTTCTGATGTCGTCGAT
TCCCGTAAGAACTCCGGCAGAGTTGAGGATGATACATTGCAGAAATTGATCGACTCTAAGTACAAAGATGGTCGC
CCTACAACGGTAGAAGAGGTAGTCGGGCTAATCATTGGCCTGTTGTTTGCTGGAAAACACACCAGCTCTCACACT
AGTACTTGGACTGCAGCTTGTTTACTCAGCCATCCAACTTTCTTGAGAGCTGCCATTGAGGAGCAACAACAAATC
AGTAGCAAATACAAGGACATGGGGCTAGACTACAATGCATTTATAGAGATGGATACACTACATAGATGCATCAAG
GAGGCGCTAAGGAAGCATCCTCCAACACCAATGCTGGTCCGCAGGGCACATAAGCAATTCATGGTGAAGACGAAA
GAGGGCAAAGAATATGACATCCCGCAAGACCACATCGTAGCAACTCCTACTATAGTGACTAATAACATCTCTTAT
ATCTATAAGGACCCTCAGGTATATGATCCATGCCGGTTTGGCCCCGAAAGAAGAGAGGACAAAGTTGGTGGCAAG
TTCTCTTACACGTCATTTAGTGGTGGAAGACACATTTGCACCGGGGAGGCCTATGCTTACATGCAACTTAAAGTG
ATATGGAGCCATTTGCTGAGGAACTTTGAGCTCGAATTGATCTCCCCTTTCCCCAAGACAGACTGGAGCAAGTTC
TTGCCAGAGCCACAAGGAAAACTACTCGTAAAATATAAGAGGAATGGCATTTTGTAG

>AsOSC1* (AS01G014480)

ATGTGGAGGCTGAAGATCGCGGAGGGCGGCGGCAACCCGCTGCTGCGCACCACCAACGGCCACGTTGGGCGGCAG
GTGTGGGAGTTCGATCCTGCCGCCTCCGACGATGACCCCGACGAGATCGCGGCCGTAGACGCCGCGCGCCGCGAG
TTCACCGGCCGCCGCCATCAGATTAAGAACAGCTCCGACCTCCTCATGCGCATGCAGTTCTCCAAGACAAACGCT
CTCAAGATTAACCTTCCTCATGTCAAGCTGGCAGAGCATGAATATGCAACAGAAGAGGATGTCCTCATATCTTTG
AAGAGGGCAATTAGCCGCTATTCCACTCTCCAGGCACATGATGGGCACTGGCCGGGGGATTATGCAGGCACTTTA
TTTCTCTTGCCCAGCTTGATTGTAGCATTGCACGTGACTGGCGCACTGACTACAGTCTTGACATCTGAACATCAG
AAAGAGATCCGGCGATATCTCTACAATCATCAGAATGAAGATGGAGGCTGGGGACTGCACATTGAGGGAACAAGC
ACAATGTTCTGTTCTGTCATGAACTATGTTGCTTTGAGATTGCTTGGGGAAGGGCTGGATTCAGGTGGAGCCATG
CTGCAAGCTCGTAGTTGGATCTTGGACCATGGAGGAGCAACTTTAACACCATCTTGGGGCAAATTCTTTCTATCG
GTCCTCGGGGTATATGACTGGTCTGGAAACAACCCATTGCCACCAGAAATATGGATGCTTCCCTATTTCTTGCCT
ATTCATCCAGGACGTATGTGGTGTAATTGCCGACTAGTATATTTGCCAATGTCGTACTTATATGGAAGAGGTTT
GTGGGTTCTATGACATCAATTGTCCTAGACTTGAGGAAAGAGCTGTACAATGTCCCATACGATGAGATTGACTGG
GACAAGGCTCGCAATGGATGTGCCAAGGAAGATCTCTATTATCCCCATTCACCGCTGCAGGATATCGTGTGGGCT
ACTCTCAAGAAAATTGGTGAACCAGTTCTCATGCACTGGCCTGGCAGCAAATTGCGGGAGAAAGCTCTGGACATT
GTGATGAAACATATTGAGTACGAGGATGAAACAACACAATATATATGCATTGCTCCTATGAATAAGATGTTGAAT
ATGATTTGCCGTTGGGTAGAAGATCCGAACTCAGTGGCATTCAAACTCCATATTGAAAGAGTATATGATTTCTTA
TGGGTCGCTGAAGACGGCATGAAGATGAAGTCTTATAATGGCAGCCAGCTGTGGGATACAGCTTTGACTGTTCAA
GCTATATTTGCTGCCGGACTCACAGAAGAGTTTGGCCAGACAGTTAAACTTGCGCATGACTATATCAAGCGTTCC
CAGATTCGTATGGACTGTCCTGGAGATCAAAGTAAATGGTATCGTCATATATCTAAAGGTGGATGGACACATTCA
ACAGCTGACGAAGGGTGGCCTGTATCAGACTGTACTGCGGAAGCACTGAAGGTTTTATTGCTGTTAACTAAGTTT
CCTCCTGAATTGGTTGGTGAATCTATAGAAGCAAGCAGGCTTGATGATACAGTCAACATTTTGCTGTCTTTGATG
AACGACGATGGTAGTTTTGGTGCATATGAGCTAACAAGATCTTATGAATGGCTTGAGTTGCTCAATCCTTCTGAG
AGCTTTGGGGGCATAATGATCGAGTATCCATATGTTGAGTGTACGTCATCAGTAATTCAAGGTCTTGTGTTATTC
AGAGAAACCTACCGTAAACATTATCGCAGGGAAGAGATAGACCATTGTATCCGAAAGGCTGCAGACTACATTGAG
AGCATCCAACGAAGTGATGGATCATGGTATGGCTGTTGGGCAATTTGTTTCACCTATGGTACATGGTTCGGGGTG
AGAGGATTGGTTGCTGCCGGTAGGACGTACAAAAACAGCCAAGCAATCAGAAAAGCGTGTGAATTTCTGCTATCT
AAGGAGCTTCTTCCGTCAGGAGGATGGGGAGAAAGCTACCTCTCTAGTCAAGACAAGGTTTACACCAATCTTGAA
GGCAGCAAGGCTCATGCAGTTAACACCAGTTGGGCCATGTTGGCCCTGATTGATGCTGGGCAGGGTAAAAGGGAC
CCTGCATGTCTACATAGAGCAGCCAAGGTTCTGATCAATTTCCAATCCGTGGATGGAGAGTTCCCTCAGCAAGAC
ATTATCGGAGCTACCAACCACAACCTCATGCTCACTTACGCCCAGTTCAGGAACATCTTCCCTATCTGGGCTCTT
GGGGAGTATTATCGCAGAGTGTTGATAGCACAAAGGCTCTAA

>CYP51H73* (AS01_002578_0375348)

ATGGACATGACAAGTGGCGCCCTGTGGTTCACCTTGGCTCTTGTTTTCATCACTCTCCTACTGAGGCTTATACAA
CGAAGGATTGCCGTTGCTCCAACCAGTAGAAACCCACTTCCACCAGTGGTGAATTGTTTTGCTTTGCTGCGAATT
TTACCTAGCCTTTTTGGAAAGGATCTTCCAACGAAGATAAATTTTCTATATAAGAAGTATGGCAGTGTGTTCACA
ATTAGTCTATTTGGACGTCATGTAACCATTCTGATCGGACCGGAGGTATCAGCTCATTTCTTTCAAGGGTTGGAT
TCAGATATTAGCCATGGCAACTTTCTTGAGTTTACTGTACCCATGTTTGGGCAAGAGGTTGCTTATGGCGTTGAT

ACGGCAACTCGGAATGAGCAGTCTCGTTTCTATCTTGAAGCATTAAGGCAATCAAAGTTGAGGGGACATTTCCAT
CCCATGCTTCAAGAGGTGGAGGAATACTTTGGCAAATGGGGCGAGGAAGGCATTGTTGATTTAAAACATGAATTT
GAAGAGATACTCATGTTGATCTCAAGCCGTTGTCTACTAGGAAAAGAGGTCCGGGAGAATATGTTTGAGGATTTC
TATGCACTGTTCCGTGATATCGAAAATGGAGTGAACTTGGCCAGTGTCCTTTTCCCATACATCCCAATTCCAGCA
AACCGCAGGCGTGACAAGGCACGTATGAAGCTAATAGAAATATTCTCTCAAATTATGAGATCACGTGAGAGCTCT
AGCAGGGTTGAGGAGGACGTGCTACAGAAATTGATGGATTCTAAGTATAAAGACGGGCGCTCCACAACCAAAACA
GAGGTAACAGGACTGATAATTGGGTTGATCTTTGCTGGAAAGCACACAACCTCTCAAGCTATAACCTGGACTGGG
GCTGACCTACTTAGCCATGAAAAATTCTTAGCAGCTGCATCCGAGGAGCAAAAGGAAATCATGATGAAATACAAG
GGCAAAATAGAATATGATGCCTTGCTAGAGATGAGTACCCTTCATAGGTGTATCAAAGAGGCACTTCGGATGCAC
CCACCAGGACCAATGTTGATTCGCAAGGCACATAAAAATTTCACGGTCCGAACAAAAGAAGGACAAGAGTATGAT
ATTCCGAAAGGGCATAGCATTGCAAGTCCTATAGTGCAAAACAATAGGATGCCTTACATTTATAAGGATCCTCAT
TTGTATGACCCAGATAGGTTTGGTCCGGCAAGACGAGAGGACGTAGTTGGCGGCAAGTTCTCTTACACATCTTTT
GGCGCTGGAAGGCATTCTTGCGTTGGGGAGGCCTATTCTTATGTGCAAATTAAAGTCATATGGAGCCATCTACTG
AACAACTTTGATCTCAAATTATTGTCCAGTTATCCTAAAACTAATTGGAATAAGCTAATCCCAGAGCCTCAAGGA
AGTATGATGGTCAGCTACAAGAGACGTCGGCTACCGGGCTAG

>BdOSC2 (Bradi3g22802)

ATGTGGAAGCTAAAGATCGCAGAGGGGTGCGGCAACCCGCTGCTGCGCACCACCAACGATCACACTGGCAGGCAG
GTATGGGAGTTCGACCCTGACGACTCCGTCGACGAGTCTCCAGCCGTGGAAGCCGCCCGCCGCGAGTTCTCCAGC
CGCCGGAATCAGACCAAGAACAGTTCTGACCTCGTCATGCGCATGCAGTTCTCCAAGTCTAACCGTCTCAAGATG
AACCTTCCTGCCATCAAGCTGGATGAGCGTGCATTCGCCACGGAAGAGGATGTCTTGATATCTTTGAAGAGGGCA
ATCGGCCGCTATTCCGCACTCCAGGCACATGATGGGCACTGGCCGGGAGATTACGCAGGAACTCTGTTTCTCTTG
CCTAGCTTGATTGTAGCATTGCATGTGACTGAATCACTCAATACTGTGTTGTCATCTGAACATCAGAAAGAGATG
CGGAGATATCTCTACAATCATCAGAATGAAGATGGGGGCTGGGGACTGCACATTGAGGGAACAAGCACAATGTTC
TGTTCTGTCATGAACTACGTCGCCTTGAGATTGCTTGGGGAAGGGCTGGATTCGTGTGGAGCCATGCTGCAAGCT
AGGAGTTGGATCTTGGACCATGGGGGGCAACTTTAACACCATCTTGGGGGAAATTCTTCCTTTCGGTCCTTGGG
GTATATGAATGGTCTGGCAACAACCCATTGCCACCAGAAATATGGATGATGCCTTATTTCTTGCCTATTCATCCA
GGGCGCATGTGGTGTAATTGCCGCCTGGTATATTTGCCCATGTCATACCTATATGGAAAGAGGTTCGTGGGAGCT
ATTACATCATCAGTTGTACTAGACTTGAGGAAAGAACTTTACATTGTGCCATACGATGAGATTGACTGGGACAAG
GCTCGCAATGGCTGTGCAAAGGAAGATCTCTATTATCCTCATTCACCGCTGCAGGATGTGGTATGGGCTACTCTG
AAGAAAGTTGGCGAGCCACTTCTTATGCACTGGCCTGGCAGCAAATTGCGGCAGAGAGCTCTGGACGTTGTCATG
GAACATATCCGTTACGAGGATGAAACAACTCAATATATCTGCATTGCTCCTCTGAACAAGATGTTGAATATGATT
TGTCGCTGGGTAGAAGATCCAAACTCAGAGGCATTCAAGCTCCATATTGAAAGAGTCTACGATTTCTTATGGGTC
GCTGAAGACGGCATGAAAATGAAGTCATACAATGGCAGCCAGTTGTGGGATACGGCTTTAACAGTCCAGGCTATT
TTTGCTACCGGCCTCACAGAAGAGTTTGGCCCCACAATTAAACTAGCCCATGACTACATCAAACGTTCCCAGATT
CGTGTTGACTGTCCTGGAGACCAGAGCAAGTGGTATCGCCATATATCCAAAGGTGGCTGGACGCACTCAACTGCT
GACCAAGGGTGGCCTGTATCAGATTGTACTGCAGAAGCATTGAAGGTTCTGTTGCTGTTAACTAAGGTTCCTCCT
GAATTGGCTGGTGAGCCTATGGAAGCCAGCAGGCTTGATGATAGTGTCAACCTTCTGTTATCTCTGATGAACGAG
GATGGCAGTTTTGGTGCATATGAGCTGACAAGATCTTATGAGTGGCTTGAGTTGCTCAATCCTTCTGAGAGCTTT
GGGGGAATAATGATCGAGTATCCGTATGTTGAGTGTACATCATCAGTAATTCAGGGGCTGGTGTTATTCAGAGAA
ACATACCCTAAACACTGTCGCAGGGAAGAGATAGACAACTGTATCCGGAAGGCTGCAGATTACACAGAGAGCATC
CAACGTGCCGATGGATCATGGTACGGCTGTTGGGCTATCTGTTTCACCTATGGTACATGGTTTGGGGTGAGAGGA
TTGATTGCTGCCGGTAGGACTTACAAGAACAGCCAATCAATCAGAAAGGCGTGCGAATTTTTGCTGTCCAAGGAG
CTCCCATCAGGAGGATGGGGGGAAAGCTACCTCTCCAGTCAAGATAAGGTTTACACCAATCTTGAAGGCAACAAG
GCTCATGCAGTTAACACCAGTTGGGCCATGTTGGCCCTGATTGATGCTGGACAGGGTAAAAGAGATCCTGCATGT
CTGCATAGAGCAGCCAAGGTTCTGATCAACTTCCAAATGGAGGATGGAGAGTTCCCTCAGCAAGACATCATCGGA
GCTACCAACCACAACCTCATGCTCACTTACGCCCAGTTCAGGAACATCTTTCCGATCTGGGCTCTTGGAGAGTAC
CATCGCAGAGTTCTGCAAGCACAAAAGGCATAA

>BdCYP51H14 (Bradi3g22820)

ATGTTCATGACAAGTAGCGCCATGTGGTTCACCGTGTCTCTTGTTTTCATCACCATCGTGCTCATCAGGATCATC
AGAGGGAAGGCAGCTGGTGGTGTTGATCCAACTTCTGGGAAACAACACCCGCCTGTGGTGAATGGCTTTGCTTTA
CTAGGACTTTTACCTAGGCTTTTGAACAAGGATCTTCCAACTAAGATAAATTGTCTATATAGTAAGTATGGTAGC
GTGTTCACGGTAAGTCTGTTTGGACGGAACGTAACCTTCTTGATCGGGCCTGAGGTCTCAGCTCATTTCTTTCAA
GGATTGGACTCAGAGATTAGCCATGGCAATCTTCTTGAGTTCACTGTCCCCATGTTTGGCCAAGAGGTCGCTCAT
GGTGTCGATACAGCCACTCGGAATGAGCAGGCCCGGTTCTATCTTGACGCATTAAAGCAATCAAAGTTGAGAAGC
CATTTTGATCCCATGCTTCAAGAGGTAGAGGAATACTTTAGCAAATGGGGGCAGGAAGGCATAGTTGATCTGAAG
CACGAATTTGAGCAGTTACTTATGTTGATCTCAAGCCGATGTCTACTCGGCAAAGAGGTCCGAGAGAAGATGTTT
GACGAGTTCTATACCCTGTTTCGTGACATTGAAAACGGTGTGAACTTGGTCAGTGTCTTCTTCCCATATATCCCA
ATCCCAGCAAACCGGAGACGCGACAGAGCACGCGTGAAGCTCATTGAGCTGCTCTCTGAAATCGTGAGGTCACGT

AAGAGCTCAGGCAGAGTGGAGGAGGACGCCCTGCAGAAACTGATAGATTCCAAGTACAAAGACGGCCGCTCCACA
AGCGAAACAGAAGTCACCGGGATGATAATTGGACTGATCTTTGGGGGAAAACACACAAGCTCTCATGCCACTACC
TGGACCGGAGCCTGCCTACTCAGCCATGAGAAGTTCTTAGCAGCCTCATCCGAAGAGCAAAAGCAAATCATGAAG
AAATACAAGGGGAGATTAGAATATGAAGCTCTGTTAGAGATGAATACACTCCATAGTTGCATCAAGGAGGCGCTT
CGGATGCACCCGCCATCGCCGATGCTGGTTCGCAAGGCAAATAAGCATTTCACTGTGAAGACAAGAGAGGGCCAG
GAGTATGATATTCCAAAAGGGAATACTATTGCAAGCCCTATAGTACAAAACAATATCATGCCATACATTTATAAG
GACCCTCACTTGTATGATCCAGATAGGTTTGGACCCGCGAGGCAGGAGGACGTAGCTGGTGGGAAATTCTCTTAC
ACGTCTTTTGGAGGTGGAAGGCATGTCTGCGTTGGCGAGGGCTATGCTTACATGCAGATTAAAATCATATGGAGC
CATTTGCTCAAGAACTTTGAGCTCGAGTTGCTGTCTCCTTTTCCCAACACCGATTGGAATAAGTTAATCCCAGAG
CCTCAAGGGAATATGATGGTTAGCTATAAGAGACATCGGCTGTTGGGCTAG

>BdCYP51H15 (Bradi3g22840)

ATGGACTTGGCAAGCACAGCCGCCGTGGGATGGGCGCTCGCTCTTCTTCTCATCACTGCTTTAGCTACCAAGATA
GCAAGAGCAAGAATCACCATTGTCGATCCACCGCGTACGACGACGACGACGACTCAACCTCCACCCGTGGTGAAT
GGCATCGCTCTCCTAAGACTGTTACCCACCCTTTCCAAGAAGGGTCTGCCGGCCATGGTGAACGATCTGTATCTG
AAGTTCGGCAGCGTGTTCATGGTAAGCGTGTTTGGGATCAAGGTAACGATCTTGATCGGGCCAGAGGTGACCCCT
CACTTCTTCCAAGGTCTGGAGTCGGAGATTAGCCACGGTAATCTGCTTGAGTTCACCGTGCCCATGTTTGGCGAA
GCGGTTGGTTACGGCAGAGATGCCGCCACCCGAATTGAACAGACCCGCTTCCACATCGAGGCGCTGAGGCCATCC
AGGTTGAAGAACCACGTCTATCCCATGATTCAAGAGGTCGAGGACTATTTCGCAAAATGGGGAAACGAGGGCATA
GTGGACCTAAAGCTTGAGTTCGAGCGGTTACTCATGTTGATCTCAAGCCGAGTTTTACTCGGAAAAGAAGTTAGA
GAAACCATGTTTGATGAGGTCTACACACTGTTTCGTGAAATCGAAAACGGTGTGACCGTGATCAGTTTCTTATTT
CCATATCTCCCGATTCCAGCAAACCACCGGCGAGATAGAGCGCGCACCAGGCTCACAGAAATACTCTCCGATGTC
GTCAAGTCGCGTAAGAACTCAGGCAGGGTTGAGGAGGACACACTCCAAAAATTGATAGACTCCAAGTATAAAGAT
GGCCGCTCCACAACCGTACCAGAGGTTGTCGGGTTGATCATCGGTCTCCTATTTGCTGGAAAACATACCAGCTCT
CACACCAGCACCTGGACTGGAGCCTGTTTGCTTAGCCACCCAATGTTCTTAAGAGCTGCCATTGAGGAGCAAAAA
CAAATCGGTAGGAAATATAAGGATGGGCTCGGCTACGATGCTATGTTAGAGATGGATACCCTGCATAGTTGCATC
AAGGAGGCGCTAAGGATGCACCCTCCAACGCCAATGCTGGTTCGCAAGGCACATAAGCACTTTACGGTGCAGACG
AAACAAGGCAAACAATATGACATCCCAGAAGGGCACACCGTAGCGAGTCCTATTGTAGTCAACAATAATATCCCC
TATATTTATAAAGACCCAGAGGTTTATGATCCAGACCGGTTTGGTCCCAAAAGACAAGAGGACAAGGTAGGTGGC
AAGTTCTCTTATACGTCCTTCAGTGGTGGAAGGCACACTTGCACTGGTGAGGCCTATGCTTACATGCAAATTAAG
GTGATATGGAGTCATTTGCTAAGTAACTTTGAACTCAAATTGATCTCACCTTTCCCCAAGACAGATTGGAGCAAA
TTCTTACCAGAGCCAGAAGGAACATTGTTTGTAAGATATAAGAGAAATGGCTGCCGTGCTAGCACTTAG

>BdCYP51H16 (Bradi3g22850)

ATGGAATTTACAAGTGGCGACGTGTGGTTCGCCGTAGCTGTTCTCTTAATTGCTACGGTCGTCACCAA
GATTGCAATTGCAAGAGCTACCGTTGATCCAACGTGCACAAGACCACTTCCACCTGTGGTGAAGGGCG
TTGCTCTCCTTGGACTTCTACACGCCCTTGTTACCAAGGATCTACCAACTGTGATACATGATCTGTAT
GAAAAATTTGGCAGTGTGTTCACAGTAAGCTTGCTTGGACAAAAGGTGACCTTCTTGGTCGGGCCAGA
GGTGTCGGCCCATTTCTTCAAAGGGTTGGAGTCAGAAATTAACATCGGCAATCTGCTGAATTTCACCG
TGCCCATATTTGGTCAAGAGGTGGGTTACGGCGTGGATCTCGCAACTCGGAATGAGCAGGCCCGATTC
TGCCTTGATGCACTGAAGCCGTCCAAGTTGAGGAGCCATGTTGACCCCATGCTTCAAGAAGTAGAGGA
GTACTTTGCAAAGTGGGGACAACATGGGATAGTTGATTTAAAACATGAGTTTGAGGTGTTACTCATGT
TGATCTCAAGCCGGGTTCTACTCGGTAAAGAGGTCCGGGAGAAGATGTTCGATGAGTTCTGCACATTA
TTTGGTCAGATCGAAAACGGGGTGAACCTTGTCAGTGTCTTCTTCCCATACATCCCAATTCCATCAAA
CCAGAGACGCGACAGAGCACGCGTCAAGCTCACACAGATACTATCTGAAGTTGTGAGTTCGCGCAAGA
GCTCCGGCCGAGTTGAAGAGGATACGCTGCAGAGACTCATCGATTCCAGGTATAAAGACGGCCGATCC
ACAACGGAAGCAGAGATCACCGGGATGATAATTGGCATGCTCTTCGCTGGGAAACACACGAGCTCTCA
CACTACTATCTGGACTGGAGCATGCCTCCTTAACAGTCCCAAATTCCTGGCTGCTGCTGTCGAGGAGC
AAAAGGAACTCGTTGTGAAATACAAAGACCAGATCGACTACAATACCTTGG:CAGAGATGGATGCCCCT
CCATTGCTGCATCAAAGAGGCACTCCGGATGCACCCTCCGTCGCCCGTGCTGGTCCGCAAGGCACAGA
AGCAGTTCACCGTGAAGACGAAAGAAGGTAACGAGTATGACATCCCAAGGGGGCATACCATTGCCAGC
CCTACGATAGTTAACAACAATATGCATTACATTTATAAGGAGCCCCAAGTGTATGATCCGGATCGTTT
TGGCCCTGGAAGAGAGGAGGACATAGTTGGTGGCAAGTTCTCTTACACGTCATTTGGCAGCGGAAGAC
ATGCTTGCATTGGCGAGTCATATGCTTACATGCAAATTAAATTGATATGGAGCCATTTGCTGAGGAAT
TTTGAGCTTGAACTGGTTTCTCCTTTCCCGAGGACAGACTGGAACAAGATTGTGCCAGGACCTCAAGG
GAAGGTCATGGTAAGTTATAAGAGATCGAGGATGTCAGCCTAG

>BdACT* (Bradi3g22830)

```
ATGGCTGCAATTGTTGCTAGAGCTGCTAACGGTGGACTAAGAGGTCGACAGGAGACATCCAGGGGTTC
ACTAGTACGCCGCAGAAGGGACTATGCTGTTGTTGTTGCTGTTGTCACTGGTGCTGGTGCCCCACCAC
CACAAGAAGGCGCATTAGAACGTCCAGCATGGAGTGGTGAAACTCCAGTTTCACGTCTAGTCGGTGCT
CTAATAGCTTTTAAACCCTTGTATGCATTGATGAAATTGGCTAGCAGAGAAGTTATCATTAGAACTGC
CGAGAAGGCTAATATTCCATGGCGAGAAATGACAAAGAAAGTTCTTGAAAGTGATGTTTATGAGGTCT
TTGAAAGAATTCGTGACCCAAATTTGGTGTATCCCGATTACTACTTATCTCCTTTCCATGCTTACGAT
GAGGGTAATCTCAGCTGGCTTGCGGCTGCAGAAGCTGAAGCAGCCACTTTATCTATAGCTAAGAGAGC
TATCCCGGAAGCCACATCAATCGAAGAGGCTAATCAAATAGTAAGAGGGAATTGGATGAACGCAATCG
AAGAACACCATCTTAAATACTCAGGTCATAGGCAAATTAACGATATACTTGATATAGGTTGCAGTGTA
GGCGTTTCTACACGTTACTTGGCCGATAGATTTCCATCTGCGAAGGCAGTCGGTTTGGACCTTTCACC
CTATTTCTTGGCTGTTGCCGCCCAGAAAGAGGAGAAGATGAGCAGGCAACACCCAATTAGATGGGTCC
ACGCTAACGGGGAAGAAACTGGTCTGAGTCCTGACCTATTCGATGTTGTTAGTTTAGCCTACGTTTGT
CATGAATGCCCCGCCAGGGCTATCCACGGTCTGGTTAAAGAGGCTTTTCGGTTGTTAAGACCTGGTGG
TACTATAGCTCTCACAGACAACTCTCCGAAGAGTAAGGTTCTACAAGAGCTGAGCCCTGTTCTATTTA
CTCTGATGAAAAGTACGGAGCCATTCTTGGACGAATATTATATGTTAGATCTTGAAGAAGCTCTGTCT
CAGGCAGGGTTCGTAAACGTTCACAGTATATTTACTGATCCGCGGCACAGGACGGTCACCGCTACAGT
TCCTTCAACAAAGAAGAAGTTCCTTAGCAGCTAA
```

**Supplementary Methods.** Purification of compounds from large-scale *N. benthamiana* agroinfiltration.

**Purification of 19-hydroxy-isoarborinol**

160 6-weeks old *N. benthamiana* plants were manually infiltrated with *Agrobacterium tumefaciens* strains containing pEAQ-HT-DEST1 vectors[1] expressing TaIAS and TaIAH. For infiltration, agrobacteria strains were mixed 1:1 in MMA buffer (10 mM $MgCl_2$, 10 mM MES/KOH pH5.6, 150 μM acetosyringone) to a final concentration of $O.D._{600}$ 0.2 per each strain. Infiltrated leaves were harvested 6 days post infiltration and lyophilized. 111 gr of powdered leaf material was extracted twice with 15 ml/gr ethanol, using a Buchi Speed extractor E916 (100ºC, 100 bar). The extract was saponified by addition of 35 ml water and 15 gr KOH per 150 ml of extract, and incubation for 2 h at 65ºC. Following addition of 50 ml water per 150 ml of extract, the saponification reaction was extracted three times with 100 ml hexane. Solvent was removed by rotary evaporation and residual was applied to a flash chromatography column (35mm diameter) filled with 100 ml of silica (LC60A35-70 µm). 19-hydroxy-isoarborinol was eluted from the column with 1:1 hexane:ethyl acetate. Fractions containing the product were pooled and evaporated. Decolorization with addition of a minimal amount of activated charcoal was followed by recrystallization in an ethanol-$H_2O$ mix, to give a final yield of 20 mg purified compound.

**Purification of ellarinacin**

100 *N. benthamiana* plants were infiltrated by vacuum[2] with *A. tumefaciens* GV3101 strains containing vectors for expression of TaIAS, TaIAH, TaHID, TaHIO and oat tHMGR. For infiltration, agrobacteria strains were mixed in MMA buffer to a final concentration of $O.D._{600}$ 0.2 per each strain. Infiltrated leaves were harvested 8 days post infiltration and lyophilized. 129 gr of powdered leaf material was extracted with ethyl acetate using a Buchi Speed extractor E916, and chlorophylls were removed from the extracts by addition of ion-exchange resin, following the protocol detailed in[2]. Three successive rounds of fractionation were performed on an Isolera Prime flash chromatography system (Biotage),

with hexane-ethyl acetate gradients specified in Supplementary Table 9, to yield 10.3 mg of purified compound.

**Purification of brachynacin**

100 *N. benthamiana* plants were infiltrated by vacuum with *A. tumefaciens* GV3101 strains containing vectors for expression of BdIAS, BdIAH, BdHIH, BdTIH, BdACT and oat tHMGR. For infiltration, agrobacteria strains were mixed in MMA buffer to a final concentration of $O.D._{600}$ 0.15 per each strain. Infiltrated leaves were harvested 9 days post infiltration and lyophilized. 69 gr of powdered leaf material was extracted with methanol using a Buchi Speed extractor E916, following the protocol detailed in[2]. Two successive rounds of fractionation were performed on an Isolera Prime flash chromatography system (Biotage), with hexane-ethyl acetate gradients specified in Supplementary Table 9. Product-containing fractions from flash chromatography were pooled and evaporated, and resulting product was dissolved in 1 ml 95% MeOH for semi-preparative HPLC purification on an Agilent 1290 Infinity (II) system. A Luna 5 µm C18(2) 100 Å, 250 x 10 mm LC column (Phenomenex) was used, with column oven temperature set at 25ºC. A mobile phase gradient was run at 3.5 ml/min with buffers A- 0.1% formic acid in $H_2O$, and B- 0.1% formic acid in acetonitrile, as follows: 20-60% B from 0-25 min, 60-100% B from 25 to 25.5 min, 100% B from 25.5 to 28 min, 100-20% B from 28 to 28.5 min, 20% B from 28.5 to 30 min. 0.5 ml of sample was injected in five 100 µl injections. Eluent was monitored by an Evaporative Light Scattering Detector (ELSD). Collected fractions containing brachynacin product were pooled and evaporated, to yield <2 mg of purified compound.

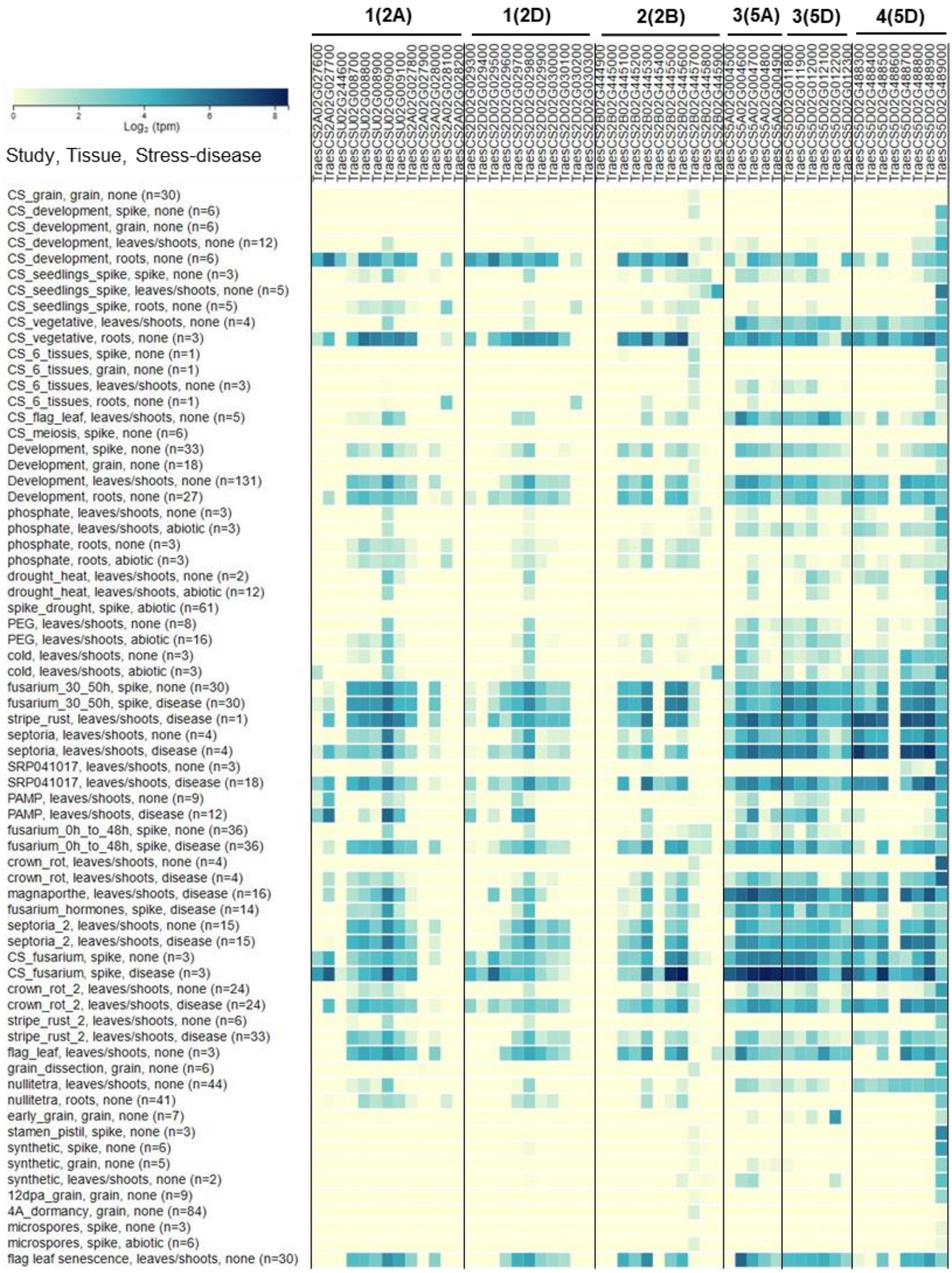


**Supplementary Fig. 1. Gene expression data of six wheat biosynthetic gene clusters**. Log2 of normalized values (transcripts per million) are shown, derived from RNA-seq data at http://www.wheat-expression.com[3,4].

**a**

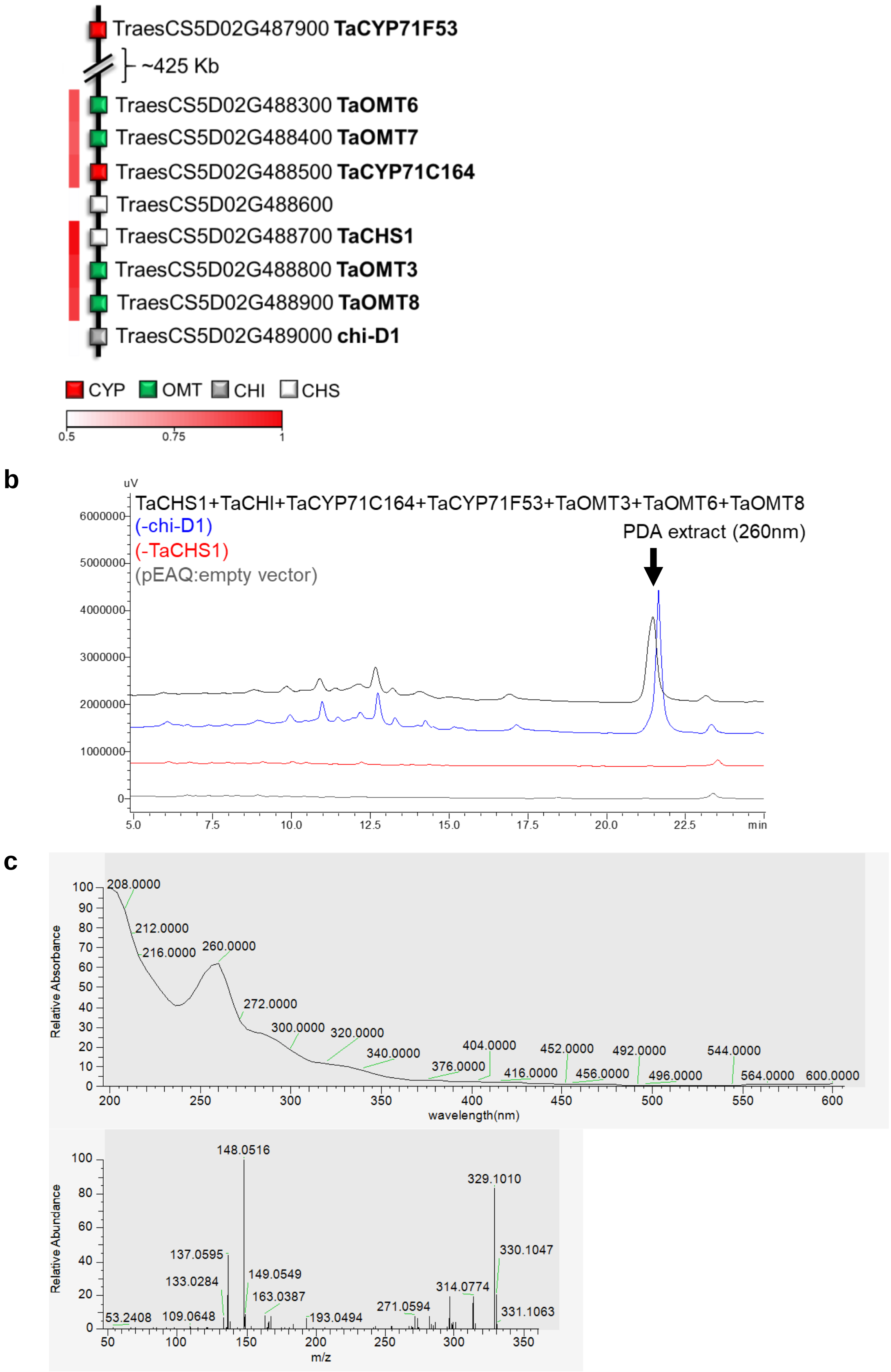


**Supplementary Fig. 2. a,** names and IWGSC IDs of wheat cluster 4(5D) genes. White to red color-coding denotes Pearson correlation (r) values for expression of each gene with TaCHS1. **b,** LC-PDA analysis of cluster 4(5D) genes expressed in *N. benthamiana*. The putative end-product of the pathway, marked with an arrow, is formed by expression of the complete cluster or with the absence of chi-D1, but not formed when TaCHS1 is not expressed. **c,** UV-VIS and ms$^2$ spectra of the [M+H=329.1] ion produced by cluster 4(5D).

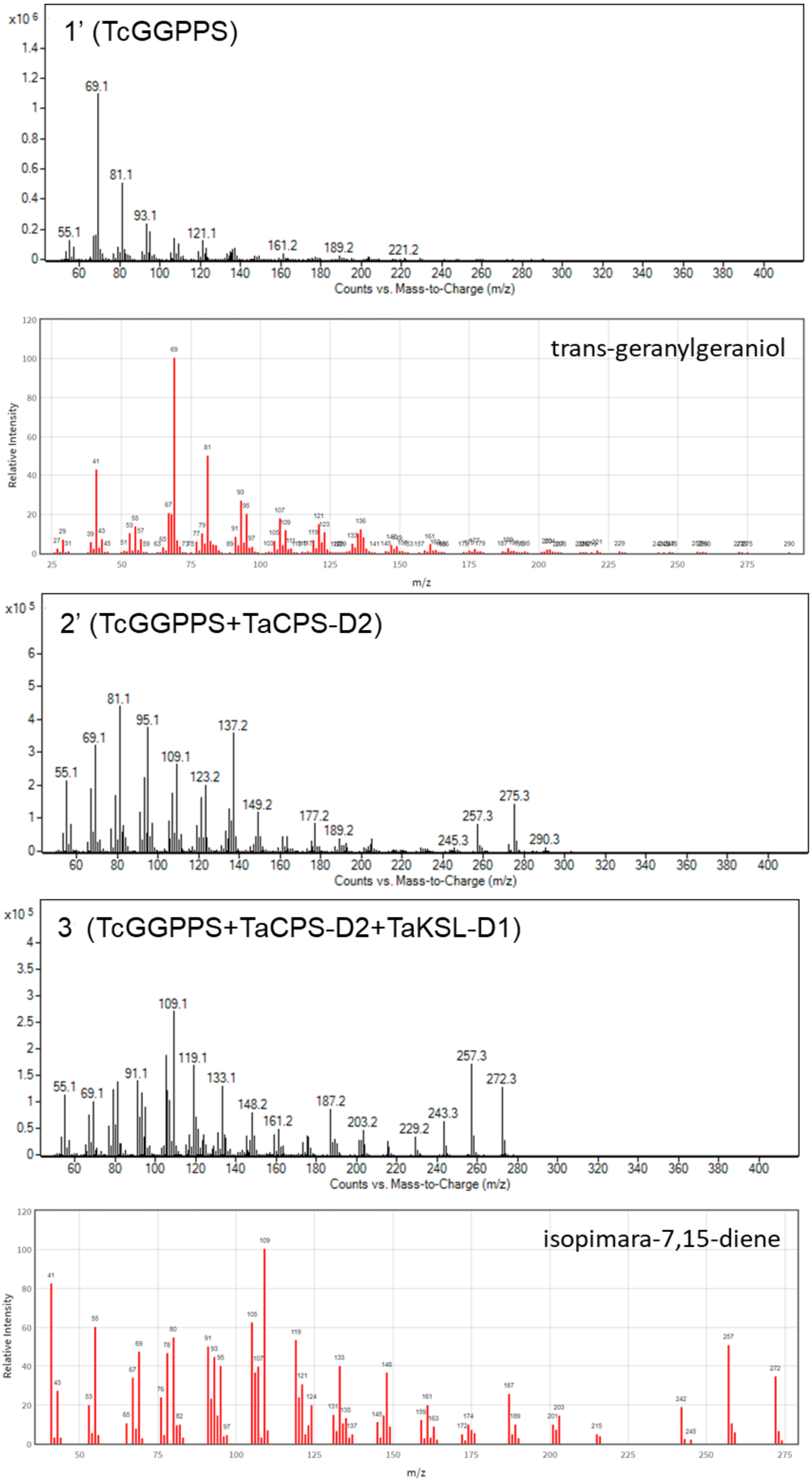


**Supplementary Fig. 3. Mass spectra of wheat TaKSL-D1 and TaCPS-D2 products in *N. benthamiana* expression**. TaKSL-D1 and TaCPS-D2 were transiently expressed together with *Taxus canadensis* GGPP synthase and oat tHMGR. Mass spectra are shown at retention times of peaks putatively identified as geranylgeraniol (1'), copalol (2') and isopimara-7,15-diene (3), based on comparison to NIST Chemistry WebBook (1', 3) (https://webbook.nist.gov/chemistry) and literature (2')[5]. tHMGR was included in all combinations of genes.

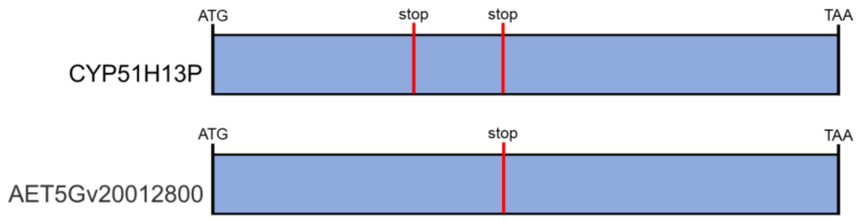


*T. aestivum* CYP51H13P coding sequence (manually corrected)

ATGGACTTGACAAGTCTCACTGCCATGTGGTGGGCCATAGCTCTTCTTTTCATTACTGTATTGGCCACCAAGATCACAAGAGCAAGAATCACCAACATTGATCCACAGCG
TACAACAGGCCAACTTCCACCCATGGTGAATGGACTTGCTCTCCTAGGATTATTACCTACCCTTCTGAAGAAGGGTCTACCACCTATGGTCAATTATCTATATGTTAACTAT
GGCAGTGTCTTCACTGTAAGTTGCTTTGGAGTGATCAAGGTAACACTCTTGATCGGGCCAGAGGCGACGACTCATTTCTTCCAAGGTTTGGAGTCAGAGATTAGCCAT
GGTAATCTGCTTGAGTTCACAGTGCCCATGTTTGGCAAGGCAGTTGGTTATGGTAGAGATACCGCCACCCGAATGGAACAAATGCGCTTCCACAGTGAAGCACTGAGG
GCATCCAGGTTAAGGAGCCATGTTTCTCCAATGCTTCAAGAAGTGGAGGATTTCTTTGCAAAGTGGGGAGAGGAAGGCGTAGTTGATCTAAAGCTTGAGTTC**TAG**CAG
CTACTTATGTTGATATCAGGC**TGA**TGTTTGCTTGGAAAAGAGGTCCGGGAGAATATGTTTGATGAAGTCTACACACTTTTTCGCGAGATCGAAAAGGGTGTGACCTTGAT
CAGCTTCTTGTTCCCATATCTCCCAACTCCAGCAAACTGGCAGCGTGATAGGGCGCGTATAAGGCTAACAGAGATCCTCTCTAATGTTGTCGAATCCCGTAAGAGCTCC
GGGCGAGTGGAGGAGGACACGCTGCAGAAACTGATTGACTCCAAGTACAAAGATGGTCGCCCTACAACAGTAGAAGAGGTAGTCGGGTTGATCATTGGCTTGCTGTT
TGCTGGAAAACACACCAGCTCTCATACTAGTACTTGGACTGGAGCTTGTCTTCTCAGCCATCCAGCCTTCTTAAGAGCTGCCATTGAGGAGCAACATCAAGTCGCCAAG
AAATACAAGGACGGACTAGAATACAATGCCTTCTTAGAGATGGATACGCTGCATAATTGCATCAAGGAGGCGCTGCGGAAGAACCCTGCGACACCACTGCTGGTTCGC
AGGGTCCATAAGCGCTTCACGGTGCAGACGAAAGAAGGCCAACAATATGACATCCAGCAGGACTACATCGTAGCAACTCCTACTATAGTAAACAACAATATCCCTTACAT
TTATACGGACCCTCAGGTGTATGATCCAAATCGGTTTGGCCCCGAAAGAAGAGAGGACAAAGTGGGTGGCAAGTTCTCTTACACATCATTTGGTGGTGGAAGGCACGT
TTGCACCGGAGAGGCTTATGCTTACATGCAACTCAAAGTGATATGGAGCCATTTGCTGAGGAACTTTGATCTCGAATTGGTCTCTCCCTTCCCTAACACAGACTGGAGC
AAGTTCTTGCCAGATCCACAGGGAAAATTACTCGTGAGATATACGAGAAATGGAATTTAA

*A. tauschii* AET5Gv20012800 coding sequence (manually corrected)

ATGGACTTGACAAGTCTCACTGCCATGTGGTGGGCCATAGCTCTTCTTTTCATTACTGTATTGGCCACCAAGATCACAAGAGCAAGAATCACCAACATTGATCCACAGCG
TACAACAGGCCAACTTCCACCCATGGTGAATGGACTTGCTCTCCTAGGATTATTACCTACCCTTCTGAAGAAGGGTCTACCACCTATGGTCAATTATCTATATGTTAACTAT
GGCAGTGTCTTCACTGTAAGTTGCTTTGGAGTGATCAAGGTAACACTCTTGATCGGGCCAGAGGCGACGACTCATTTCTTCCAAGGTTTGGAGTCAGAGATTAGCCAT
GGTAATCTGCTTGAGTTCACAGTGCCCATGTTTGGCAAGGCAGTTGGTTATGGTAGAGATACCGCCACCCGAATGGAACAAATGCGCTTCCACAGTGAAGCACTGAGG
GCATCCAGGTTAAGGAGCCATGTTTCTCCAATGCTTCAAGAAGTGGAGGATTTCTTTGCAAAGTGGGGAGAGAAAGGCGTAGTTGATCTAAAGCTTGAGTTCGAGCAG
CTACTTATGTTGATATCAGGC**TGA**TGTTTGCTTGGAAAAGAGGTCCGGGAGAATATGTTTGATGAAGTCTACACACTTTTTCGCGAGATCGAAAAGGGTGTGACCTTGAT
CAGCTTCTTGTTCCCATATCTCCCAACTCCAGCAAACTGGCAGCGTGATAGGGCGCGTATAAGGCTAACAGAGATCCTCTCTAATGTTGTCGAGTCCCGTAAGAGCTCC
GGGCGAGTGGAGGAGGACACGCTGCAGAAACTGATTGACTCCAAGTACAAAGATGGTCGCCCTACAACAGTAGAAGAGGTAGTCGGGTTGATCATTGGCTTGCTGTT
TGCTGGAAAACACACCAGCTCTCATACTAGTACTTGGACTGGAGCTCGTCTTCTCAGCCATCCAGCCTTCTTAAGAGCTGCCATTGAGGAGCAACATCAAGTCGCCAA
GAAATACAAGGACGGACTAGAATACAATGCCTTCTTAGAGATGGATACGCTGCATAATTGCATCAAGGAGGCGCTGCGGAAGAACCCTGCGACACCACTGCTGGTTCG
CAGGGTCCATAAGCGCTTCACGGTGCAGACGAAAGAAGGCCAACAATATGACATCCCGCAGGACTACATCGTAGCAACTCCTACTATAGTAAACAACAATATCCCTTACA
TTTATACGGACCCTCAGGTGTATGATCCAAATCGGTTTGGCCCCGAAAGAAGAGAGGACAAAGTGGGTGGCAAGTTCTCTTACACATCATTTGGTGGTGGAAGGCACG
TTTGCACCGGAGAGGCTTATGCTTACATGCAACTCAAAGTGATATGGAGCCATTTGCTGAGGAACTTTGATCTCGAATTGGTCTCTCCCTTCCCTAACACAGACTGGAG
CAAGTTCTTGCCAGATCCACAGGGAAAATTACTCGTGAGATATACGAGAAATGGAATTTAA

**Supplementary Fig. 4. Coding sequences of *T. aestivum TaCYP51H13P* and its *A. tauschii* ortholog, *AET5Gv20012800*.** Early stop codons are marked in red. TaCYP51H13P is the product of two IWGSC annotated genes, *TraesCS5D01G012100* and *TraesCS5D01G012200*. Manual annotation of these genes showed that they represent a single transcript with two premature stop codons and were therefore designated as a single pseudogene. The manual annotation was verified by cloning and sequencing of the full transcript from a cDNA library.

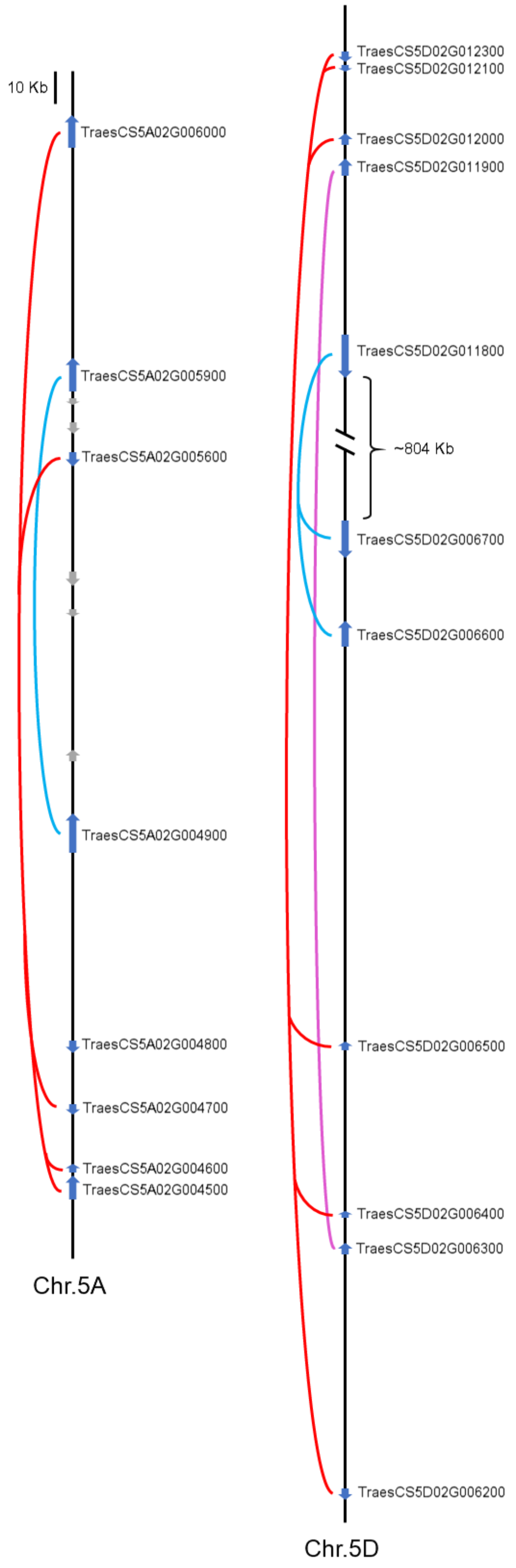


**Supplementary Fig. 5. Paralogs of the 3-5A and 3-5D cluster genes on wheat chromosomes 5A and 5D**. Blue lines connect paralog OSC genes, red lines CYP51 genes, and pink line HSD genes.

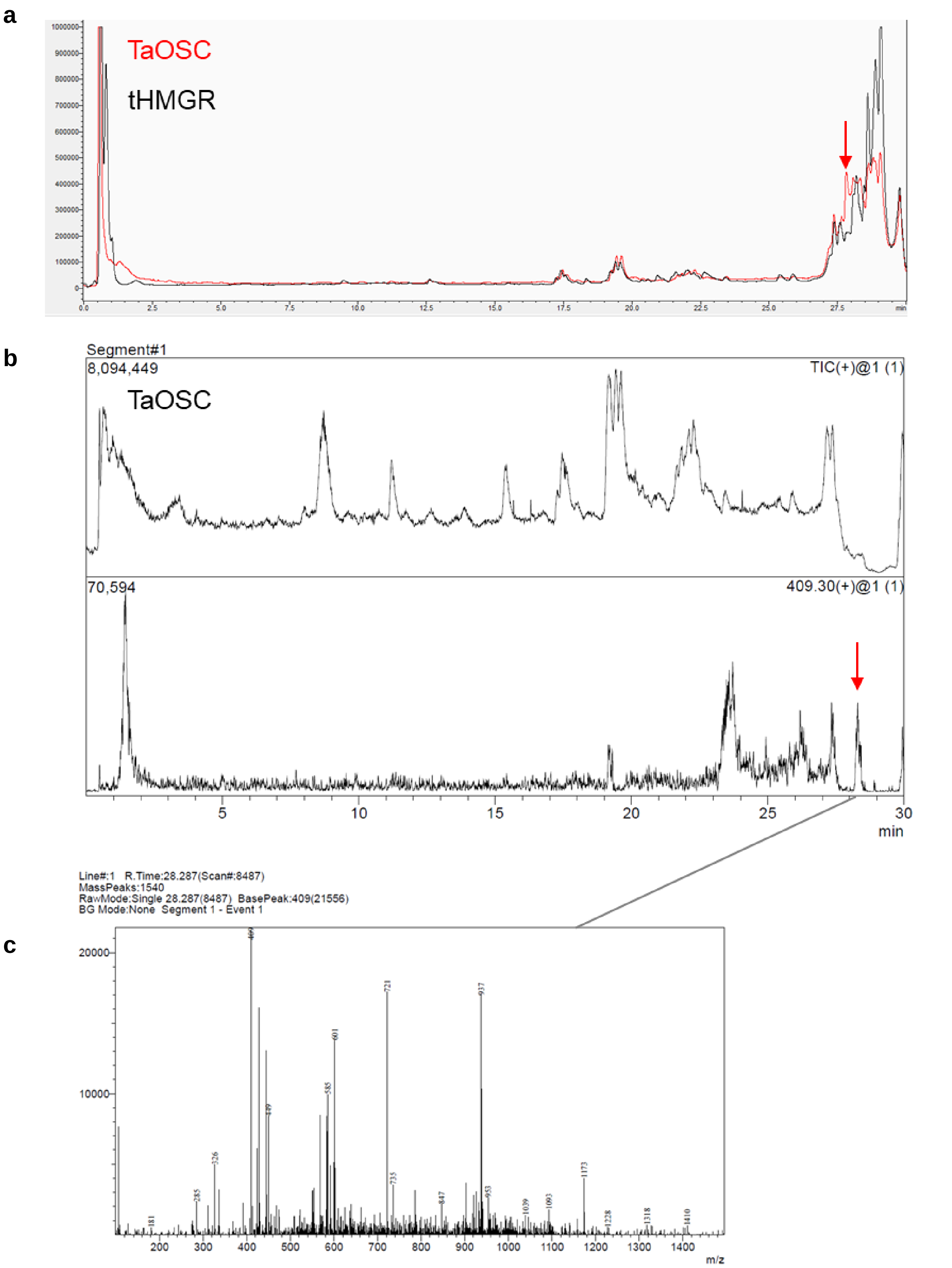


**Supplementary Fig. 6. LC-MS detection of isoarborinol molecular ion**. **a**, CAD chromatogram. **b**, MS chromatogram- TIC and EIC of 409.3. **c**, Mass spectra at peak retention time.

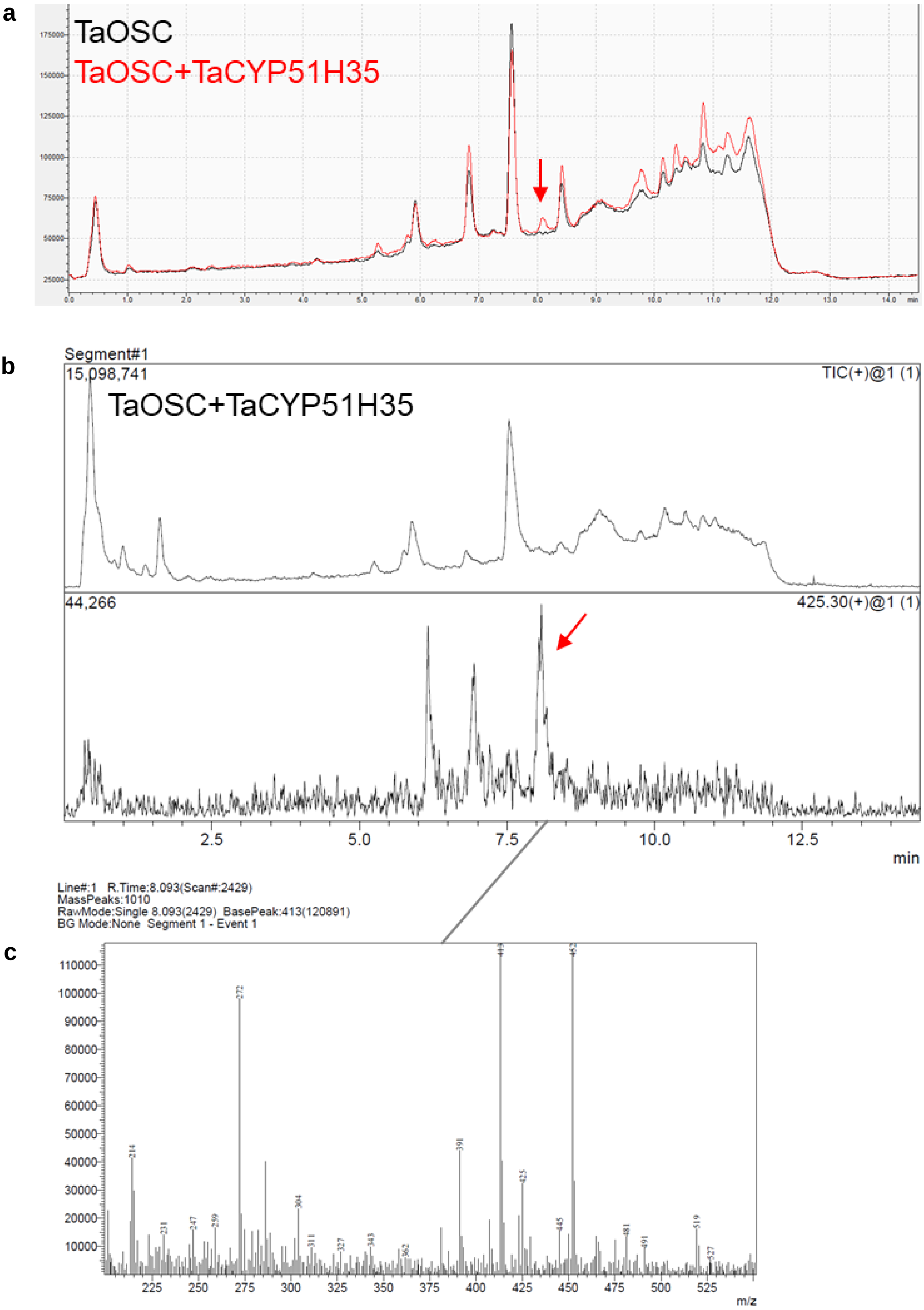


**Supplementary Fig. 7. LC-MS detection of 19-hydroxy-isoarborinol molecular ion. a**, CAD chromatogram. **b**, MS chromatogram- TIC and EIC of 441.3. **c**, Mass spectra at peak retention time.

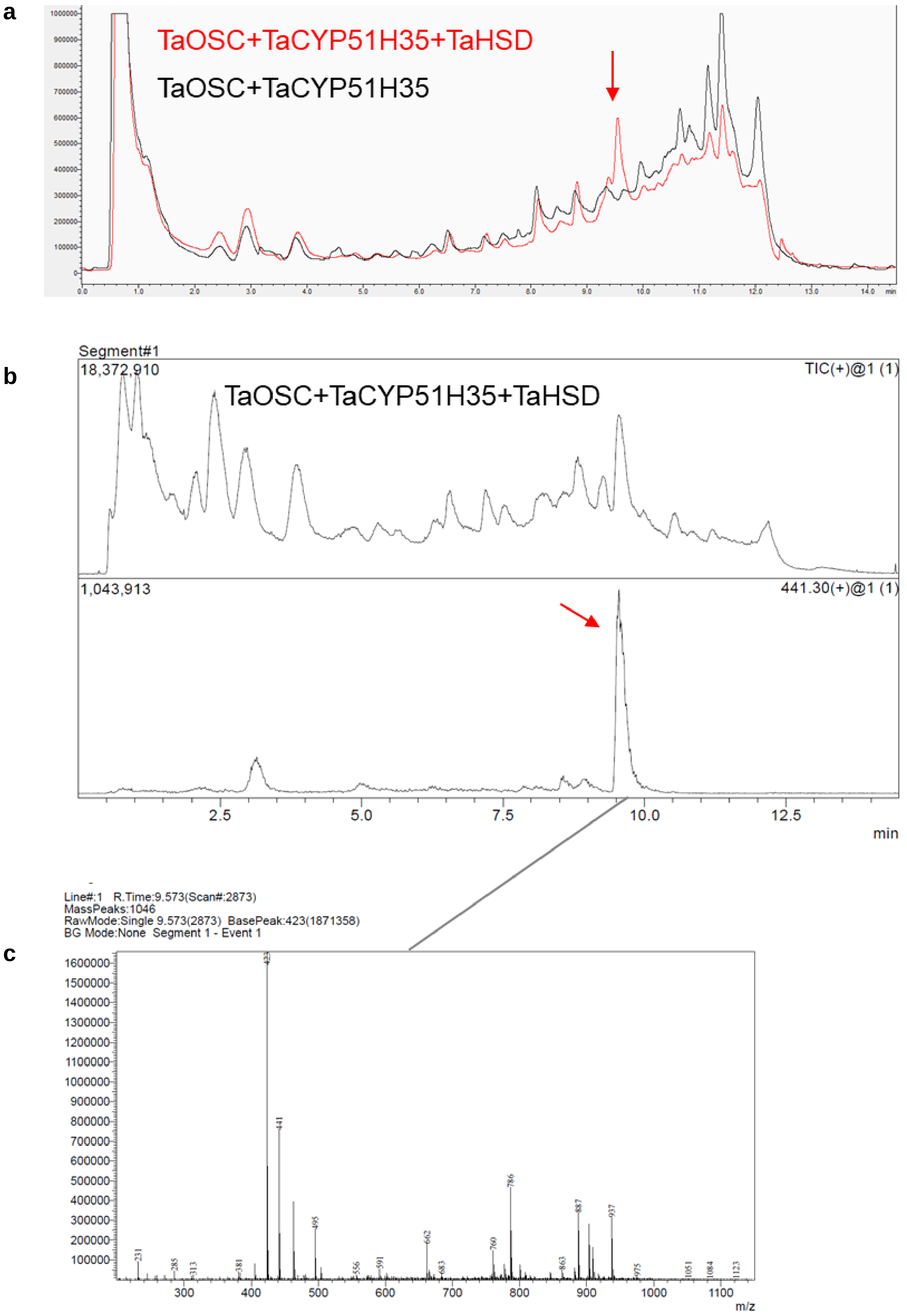


**Supplementary Fig. 8. LC-MS detection of 19-hydroxy-isoarborinone molecular ion**. **a**, CAD chromatogram. **b**, MS chromatogram- TIC and EIC of 441.3. **c**, mass spectra at peak retention time.

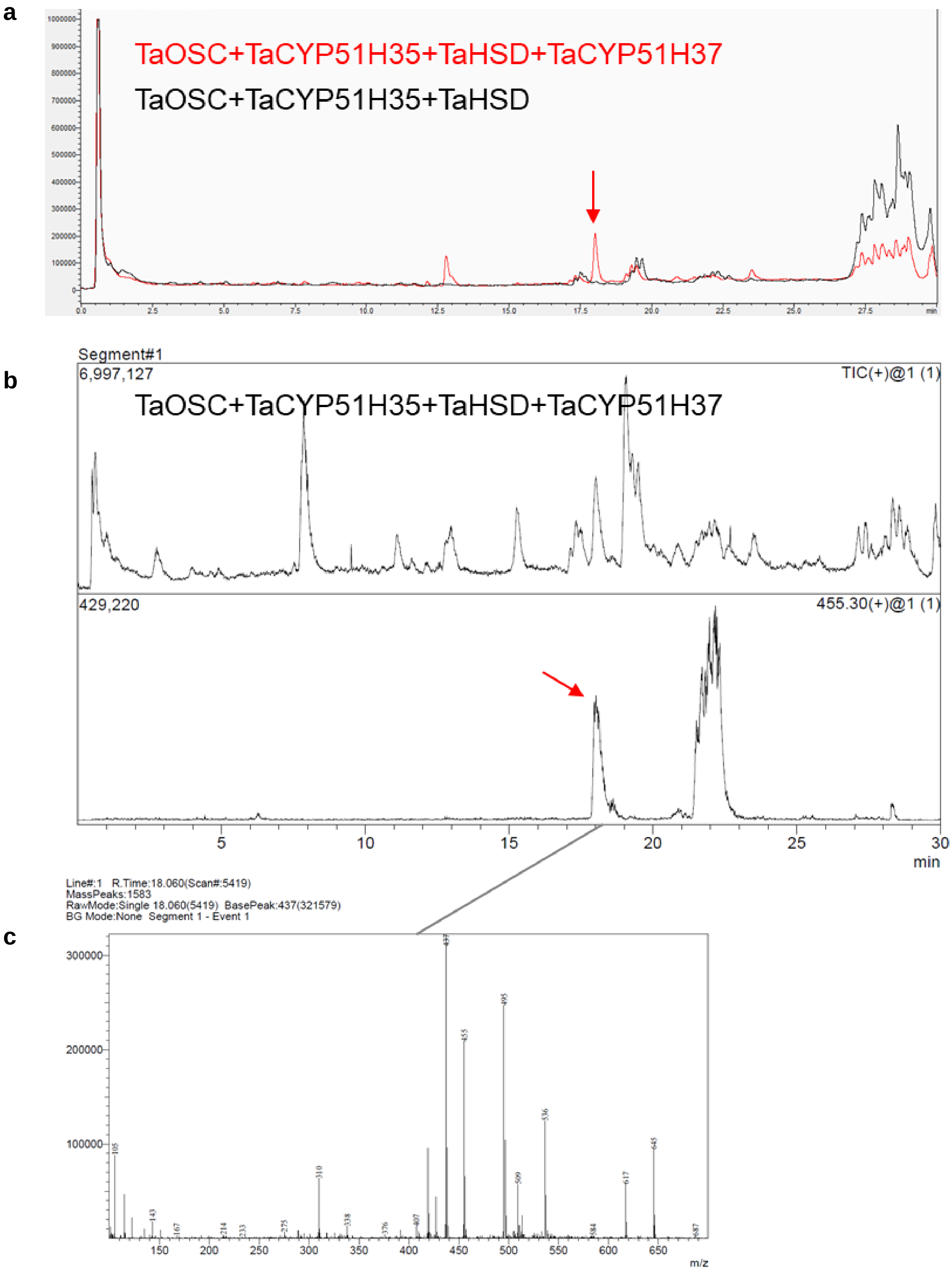


**Supplementary Fig. 9. LC-MS detection of ellarinacin molecular ion**. **a**, CAD chromatogram. **b**, MS chromatogram- TIC and EIC of 455.3. **c**, Mass spectra at peak retention time.

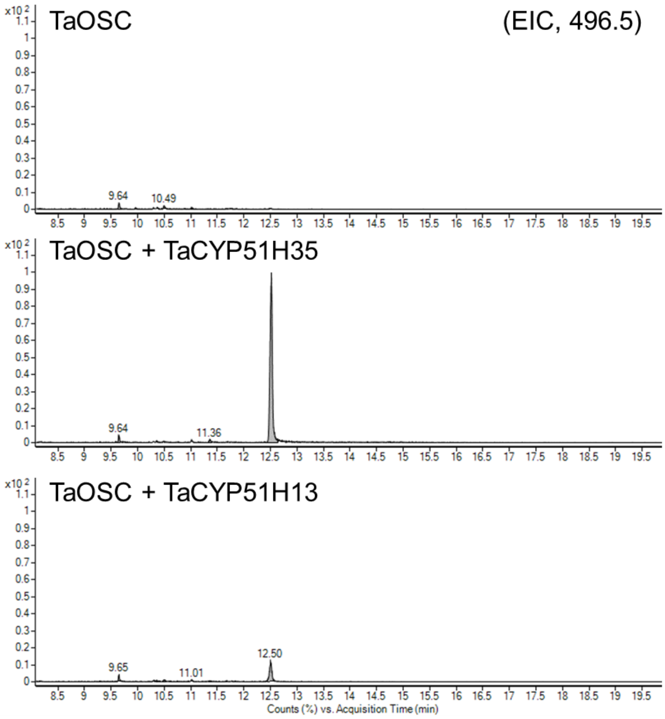


**Supplementary Fig. 10. GC-MS analysis of wheat TaCYP51H35 and TaCYP51H13 expression in *N. benthamiana.*** EIC, extracted ion chromatogram of fragment ion representing 19-hydroxy-isoarborinol (496.5). Y-axes are linked. Oat tHMGR was included in all combinations of genes.

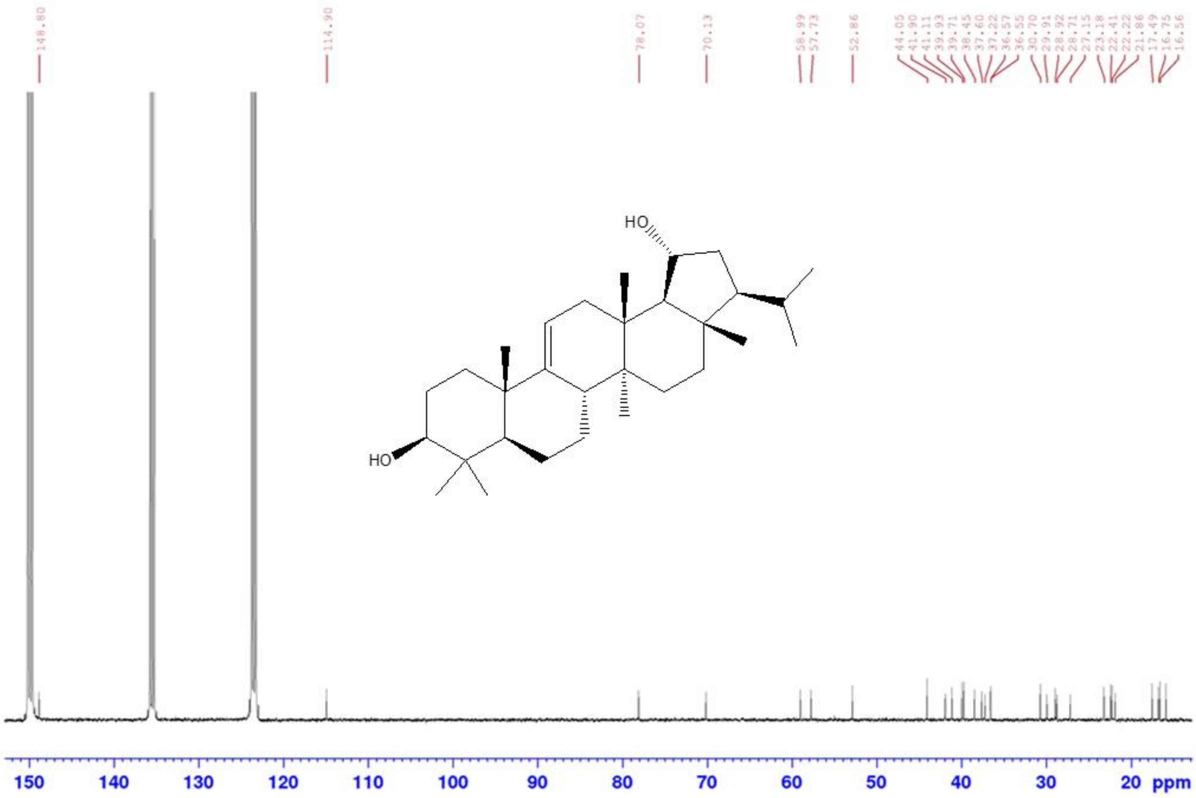


**Supplementary Fig. 11.** $^{13}$C **19-hydroxy-isoarborinol** (**Pyridine-d5)**. Referenced to the most downfield peak reported in the literature[6]. 400 MHz instrument.

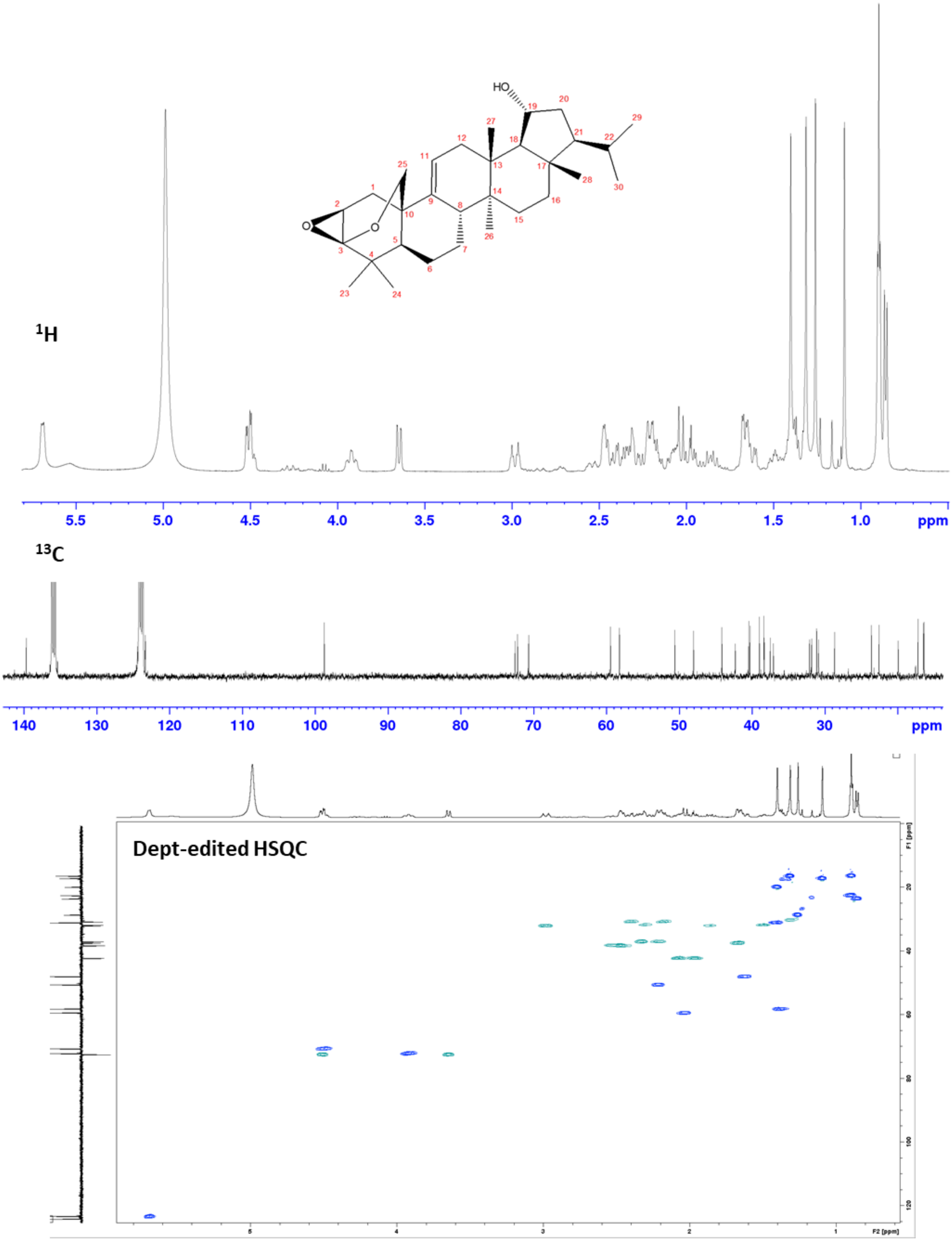


**Supplementary Fig. 12. ¹H, ¹³C, and dept-edited HSQC spectra for ellarinacin, (pyridine-d5).** Referenced to residual solvent peak (¹H δ: 8.74) (¹³C δ: 150.3)]. 400 MHz instrument.

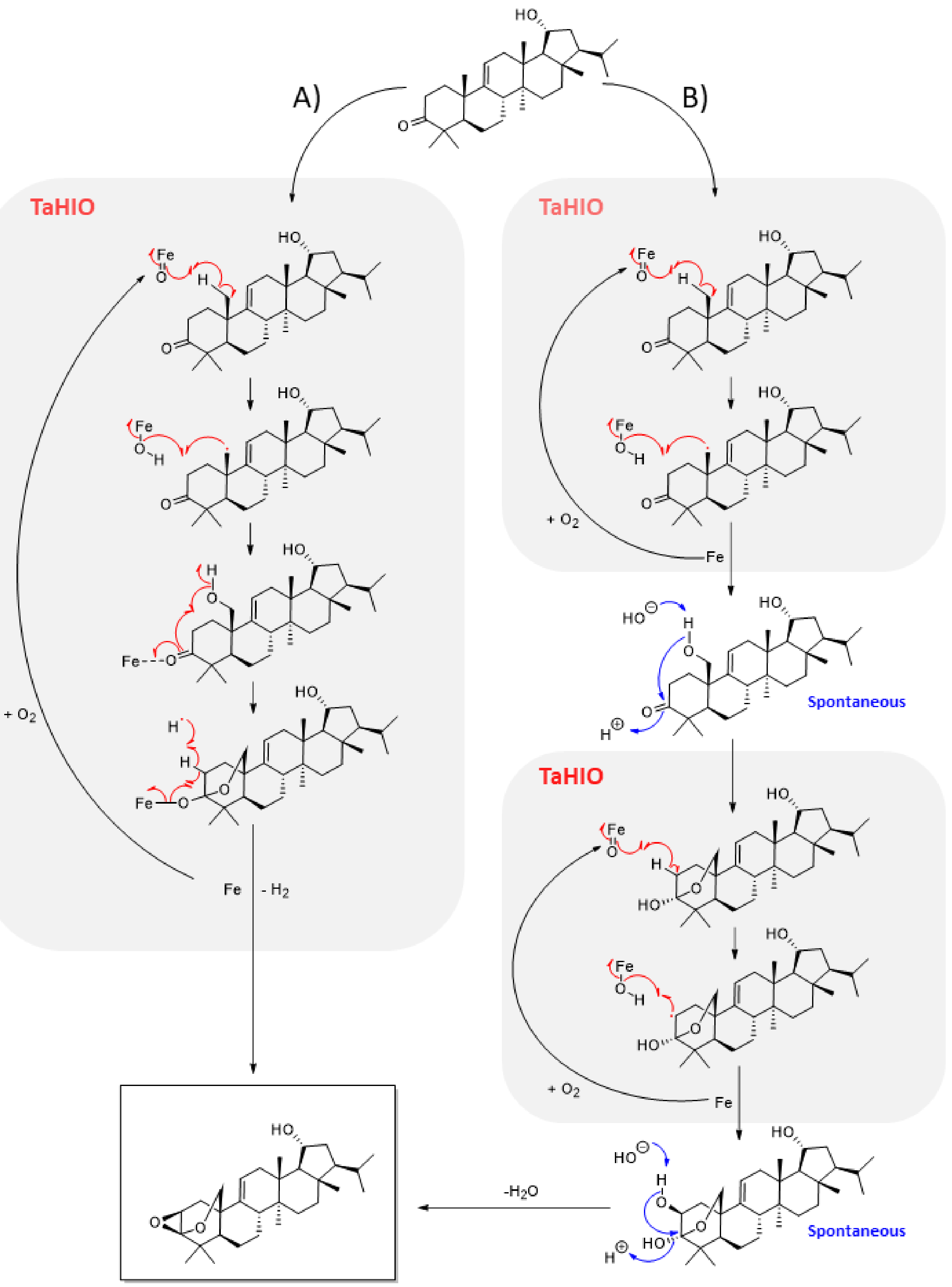


**Supplementary Fig. 13. Possible reaction mechanisms of TaHIO (TaCYP51H37).** Reaction mechanism may involve one catalytic cycle (A) or alternatively involve two independent catalytic cycles (B).

| | | 1. | 2. | 3. | 4. | 5. | 6. | 7. | 8. | 9. | 10. | 11. | 12. |
|---|---|---|---|---|---|---|---|---|---|---|---|---|---|
| 1. | AK067451\|O.sativa\|IAS | 100 | 62.3 | 63.8 | 49.8 | 56.4 | 56.4 | 56.2 | 54.7 | 54.8 | 56.4 | 56.2 | 55.6 |
| 2. | AJ311789\|A.strigosa\|bAS | 62.3 | 100 | 68 | 48.3 | 53.8 | 53.7 | 53.6 | 51.9 | 52.1 | 55.4 | 54.4 | 53.3 |
| 3. | AK070534\|O.sativa\|ABS | 63.8 | 68 | 100 | 48.1 | 55.6 | 57.1 | 57.1 | 53.3 | 53.4 | 57.2 | 56.9 | 56.6 |
| 4. | LOC4344966\|O.sativa\|PTS | 49.8 | 48.3 | 48.1 | 100 | 56.1 | 55.3 | 55.5 | 53.9 | 53.2 | 56.1 | 54.9 | 54.4 |
| 5. | Bradi3g22802\|B.distachyon\|IAS | 56.4 | 53.8 | 55.6 | 56.1 | 100 | 90.4 | 88.3 | 63 | 62.6 | 71.5 | 69.8 | 69.6 |
| 6. | AS01G014480\|A.strigosa\|IAS | 56.4 | 53.7 | 57.1 | 55.3 | 90.4 | 100 | 91.3 | 63.5 | 63 | 73.5 | 71.2 | 71.5 |
| 7. | TraesCS5D02G011800\|T.aestivum\|IAS | 56.2 | 53.6 | 57.1 | 55.5 | 88.3 | 91.3 | 100 | 62.5 | 62.1 | 71.6 | 69.5 | 69.8 |
| 8. | AK066327\|O.sativa\|PS | 54.7 | 51.9 | 53.3 | 53.9 | 63 | 63.5 | 62.5 | 100 | 95.7 | 71.5 | 69 | 67.6 |
| 9. | AYV65354\|O.sativa\|ORS | 54.8 | 52.1 | 53.4 | 53.2 | 62.6 | 63 | 62.1 | 95.7 | 100 | 70.3 | 68.1 | 67.1 |
| 10. | AK121211\|O.sativa\|CAS | 56.4 | 55.4 | 57.2 | 56.1 | 71.5 | 73.5 | 71.6 | 71.5 | 70.3 | 100 | 87.2 | 86.4 |
| 11. | AJ311790\|A.strigosa\|CAS | 56.2 | 54.4 | 56.9 | 54.9 | 69.8 | 71.2 | 69.5 | 69 | 68.1 | 87.2 | 100 | 92.2 |
| 12. | TraesCS6D02G099400\|T.aestivum\|CAS* | 55.6 | 53.3 | 56.6 | 54.4 | 69.6 | 71.5 | 69.8 | 67.6 | 67.1 | 86.4 | 92.2 | 100 |

**Supplementary Fig. 14. Percent identity matrix of amino acid sequences of characterised Poaceae cycloartenol- and triterpene- synthases**. Sequence alignment and percent identity matrix was generated with Clustal Omega v2.1. IAS, isoarborinol synthase; bAS, beta-amyrin synthase; ABS, achilleol b synthase; PTS, poaceaetapetol synthase; PS, parkeol synthase; ORS, orysatinol synthase; CAS, cycloartenol synthase. Asterisk denotes a predicted *T. aestivum* CAS annotation, based on phylogeny and a constitutive expression pattern in wheat RNA-seq data.

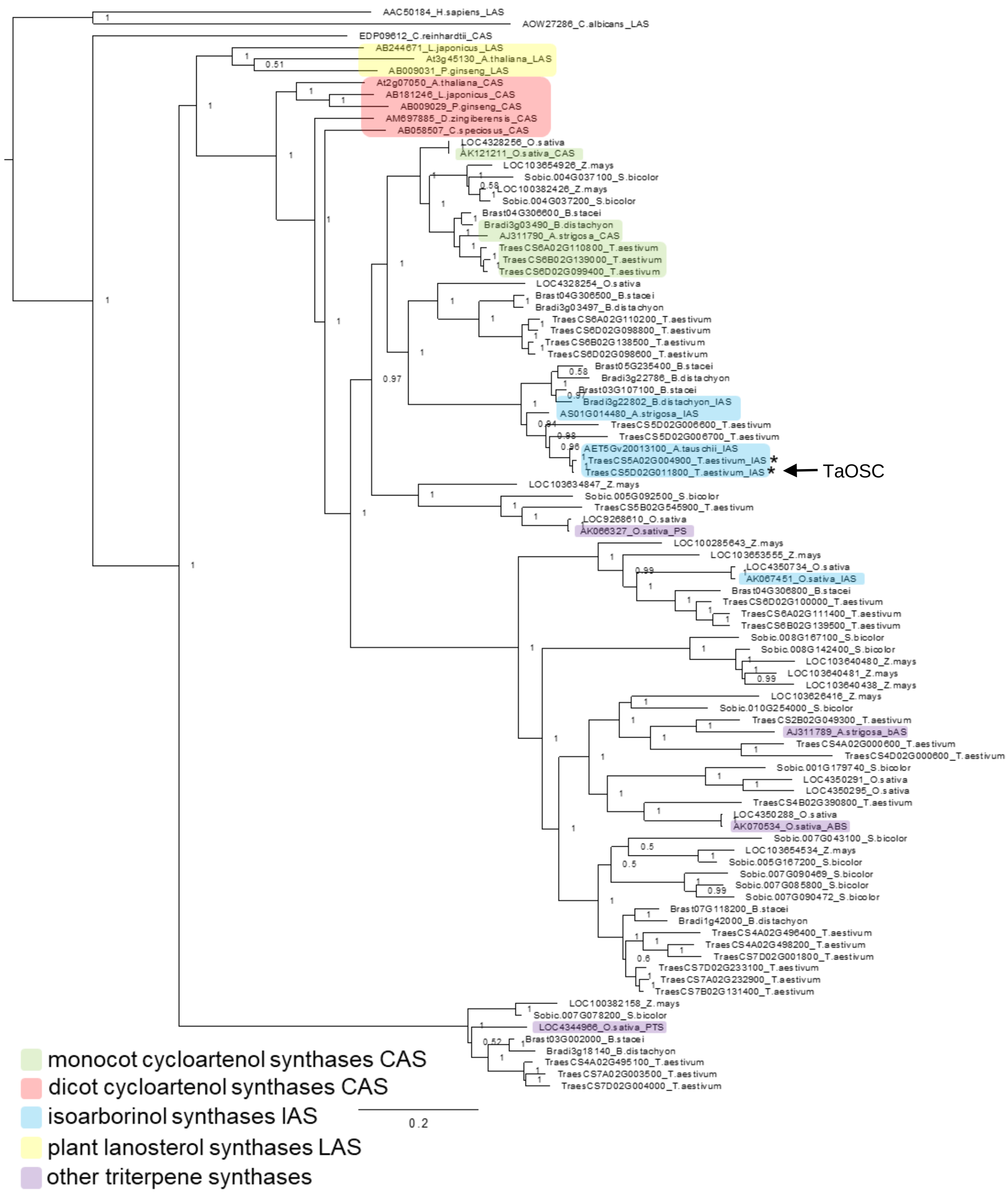


**Supplementary Fig. 15. phylogeny of predicted oxidosqualene cyclases from wheat, together with selected monocot and dicot OSCs**. Bayesian tree comprised of protein sequences of characterized and non-characterized monocot OSCs and characterized dicot cycloartenol and lanosterol synthases. Cycloartenol synthase from *Chlamydomonas reinhardtii* and lanosterol synthases from *Homo sapiens* and *Candida albicans* are included as outgroups. Sequences were aligned with MUSCLE (with a maximum of 100 iterations) and a phylogenetic tree was generated using MrBayes[7], with a mixed amino acid probability model and default MCMC parameters except 0.7 temperature. TaOSC sequences from 3(5D) and 3(5A) clusters are asterisked. *T. aestivum* and *B. distachyon* CAS annotations are predicted, based on phylogeny and constitutive expression patterns in RNA-seq data. CAS, cycloartenol synthase; IAS, isoarborinol synthase; LAS, lanosterol synthase; bAS, beta-amyrin synthase; PS, parkeol synthase; ABS, achilleol B synthase; PTS, poaceatapetol synthase.

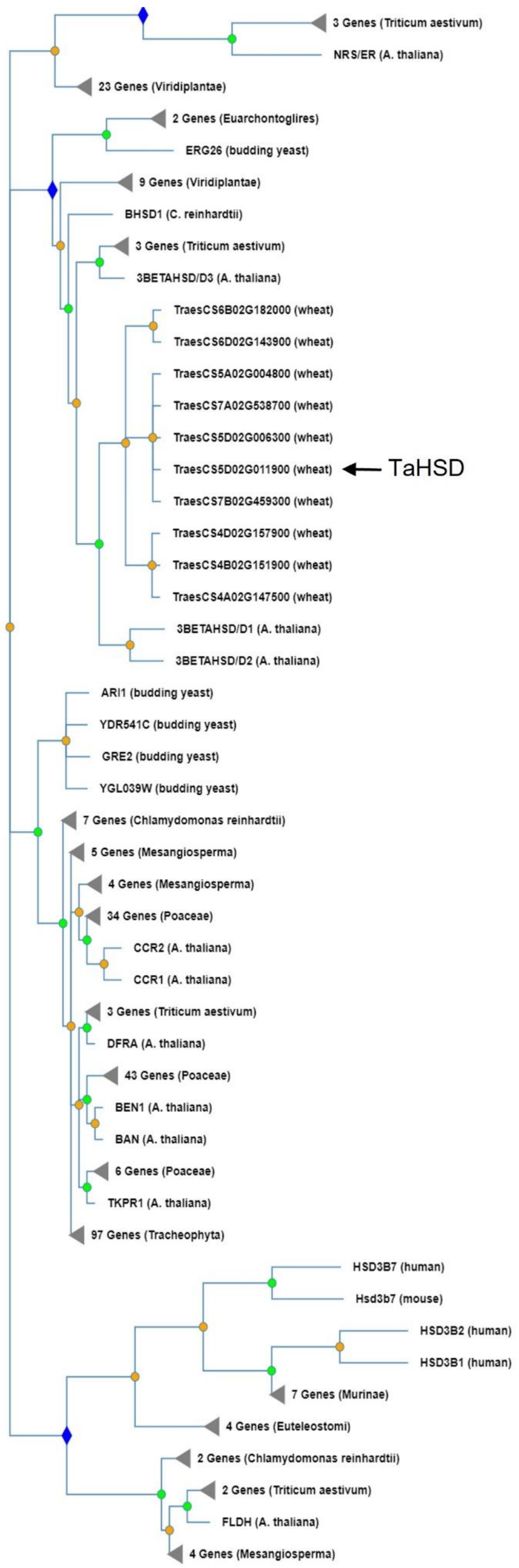


**Supplementary Fig. 16. Phylogenetic tree of selected members of the NAD dependent epimerase/dehydratase family (Panther family PTHR10366).** The tree was constructed in PhyloGenes v2.0 (http://www.phylogenes.org/tree/PTHR10366). TaHSD is clustered with 3βHSD/D1 and 3βHSD/D2 that are take part in sterol biosynthesis in *Arabidopsis thaliana*[8].

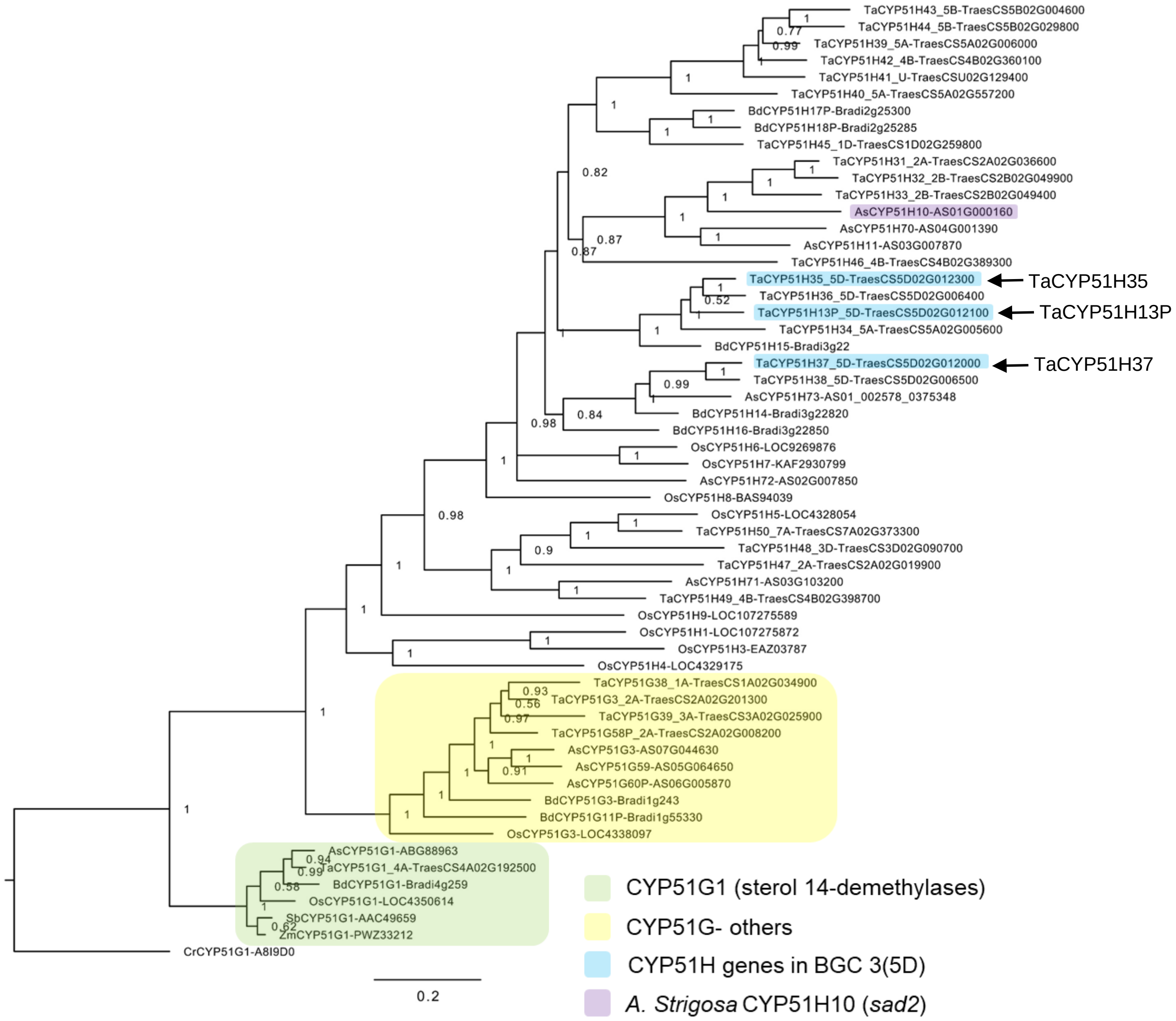


**Supplementary Fig. 17. Phylogeny of CYP51-family cytochrome P450 proteins identified in wheat, brachypodium, oat and rice**. Peptide sequences of CYP51s were aligned with MUSCLE (with a maximum of 100 iterations) and a phylogenetic tree was generated using MrBayes[7], with a mixed amino acid probability model and default MCMC parameters. The tree includes one representative homoeolog of each assigned wheat CYP51. For a tree including all identified wheat homoeologs, see Supplementary Fig. 18. Ta, *Triticum aestivum*; Bd, *Brachypodium distachyon*; Os, *Oryza sativa*; As, *Avena strigosa*; Zm, *Zea mays*; Sb, *Sorghum bicolor*; Cr, *Chlamydomonas reinhardtii*.

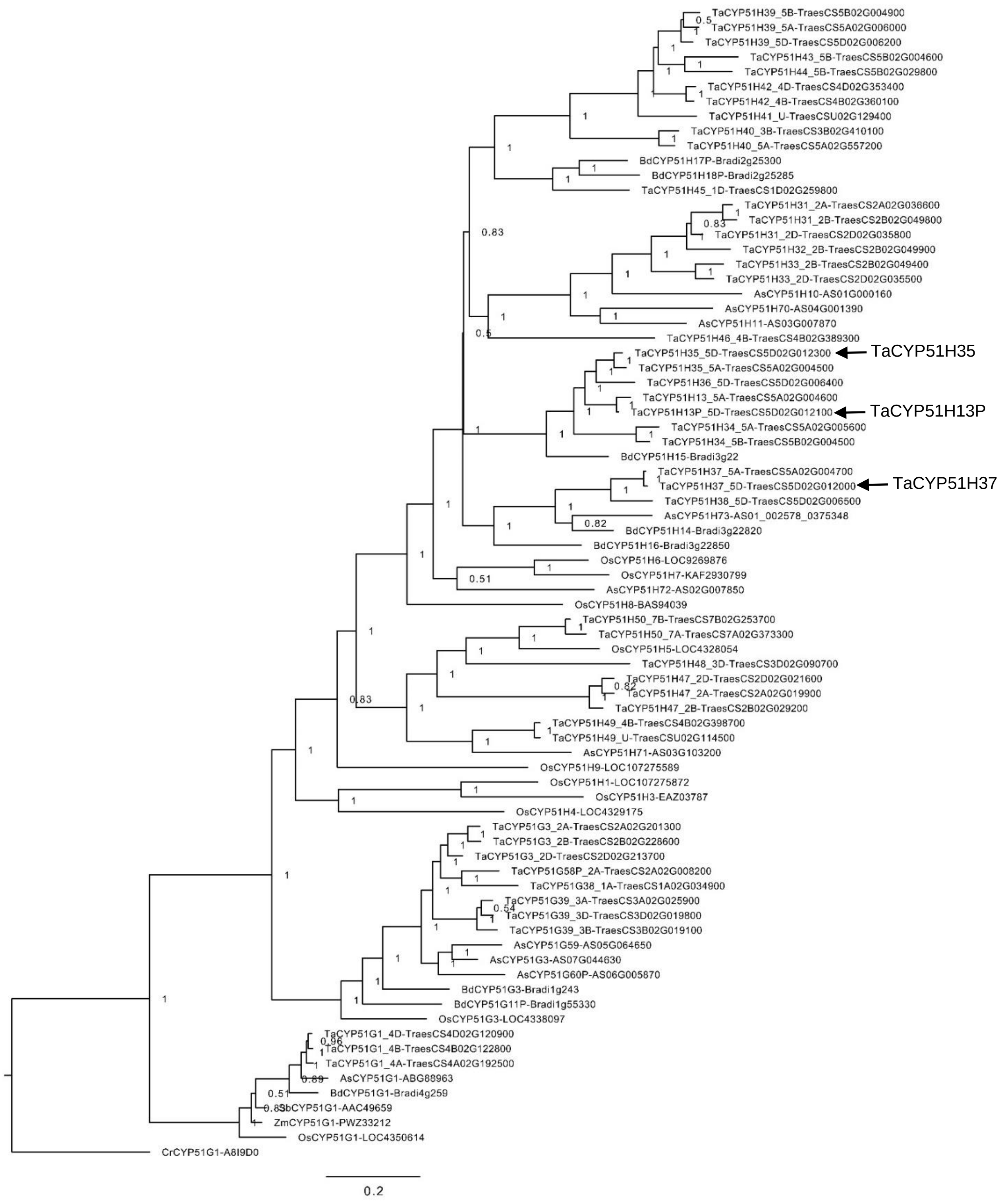


**Supplementary Fig. 18.** Bayesian tree of CYP51 genes from wheat, brachypodium, oat and rice.

**a**

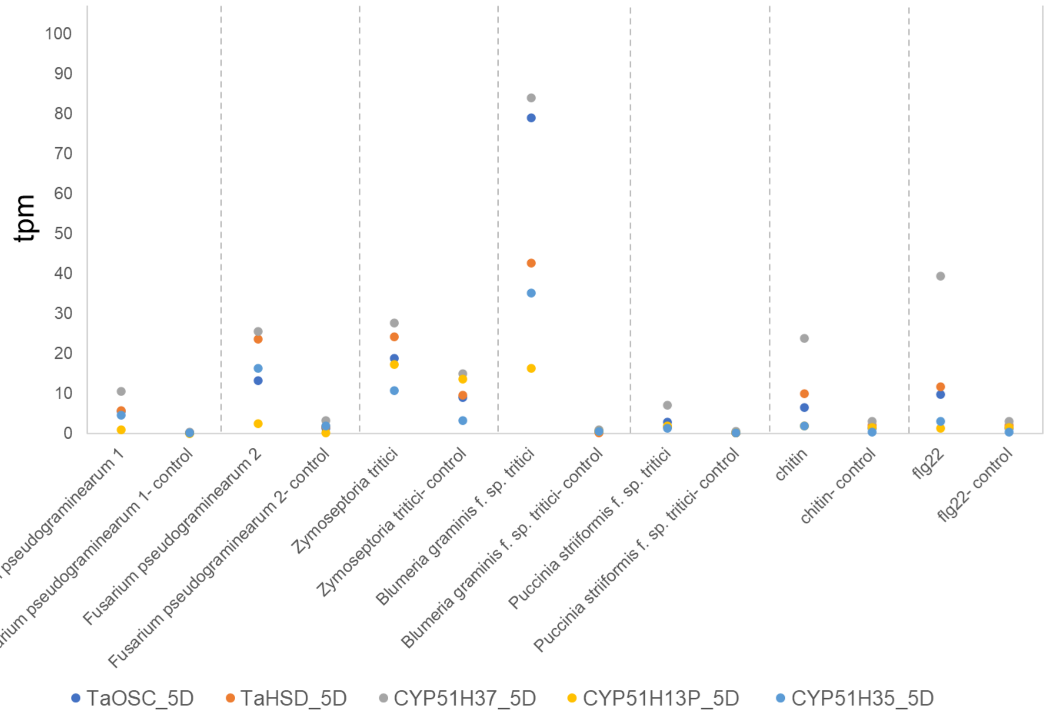


**b**

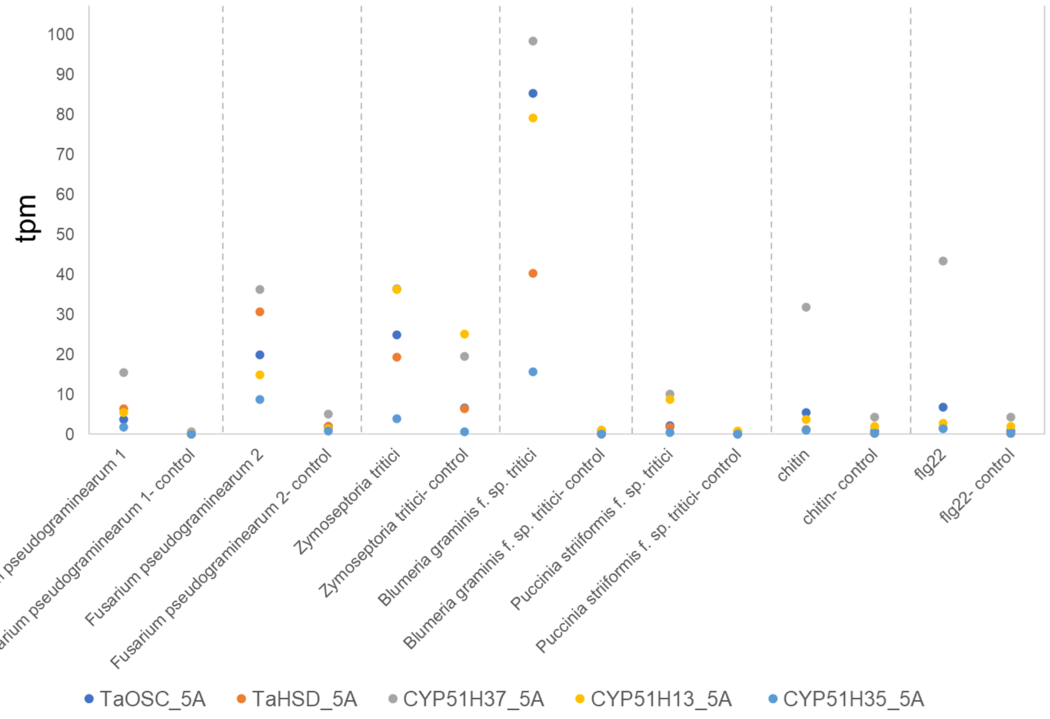


**Supplementary Fig. 19. Expression of wheat clustered genes under biotic stress. a,** genes in Chr.5D cluster. **b**, genes in Chr.5A cluster. Expression data extracted from http://www.wheat-expression.com includes studies applying treatment with *Fusarium pseudograminearum* (study 1:[9], study 2:[10]), *Zymoseptoria tritici*[11], *Blumeria graminis*[12], *Puccinia striiformis*[13], chitin[4], or flg22[4]. TaCYP51H13P_5D values are average of TraesCS5D01G012100 and TraesCS5D01G012200. tpm, transcripts per million.

**a**

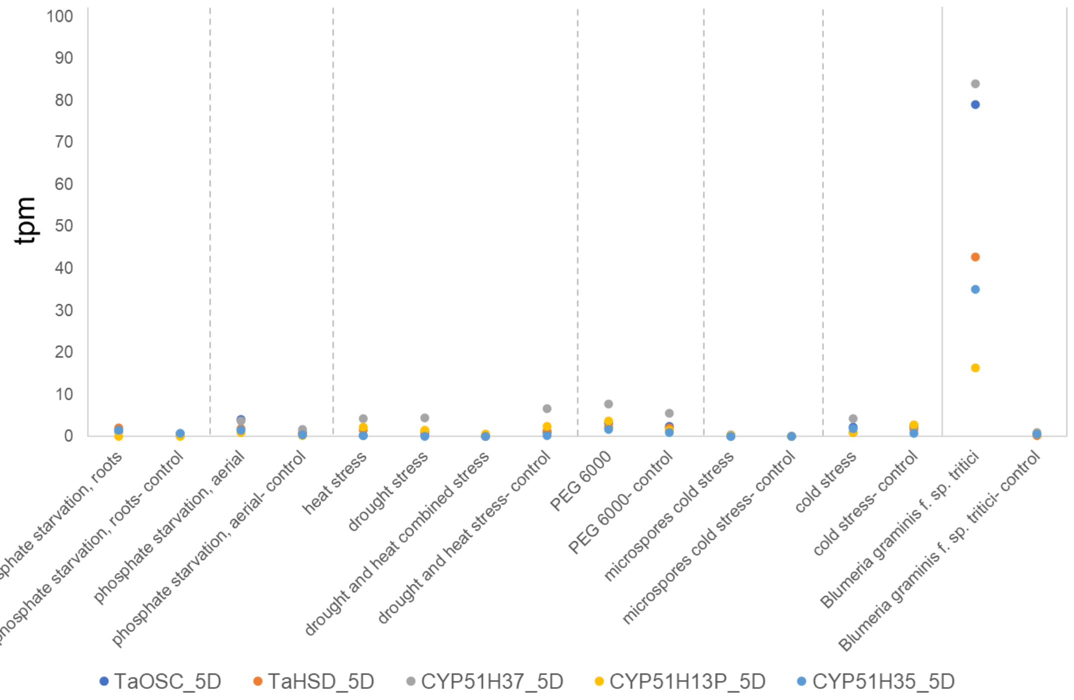


**b**

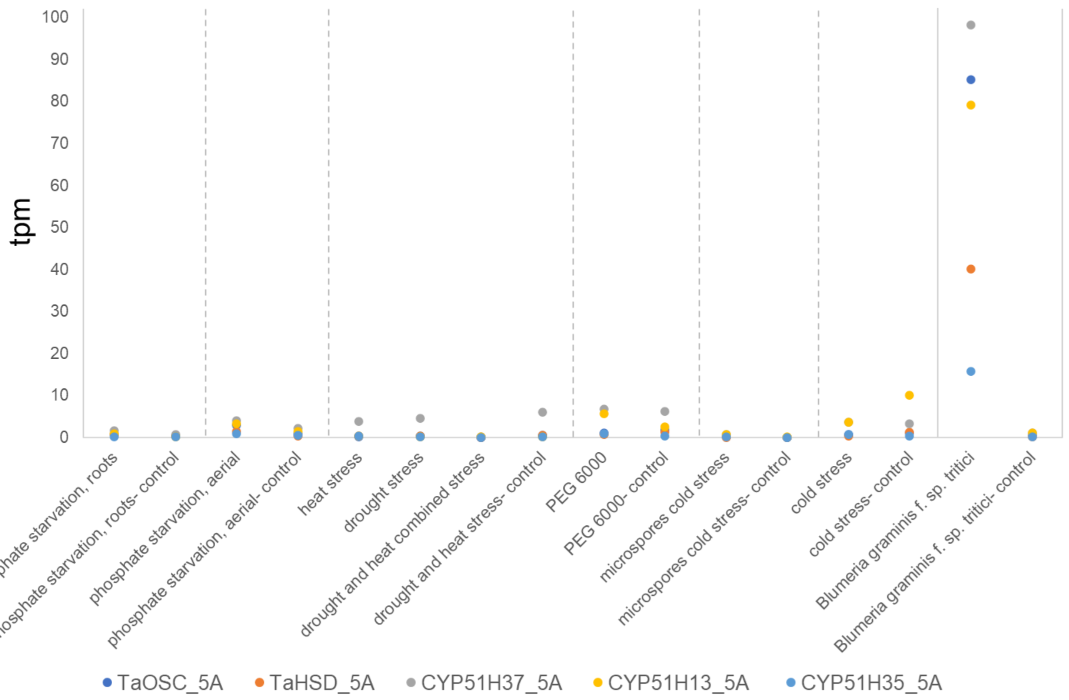


**Supplementary Fig. 20. Expression of wheat clustered genes under abiotic stress**. **a**, genes in Chr.5D cluster. **b**, genes in Chr.5A cluster. Expression data extracted from http://www.wheat-expression.com includes studies applying phosphate starvation[14], heat and drought stress[15], PEG 6000, microspores cold stress[16] and cold stress[17]. TaCYP51H13P_5D values are average of TraesCS5D01G012100 and TraesCS5D01G012200. Data of powdery mildew infection experiment (*Blumeria graminis f. sp. tritici*)[12] is shown for comparison. tpm, transcripts per million.

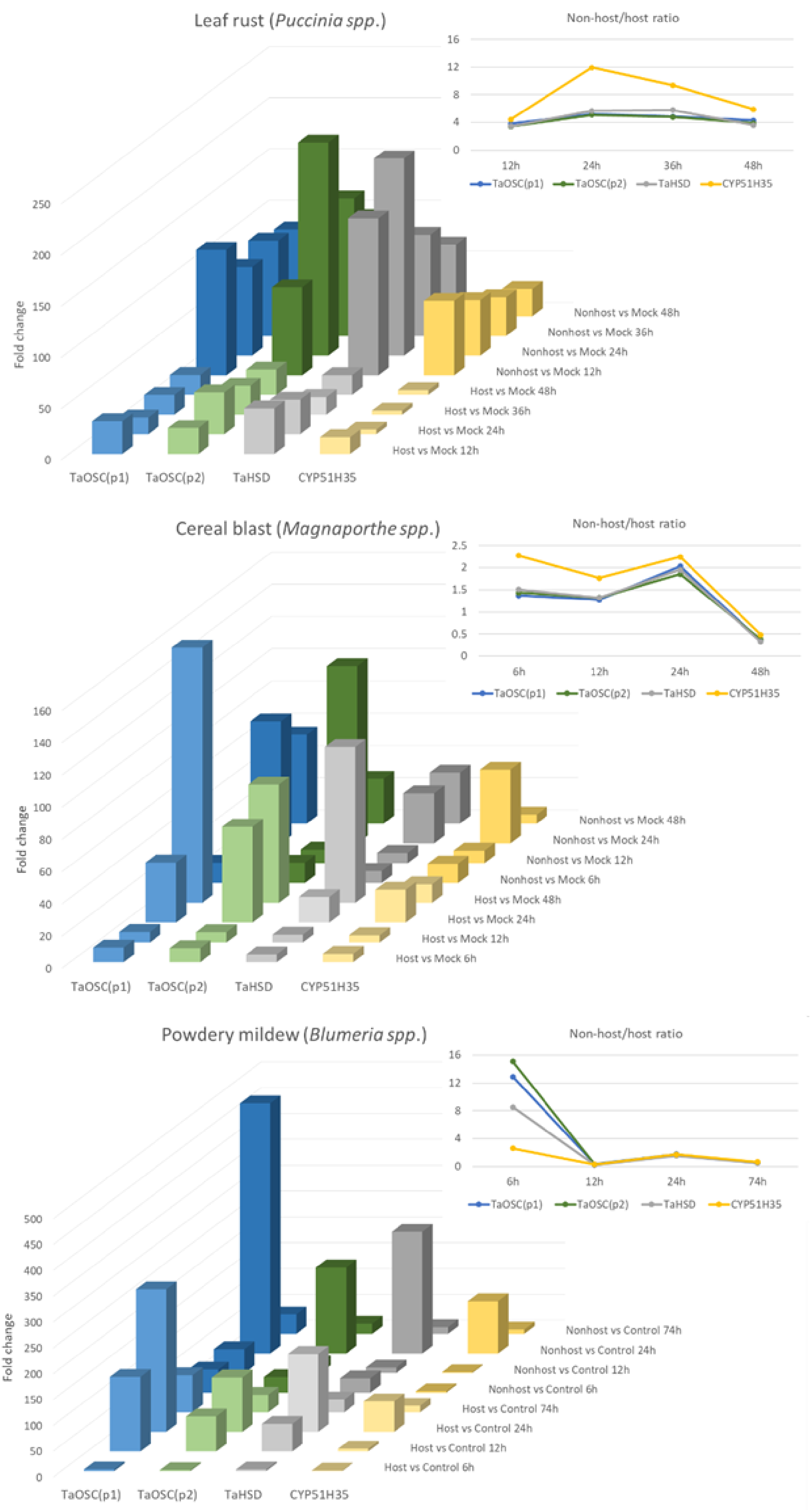


**Supplementary Fig. 21. Wheat clustered genes TaOSC, TaHSD and TaCYP51H35 are induced by infection of wheat plants with adapted and non-adapted isolates of leaf rust, cereal blast, or powdery mildew.** Microarray probe signal intensities are plotted in 3D-columns showing fold change expression in adapted and non-adapted vs. control experiments. 2D plots show fold change ratio of non-adapted/adapted vs. control, per each time point. TaOSC is represented by two probes.

**a**

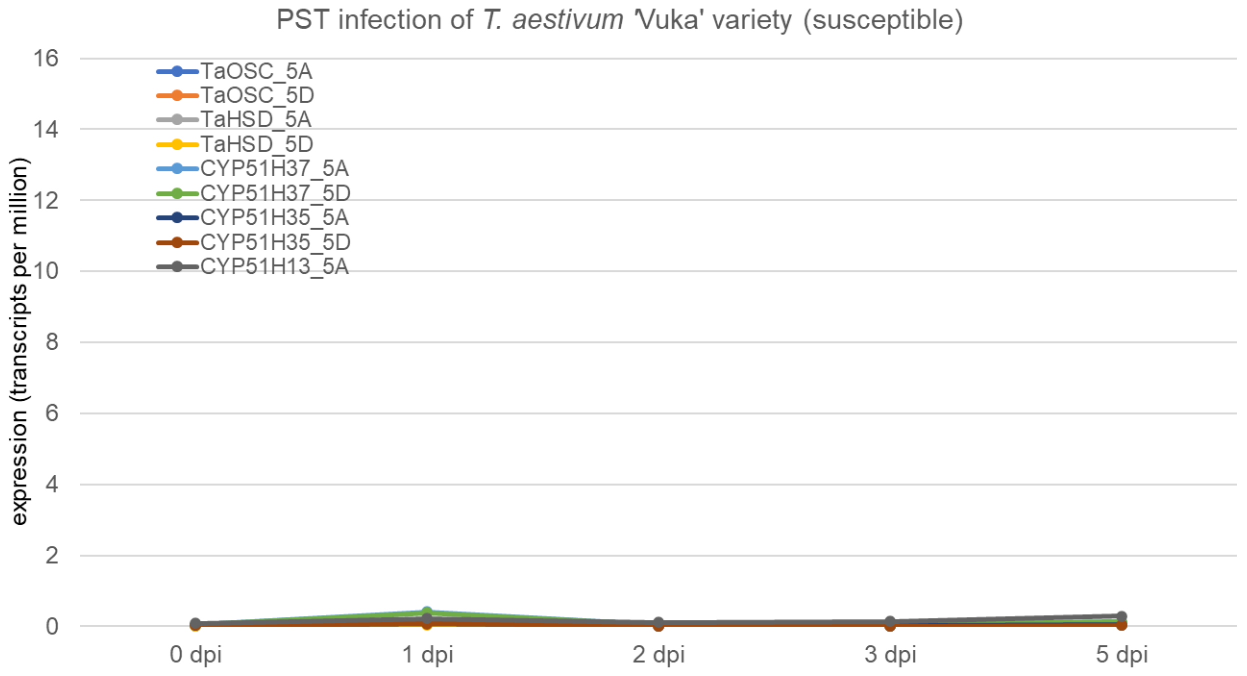


**b**

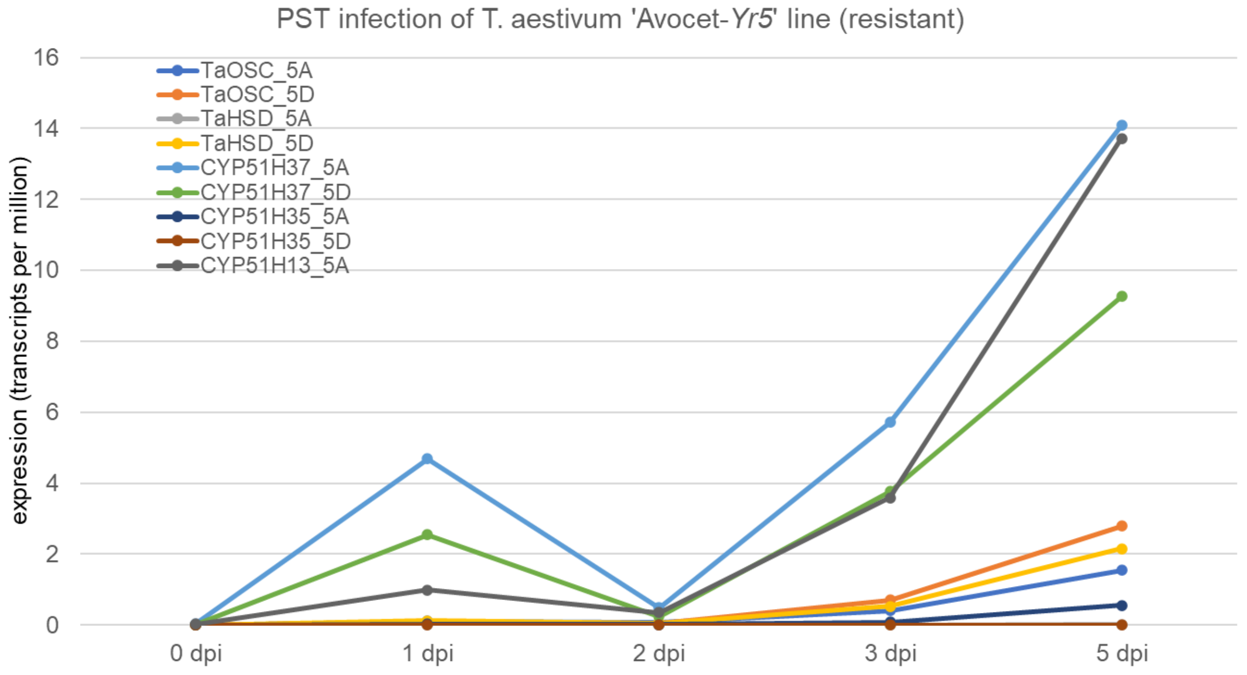


**Supplementary Fig. 22. Induction of wheat cluster genes following infection with yellow rust (*Puccinia striiformis f. sp. tritici*) is suppressed in a susceptible variety.** Expression values were extracted from RNA-seq data of a susceptible wheat variety 'Vuka' (**a**), or a resistant 'Avocet' introgression line (**b**), infected with yellow rust, 0, 1, 2, 3, and 5 days post infection.

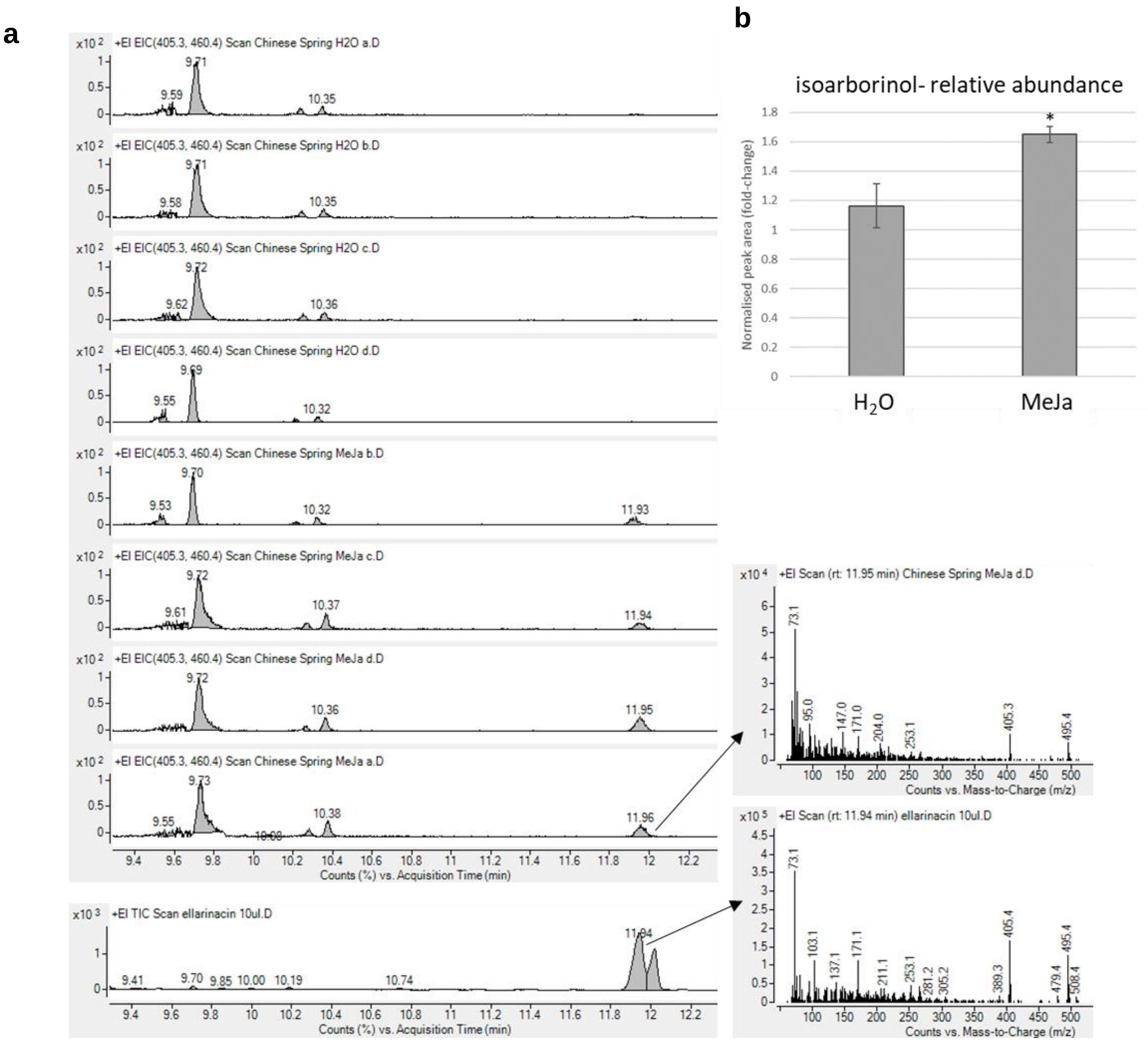


**Supplementary Fig. 23. GC-MS analysis of TMS-derivatized extracts from wheat leaves treated with methyl jasmonate (MeJa), or $H_2O$ (control)**. **a**, extracted ion chromatograms for ions representing ellarinacin (405.3, Rt 11.94) and 5α-cholestan-3β-ol (460.4, Rt 9.70) are shown for extracts from four biological replicates, together with mass spectra of ellarinacin peak at Rt 11.94. Y-axes are linked to peak of internal standard- 5α-cholestan-3β-ol. Wheat leaf extracts were compared to ellarinacin purified from *N. benthamiana* for identification. **b**, relative abundance of isoarborinol in TMS-derivatized extracts of MeJa or $H_2O$-treated wheat leaves, based on GC-MS analysis of four biological replicates.

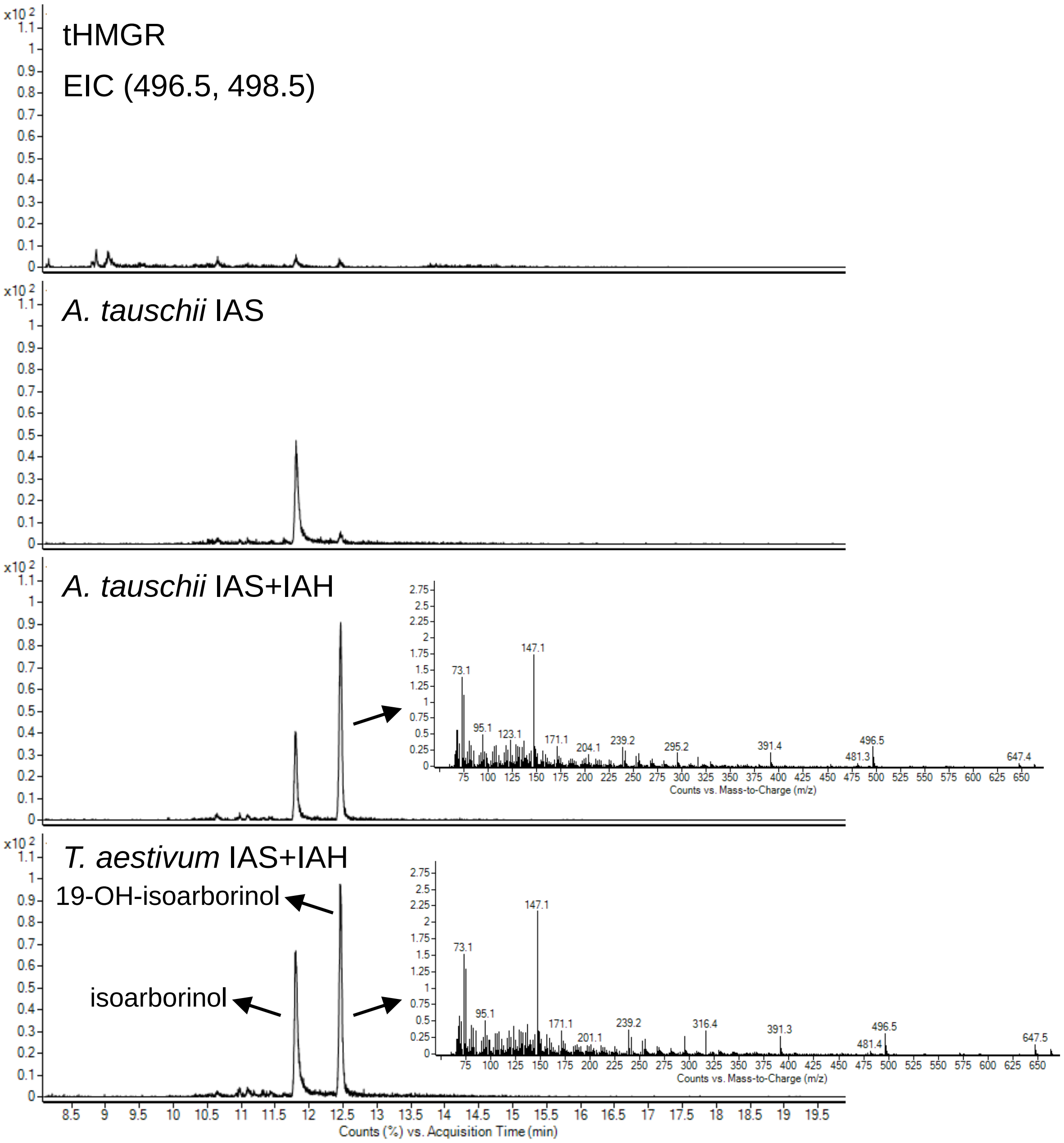


**Supplementary Fig. 24. GC-MS analysis of *Aegilops tauschii* IAS and IAH expression in *N. benthamiana.*** EIC, extracted ion chromatogram of fragment ions representing isoarborinol (498.5) and 19-hydroxy-isoarborinol (496.5). Analysis of wheat IAS and IAH expression is provided for reference.

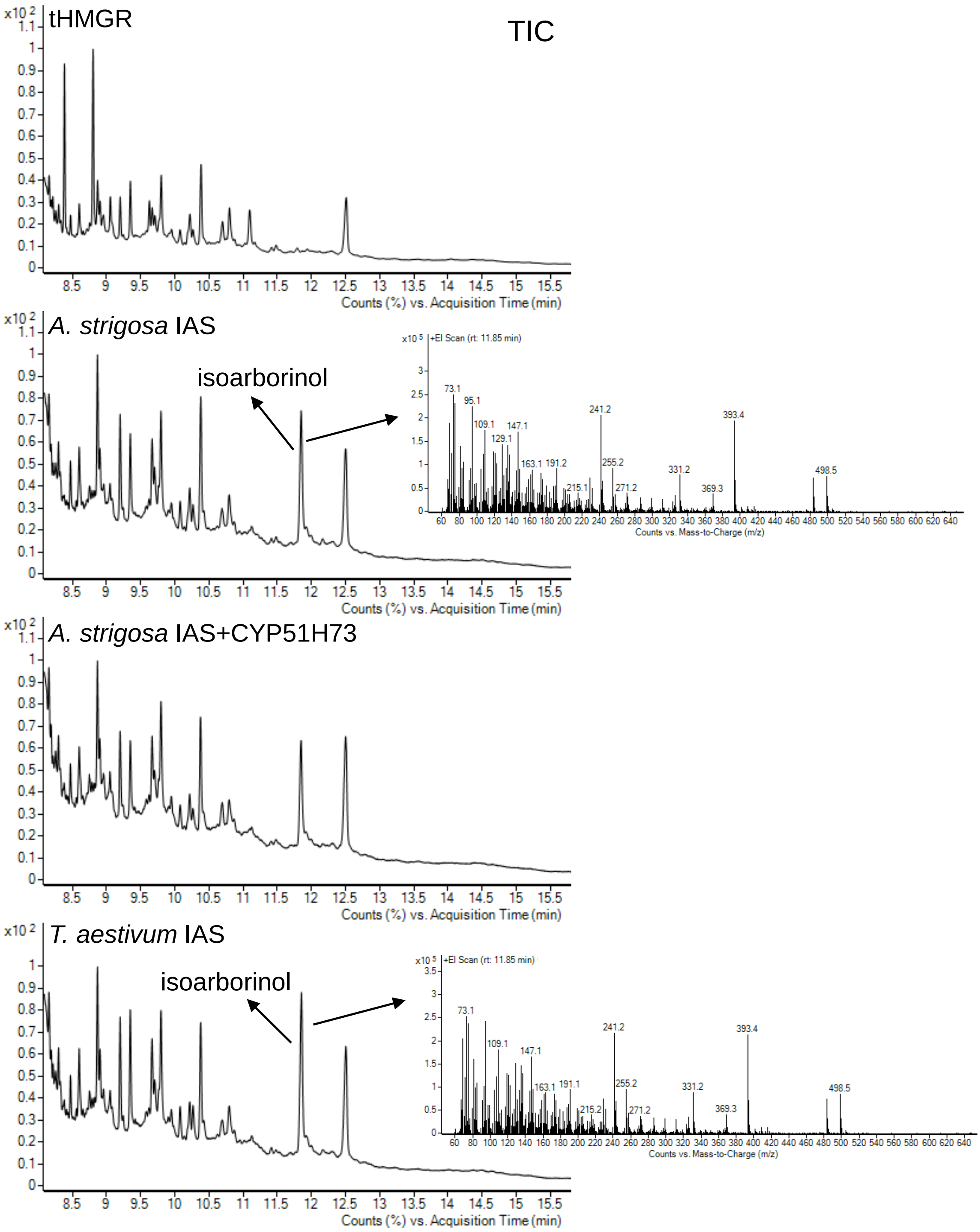


**Supplementary Fig. 25. GC-MS analysis of *Avena strigosa* IAS and CYP51H73 expression in *N. benthamiana*.** TIC, total ion chromatogram. Analysis of wheat IAS expression is provided for reference.

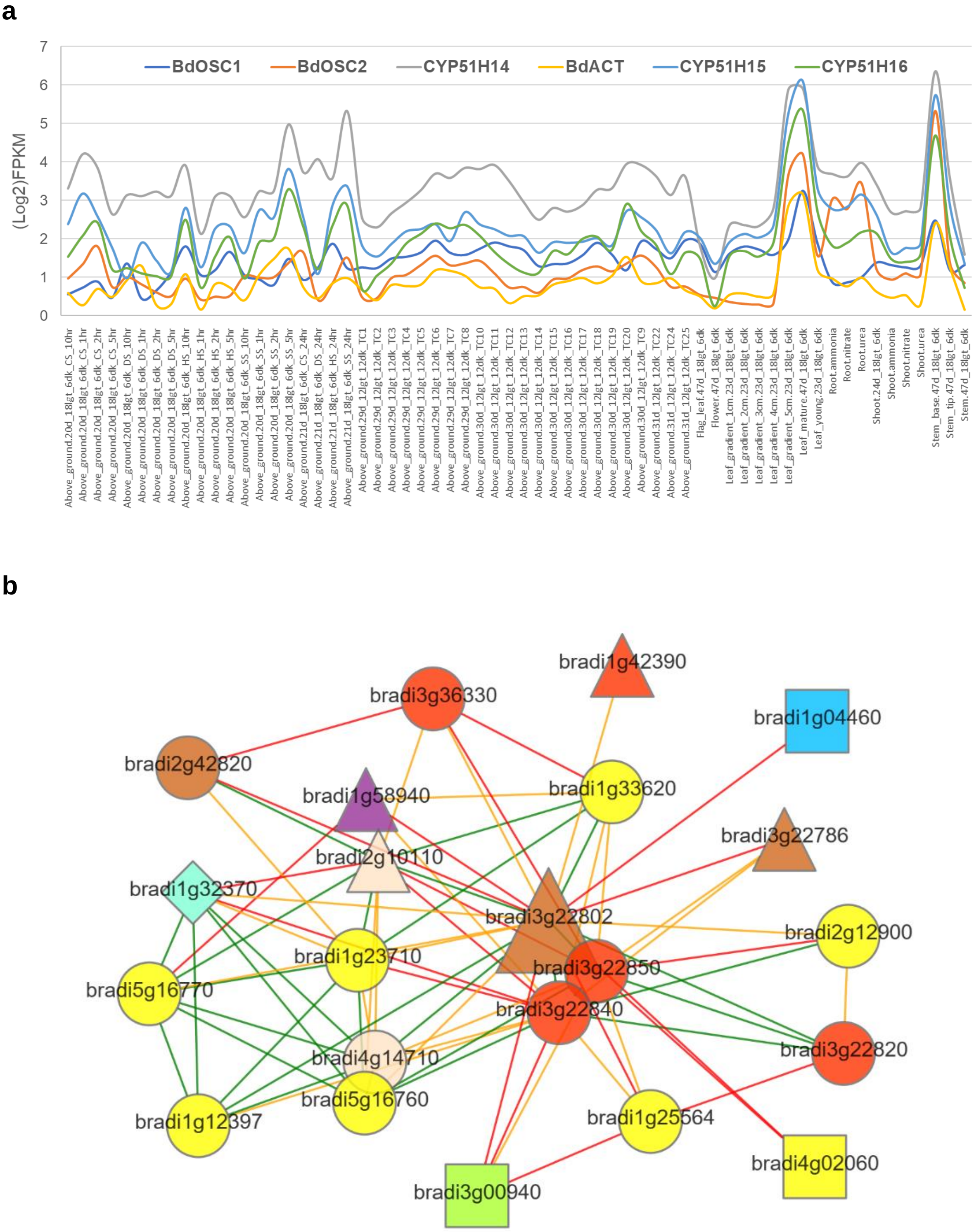


**Supplementary Fig. 26. Triterpene-related clustered genes on *B. distachyon* Chr.03 are co-expressed. a**, gene expression data extracted from JGI Phytozome Gene Atlas (release 1.0), plotted as Log2 of FPKM values. **b**, co-expression network (1st level) neighbourhood of bradi3g22802 (BdOSC2) extracted from PlaNet (http://www.gene2function.de/), shows it is co-expressed with cluster genes: Bradi3g22786, Bradi3g22820, Bradi3g22840 and Bradi3g22850.

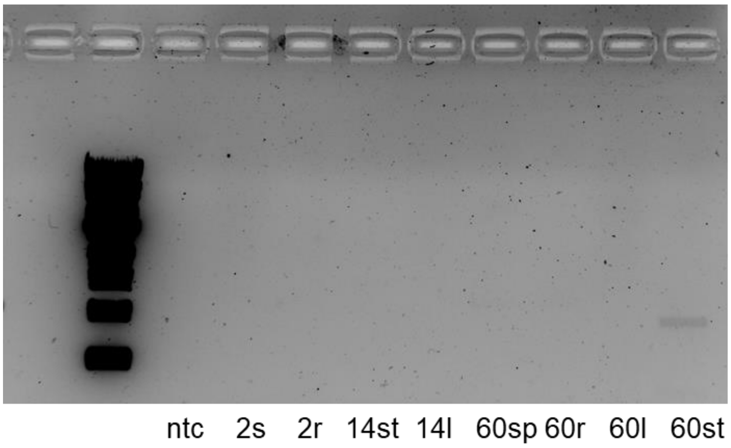


**Supplementary Fig. 27. Semiquantitative RT-PCR of Bradi3g22830.** Amplification of Bradi3g22830 from *B. distachyon* mature stem base cDNA is observed only with prolonged camera exposure (8 sec.). ntc, no template control; 2s, seedling shoot (2 day old); 2r, seedling root; 14st, young plant stem base (14 day old); 14l, young plant leaf; 60sp, mature plant spike (60 day old); 60r, mature plant root; 60l, mature plant leaf; 60st, mature plant stem base.

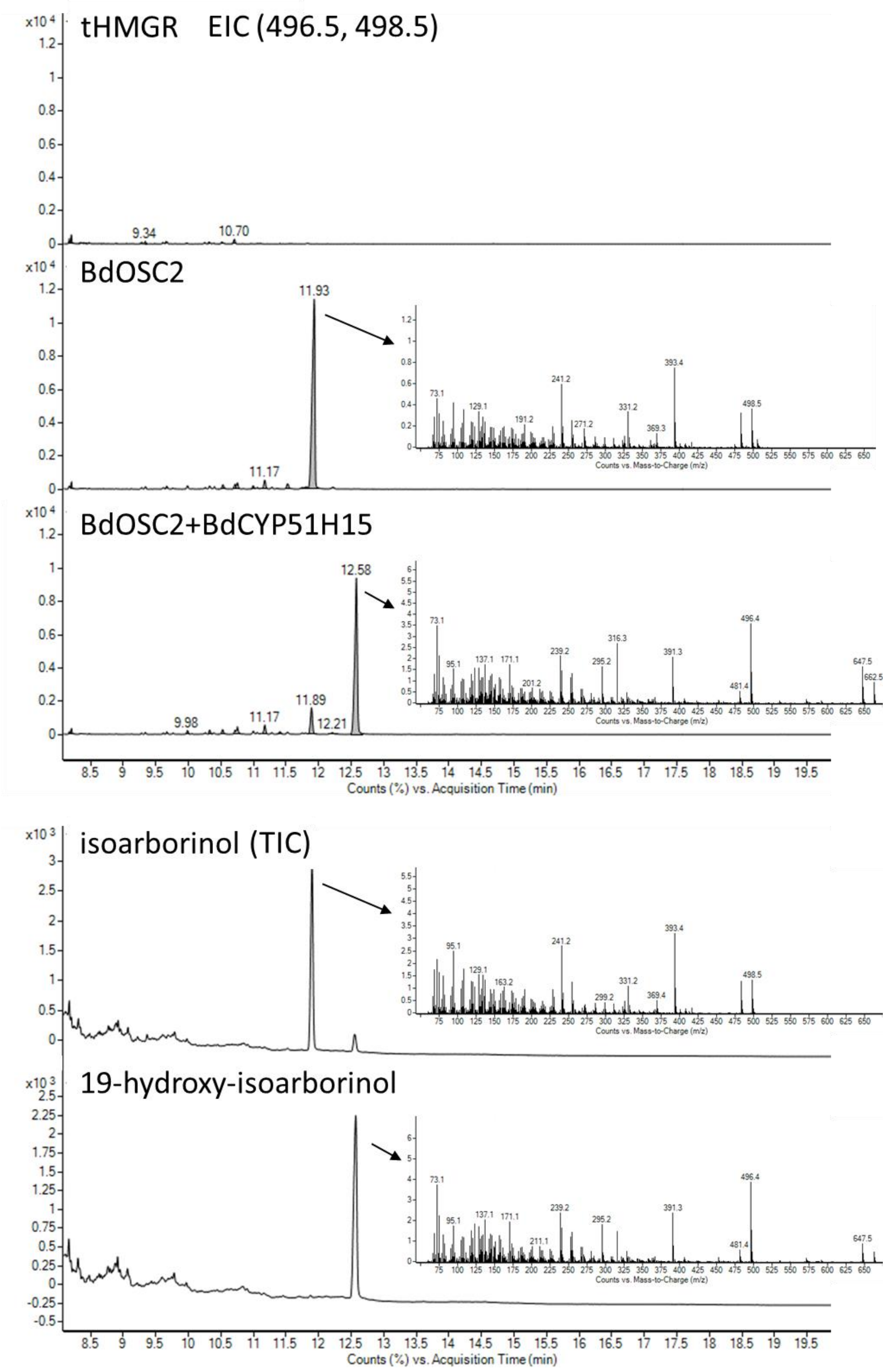


**Supplementary Fig. 28. GC-MS analysis of *B. distachyon* genes BdOSC2 and BdCYP51H15 expression in *N. benthamiana***. Extracted ion chromatograms for ions representing isoarborinol (498.5, Rt 11.93), and 19-hydroxy-isoarborinol brachynacin (496.5, Rt 12.58) and mass spectra at retention times of product peaks are shown. All gene expression combinations included oat tHMGR. Products were identified based on comparison to isoarborinol and 19-hydroxy-isoarborinol purified from *N. benthamiana*.

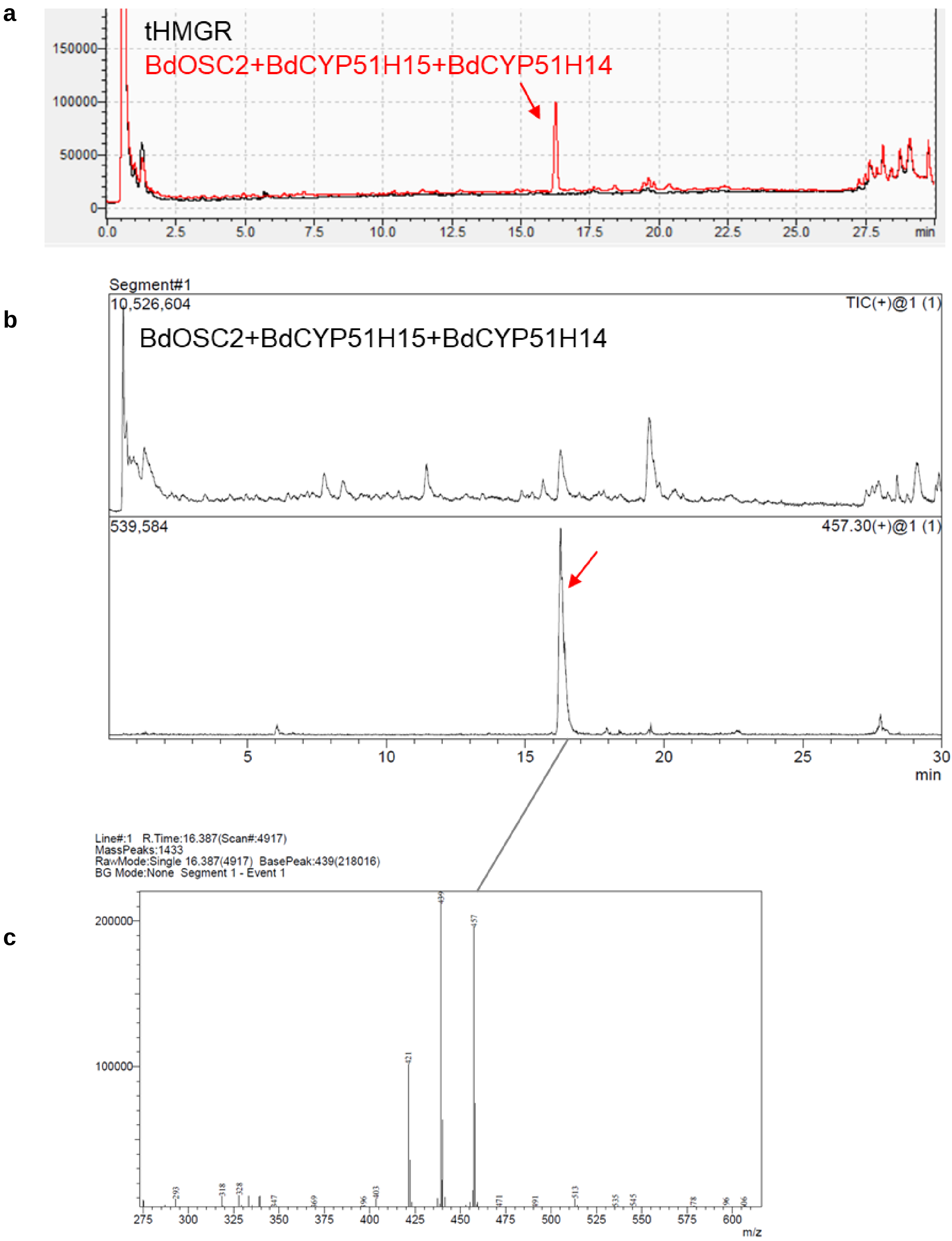


**Supplementary Fig. 29. LC-MS detection of 7,19,28-trihydroxy-isoarborinol molecular ion**. **a**, CAD chromatogram. **b**, MS chromatogram- TIC and EIC of 457.3. **c**, mass spectra at peak retention time.

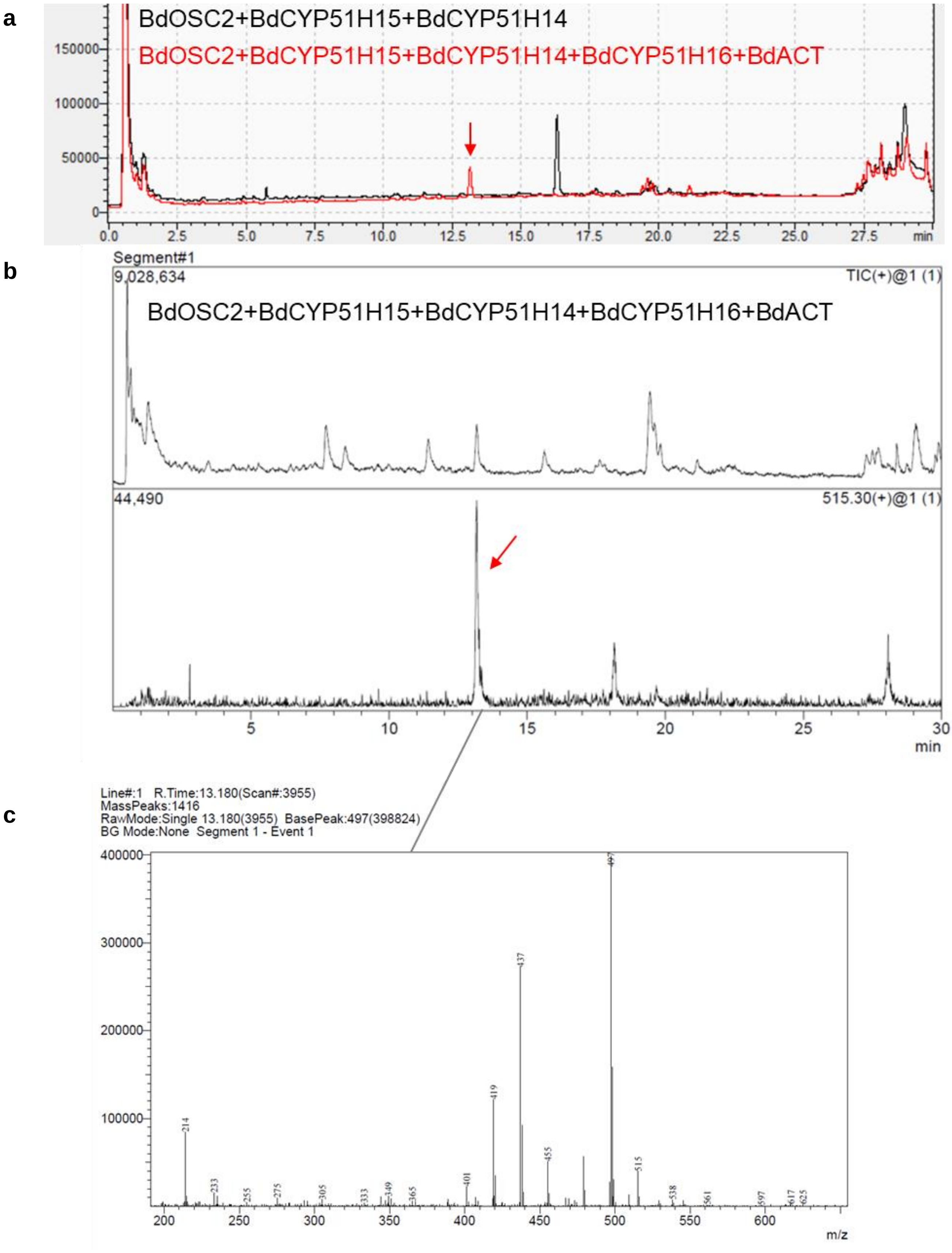


**Supplementary Fig. 30. LC-MS detection of brachynacin molecular ion**. **a**, CAD chromatogram, **b**, MS chromatogram- TIC and EIC of 515.3. **c**, mass spectra at peak retention time.

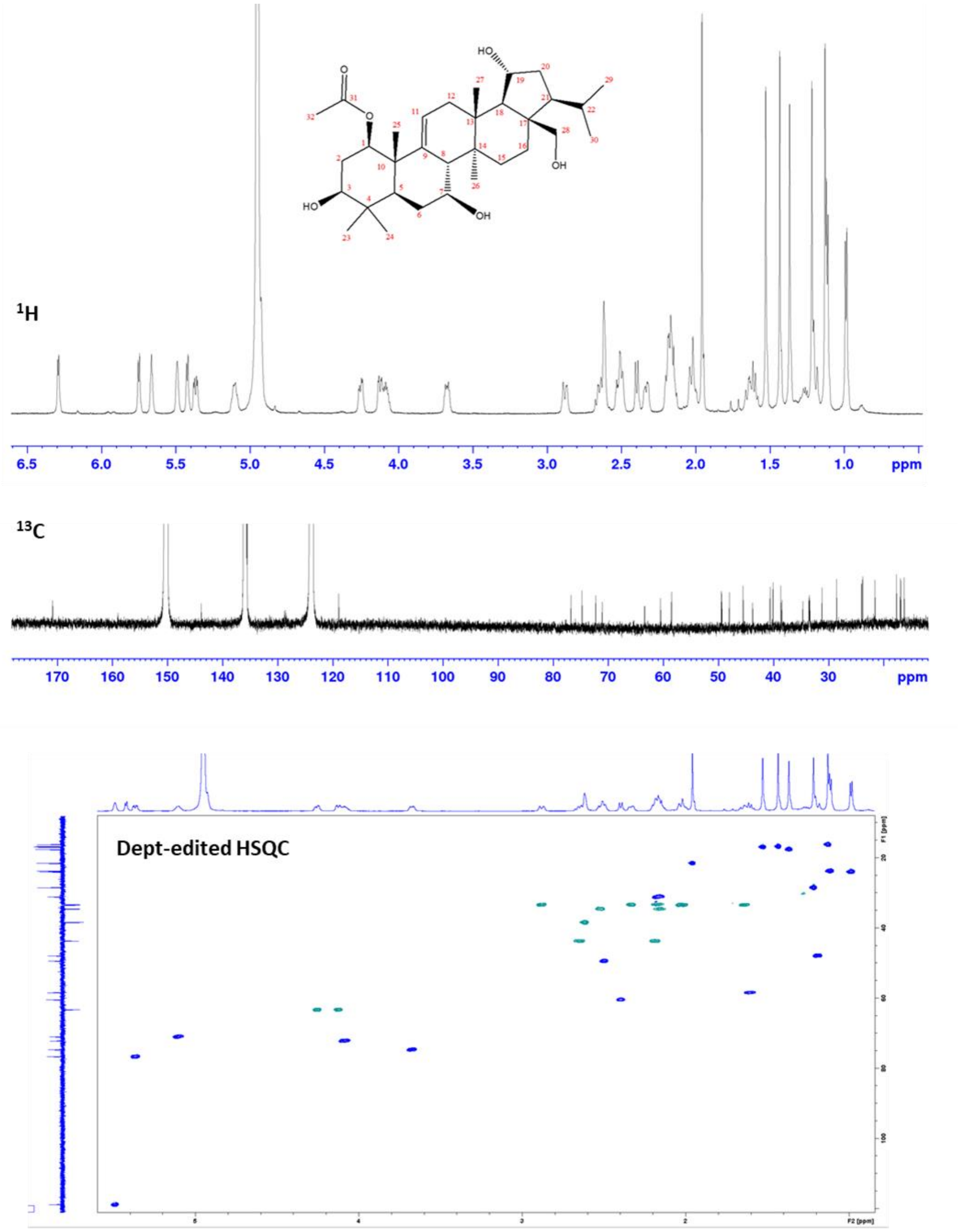


**Supplementary Fig. 31. $^{1}$H, $^{13}$C, and dept-edited HSQC spectra for brachynacin** (**Pyridine-d5**). Referenced to residual solvent peak ($^{1}$H δ: 8.74) ($^{13}$C δ: 150.3)]. 600 MHz instrument.

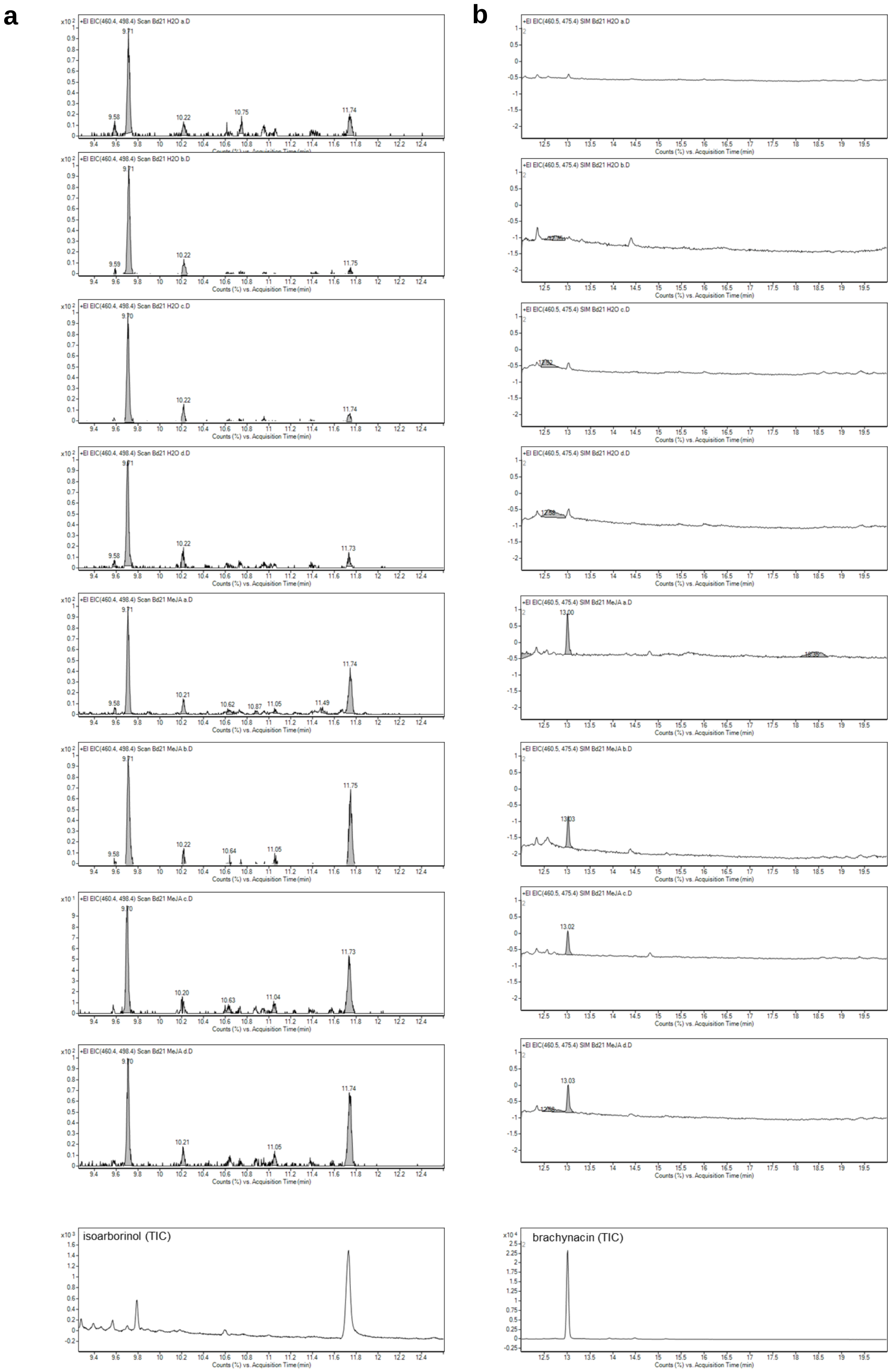


**Supplementary Fig. 32. GC-MS analysis of TMS-derivatized extracts from *B. distachyon* leaves treated with methyl jasmonate (MeJa), or $H_2O$ (control) for 12 hours**. Extracted ion chromatograms are for ions representing **a**, isoarborinol (498.4, Rt 11.73), 5α-cholestan-3β-ol (460.4, Rt 9.70) and **b**, brachynacin (475.4, Rt 13.02). Y-axes are linked to peak of internal standard- 5α-cholestan-3β-ol. Extracts from four biological replicates are shown. *B. distachyon* extracts were compared to isoarborinol and brachynacin purified from *N. benthamiana* for identification.

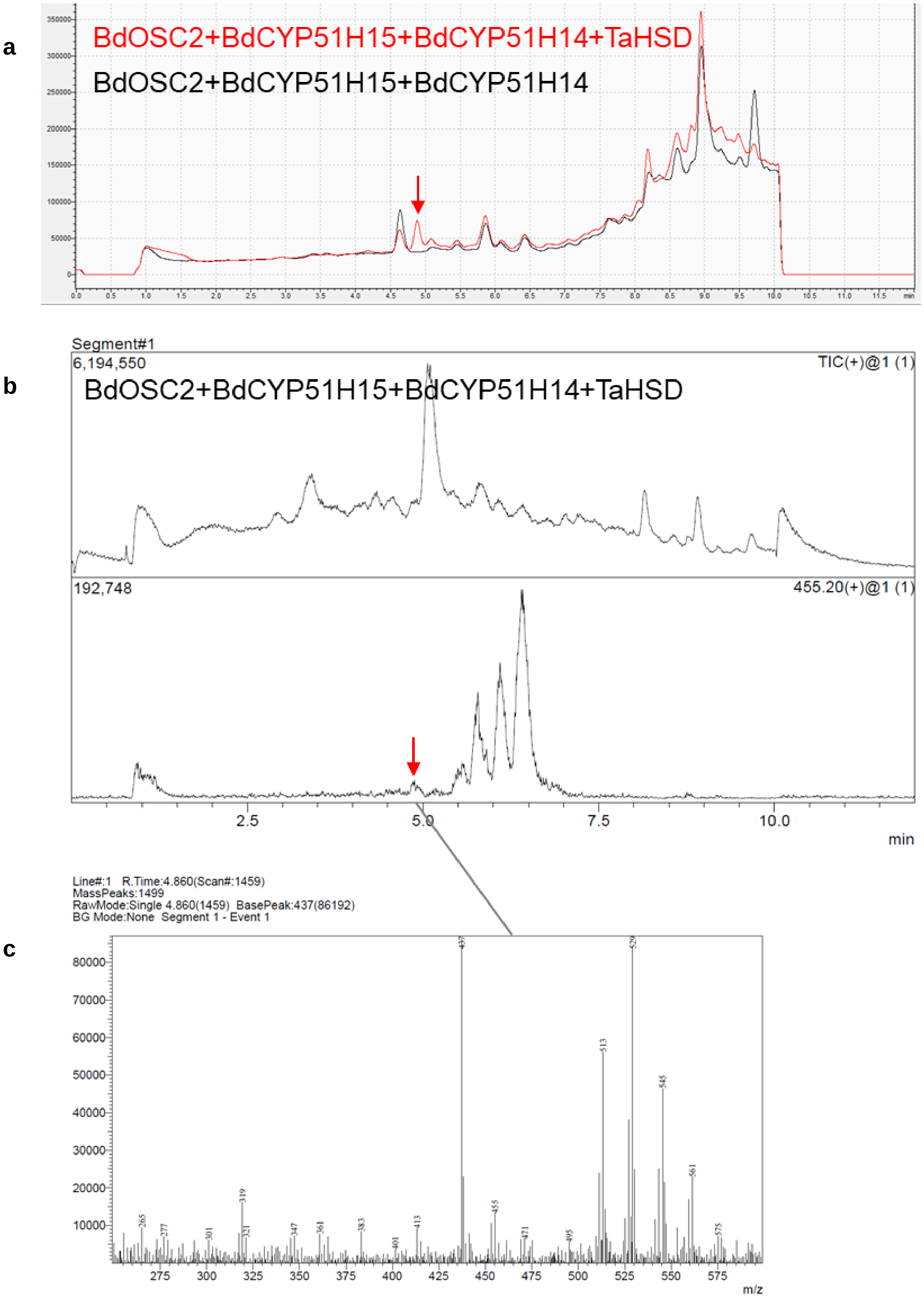


**Supplementary Fig. 33. LC-MS detection of 7,9,28-trihydroxy-isoarborinone molecular ion**. **a**, CAD chromatogram. **b**, MS chromatogram- TIC and EIC of 497.3. **c**, mass spectra at peak retention time.

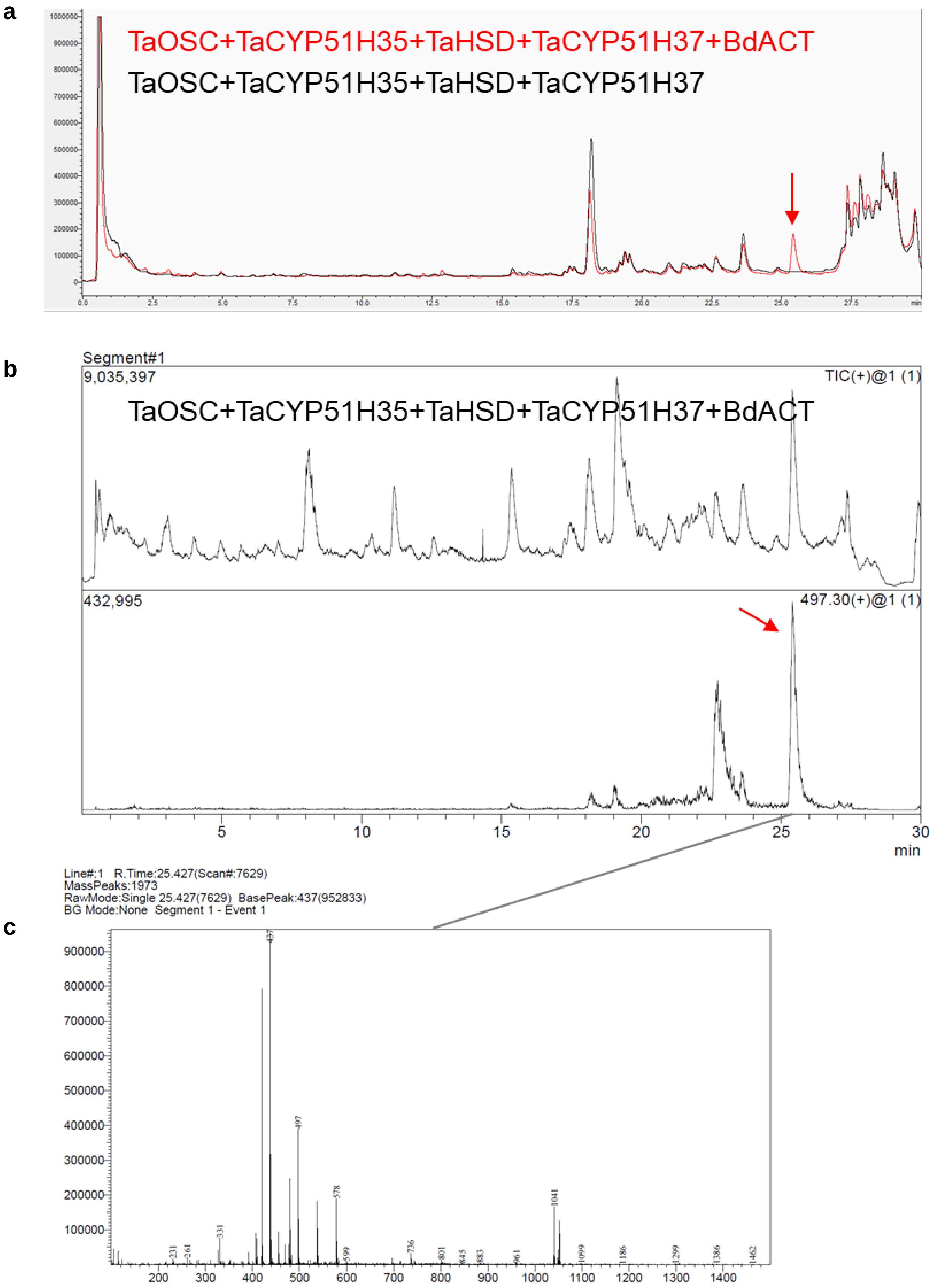


**Supplementary Fig. 34. LC-MS detection of acetyl-ellarinacin molecular ion**. **a**, CAD chromatogram. **b**, MS chromatogram- TIC and EIC of 497.3. **c**, mass spectra at peak retention time.

**Supplementary Table 1.** Genes comprising six wheat BGCs, and the co-expression network modules to which they belong. In bold: genes that are co-expressed (r-val>0.7) with bait genes (asterisked)

| **BGC no.** | **Gene ID** | **Gene Annotation** | **Gene name in this study** | **Co-expression network module** |
|---|---|---|---|---|
| **1(2A)** | TraesCS2A02G027600 | Kaurene synthase | | 8 |
| | TraesCS2A02G027700 | Cytochrome P450 | | 8 |
| | TraesCSU02G252000 | Copalyl diphosphate synthase | | 34 |
| | TraesCSU02G244600 | Copalyl diphosphate synthase | | 34 |
| | TraesCSU02G244600 | Copalyl diphosphate synthase | | 34 |
| | **TraesCSU02G008700** | **Kaurene synthase*** | **TaKSL1** | **25** |
| | **TraesCSU02G008800** | **Cytochrome P450** | | **25** |
| | **TraesCSU02G008900** | **Cytochrome P450** | | **25** |
| | **TraesCSU02G009000** | **Glycosyltransferase** | | **12** |
| | **TraesCSU02G009100** | **Cytochrome P450** | | **25** |
| | **TraesCS2A02G027800** | **Copalyl diphosphate synthase** | **TaCPS2** | **25** |
| | TraesCS2A02G027900 | RING/FYVE/PHD zinc finger superfamily protein | | N/A |
| | **TraesCS2A02G028000** | **Glycosyltransferase** | | **25** |
| | TraesCS2A02G028100 | Kaurene synthase | | N/A |
| | TraesCS2A02G028200 | Copalyl diphosphate synthase | | N/A |
| **1(2D)** | TraesCS2D02G029300 | Cytochrome P450 | | 8 |
| | TraesCS2D02G029400 | Kaurene synthase | | 34 |
| | TraesCS2D02G029500 | Cytochrome P450 | | 34 |
| | **TraesCS2D02G029600** | **Copalyl diphosphate synthase** | **TaCPS-D2** | **25** |
| | **TraesCS2D02G029700** | **Cytochrome P450** | | **12** |
| | **TraesCS2D02G029800** | **Glycosyltransferase** | | **25** |
| | **TraesCS2D02G029900** | **Cytochrome P450** | | **25** |
| | **TraesCS2D02G030000** | **Cytochrome P450** | | **25** |
| | **TraesCS2D02G030100** | **Kaurene synthase*** | **TaKSL-D1** | **25** |
| | TraesCS2D02G030200 | Kaurene synthase | TaKSL4 | N/A |
| | TraesCS2D02G030300 | Copalyl diphosphate synthase | | N/A |
| **2(2B)** | TraesCS2B02G444900 | Kaurene synthase | | N/A |
| | TraesCS2B02G445000 | Cytochrome P450 | | N/A |
| | **TraesCS2B02G445100** | **Kaurene synthase** | **TaKSL3** | **25** |
| | **TraesCS2B02G445200** | **Kaurene synthase*** | **TaKSL2** | **25** |
| | **TraesCS2B02G445300** | **Cytochrome P450** | | **25** |
| | TraesCS2B02G445400 | Cytochrome P450 | | 0 |
| | **TraesCS2B02G445500** | **Ent-copalyl diphosphate synthase** | **TaCPS1** | **25** |
| | **TraesCS2B02G445600** | **Cytochrome P450** | | **25** |
| | TraesCS2B02G445700 | Ent-kaurene synthase | | 0 |
| | TraesCS2B02G445800 | Kaurene synthase | | N/A |
| | TraesCS2B02G445900 | Kaurene synthase | | N/A |
| **3(5A)** | **TraesCS5A02G004500** | **Cytochrome P450-like protein** | **TaCYP51H35_5A** | **34** |
| | **TraesCS5A02G004600** | **Cytochrome P450-like protein** | **TaCYP51H13_5A** | **25** |
| | **TraesCS5A02G004700** | **Cytochrome P450-like protein** | **TaCYP51H37_5A** | **25** |
| | **TraesCS5A02G004800** | **Hydroxysteroid dehydrogenase, putative** | **TaHSD_5A** | **25** |
| | **TraesCS5A02G004900** | **Terpene cyclase/mutase family member*** | **TaOSC_5A** | **25** |
| **3(5D)** | **TraesCS5D02G011800** | **Terpene cyclase/mutase family member*** | **TaOSC (TaIAS)** | **25** |
| | **TraesCS5D02G011900** | **Hydroxysteroid dehydrogenase, putative** | **TaHSD (TaHID)** | **25** |
| | **TraesCS5D02G012000** | **Cytochrome P450-like protein** | **TaCYP51H37_5D (TaHIO)** | **25** |
| | **TraesCS5D02G012100** | **Cytochrome P450-like protein** | **TaCYP51H13P_5D** | **25** |
| | **TraesCS5D02G012200** | **Obtusifoliol 14-alpha demethylase** | **TaCYP51H13P_5D** | **25** |
| | **TraesCS5D02G012300** | **Cytochrome P450-like protein** | **TaCYP51H35_5D (TaIAH)** | **34** |
| **4(5D)** | **TraesCS5D02G488300** | **O-methyltransferase-like protein** | **TaOMT6** | **25** |
| | **TraesCS5D02G488400** | **O-methyltransferase-like protein** | **TaOMT7** | **25** |
| | **TraesCS5D02G488500** | **Cytochrome P450** | **TaCYP71C164_5D** | **25** |
| | TraesCS5D02G488600 | Chalcone synthase | | 0 |
| | **TraesCS5D02G488700** | **Chalcone synthase*** | **TaCHS1** | **25** |
| | **TraesCS5D02G488800** | **O-methyltransferase** | **TaOMT3** | **25** |
| | **TraesCS5D02G488900** | **O-methyltransferase** | **TaOMT8** | **25** |
| | TraesCS5D02G489000 | Chalcone-flavanone isomerase family protein | chi-D1 | 10 |

**Supplementary Table 2.** Gene expression values of 3-5A and 3-5D cluster paralogs

| Gene accession | Annotation | Average expression (tpm) | Max expression (tpm) |
|---|---|---|---|
| TraesCS5D02G006200 | Cytochrome P450-like protein | 0.112±0.035 | 8.896 |
| TraesCS5D02G006300 | Hydroxysteroid dehydrogenase, putative | 0.009±0.003 | 0.792 |
| TraesCS5D02G006400 | Cytochrome P450-like protein | 0±0 | 0.018 |
| TraesCS5D02G006500 | Cytochrome P450-like protein | 0.018±0.004 | 0.963 |
| TraesCS5D02G006600 | Terpene cyclase/mutase family member | 0.009±0.004 | 1.257 |
| TraesCS5D02G006700 | Terpene cyclase/mutase family member | 0.011±0.003 | 0.566 |
| TraesCS5A02G005600 | Cytochrome P450-like protein | 0.006±0.001 | 0.186 |
| TraesCS5A02G005900 | Terpene cyclase/mutase family member | 0.025±0.01 | 2.936 |
| TraesCS5A02G006000 | Cytochrome P450-like protein | 0.0075±0.003 | 0.897 |
| TraesCS5B01G004500 | Cytochrome P450-like protein | 0.148±0.031 | 5.928 |
| TraesCS5B01G004800 | Terpene cyclase/mutase family member | 0.005±0.001 | 0.213 |
| TraesCS5B01G004900 | Cytochrome P450-like protein | 0.669±0.2334 | 61.012 |
| TraesCS5B01G004600 | Cytochrome P450-like protein | 0.250±0.0779 | 24.462 |

Average and maximal normalized gene expression values of TaOSC, TaHSD and TaCYP51H35/37 paralogs, across 379 datapoints in http://www.wheat-expression.com transcriptomes dataset. tpm, transcripts per million.

**Supplementary Table 3.** $^{13}C$ δ of **19-hydroxy-isoarborinol** in this work [100 MHz] compared to the literature (**rubiarbonol K**) [125 MHz]. P**yridine-d5** [referenced to the most downfield peak reported in the literature] [6].

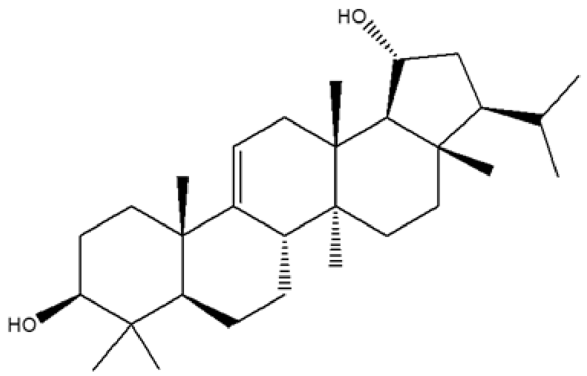


| $^{13}C$ δ This work | $^{13}C$ δ Lit | Δ ($^{13}C$ δ This work - Lit) |
|---|---|---|
| 148.8 | 148.8 | 0.0 |
| 114.9 | 114.9 | 0.0 |
| 78.1 | 78.1 | 0.0 |
| 70.1 | 70.2 | -0.1 |
| 59.0 | 59.0 | 0.0 |
| 57.7 | 57.8 | -0.1 |
| 52.9 | 52.9 | 0.0 |
| 44.0 | 44.1 | -0.1 |
| 41.9 | 42.0 | -0.1 |
| 41.1 | 41.1 | 0.0 |
| 39.9 | 40.0 | -0.1 |
| 39.7 | 39.8 | -0.1 |
| 38.4 | 38.5 | -0.1 |
| 37.6 | 37.6 | 0.0 |
| 37.2 | 37.2 | 0.0 |
| 36.6 | 36.6 | 0.0 |
| 36.5 | 36.6 | -0.1 |
| 30.7 | 30.8 | -0.1 |
| 29.9 | 29.9 | 0.0 |
| 28.9 | 29.0 | -0.1 |
| 28.7 | 28.7 | 0.0 |
| 27.1 | 27.2 | -0.1 |
| 23.2 | 23.2 | 0.0 |
| 22.4 | 22.5 | -0.1 |
| 22.2 | 22.3 | -0.1 |
| 21.9 | 21.9 | 0.0 |
| 17.5 | 17.6 | -0.1 |
| 16.8 | 16.8 | 0.0 |
| 16.6 | 16.6 | 0.0 |
| 15.9 | 15.9 | 0.0 |

**Supplementary Table 4.** $^{13}C$ & $^{1}H$ δ assignments for **ellarinacin**. **Pyridine-d5** [referenced to residual solvent peak ($^{1}H$ δ: 8.74) ($^{13}C$ δ: 150.3)]. Assignments were made via a combination of $^{1}H$, $^{13}C$, DEPT-edited HSQC, HMBC, COSY and 2D NOESY experiments. Where signals overlap $^{1}H$ δ is reported as the centre of the respective HSQC crosspeak. C3-C2 epoxide was assigned as beta due to an NOE observed between C2-H and C5-H.

**Carbon numbering scheme and selected COSY, HMBC and NOESY**

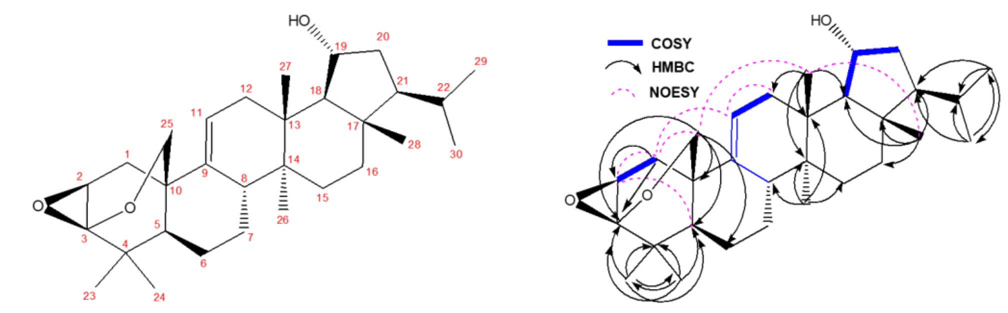


| Carbon # | $^{13}C$ δ (100 MHz) | $^{1}H$ δ (400 MHz) | Carbon # | $^{13}C$ δ (100 MHz) | $^{1}H$ δ (400 MHz) |
|---|---|---|---|---|---|
| **9** | 139.69 | / | **13** | 38.37 | / |
| **11** | 123.34 | 5.69 (1H, m) | **12** | 38.34 | 2.47 (2H, m) |
| **3** | 98.76 | / | **16** | 37.50 | 1.67 (2H, m) |
| **25** | 72.56 | 4.51 (1H, dd *J* = 8.5, 3.1)<br>3.65 (1H, dd *J* = 8.5, 1.0) | **1** | 37.08 | 2.33 (1H, m)<br>2.21 (1H, m) |
| **2** | 72.21 | 3.92 (1H, td *J* = 10.7, 3.1) | **15** | 32.11 | 2.98 (1H, m)<br>1.86 (1H, m) |
| **19** | 70.69 | 4.50 (1H, m) | **6** | 31.83 | 2.31 (1H, m)<br>1.50 (1H, m) |
| **18** | 59.45 | 2.04 (1H, m) | **22** | 31.13 | 1.41 (1H, m) |
| **21** | 58.21 | 1.39 (1H, m) | **7** | 30.87 | 2.40 (1H, m)<br>2.18 (1H, m) |
| **8** | 50.62 | 2.22 (1H, m) | **24** | 28.67 | 1.26 (3H, s) |
| **5** | 48.05 | 1.63 (1H, m) | **29** | 23.61 | 0.86 (3H, d *J* = 5.9) |
| **17** | 44.15 | / | **30** | 22.57 | 0.90 (3H, d *J* = 5.9) |
| **20** | 42.31 | 2.07 (1H, m)<br>1.97 (1H, m) | **23** | 19.93 | 1.40 (3H, s) |
| **4** | 40.44 | / | **27** | 17.21 | 1.10 (3H, s) |
| **14** | 40.29 | / | **26** | 16.45 | 1.31 (3H, s) |
| **10** | 39.00 | / | **28** | 16.39 | 0.90 (3H, s) |

**Supplementary Table 5**. Pairwise alignment of putative proteins from orthologous biosynthetic gene clusters in common wheat, *Aegilops tauschii*, and wild emmer wheat.

| ***T. aestivum* (Chr.5D)** | ***A. tauschii*** | **% identity** |
|---|---|---|
| TraesCS5D02G011800<br>TaOSC_5D | AET5Gv20013100 | 99.9 |
| TraesCS5D02G011900<br>TaHSD_5D | AET5Gv20013000 | 100 |
| TraesCS5D02G012000<br>TaCYP51H37_5D | AET5Gv20012900 | 100 |
| TraesCS5D02G012100/<br>TraesCS5D02G012200<br>TaCYP51H13P_5D | AET5Gv20012800 | 99.4 |
| TraesCS5D02G012300<br>TaCYP51H35_5D | AET5Gv20012700 | 99.6 |

| ***T. aestivum* (Chr.5A)** | ***T. turgidum ssp. diccocoides*** | **% identity** |
|---|---|---|
| TraesCS5A02G004500<br>TaCYP51H35_5A | TRIDC5AG000720 | 99.6 |
| TraesCS5A02G004600<br>TaCYP51H13_5A | TRIDC5AG000730 | 99.4 |
| TraesCS5A02G004700<br>TaCYP51H37_5A | TRIDC5AG000740 | 100 |
| TraesCS5A02G004800<br>TaHSD_5A | TRIDC5AG000750 | 99.7 |
| TraesCS5A02G004900<br>TaOSC_5A | TRIDC5AG000760 | 99.9 |

**Supplementary Table 6.** $^{13}$C & $^{1}$H δ assignments for **brachynacin**. **Pyridine-d5** [referenced to residual solvent peak ($^{1}$H δ: 8.74) ($^{13}$C δ: 150.3)]. Assignments were made via a combination of $^{1}$H, $^{13}$C, DEPT-edited HSQC, HMBC, COSY, TOCSY and 2D ROESY experiments. Where signals overlap $^{1}$H δ is reported as the centre of the respective HSQC crosspeak. C1-OAc was assigned as beta due to NOEs observed between C1-H and C3-H, and C5-H. C7-OH was assigned as beta due to NOEs observed between C7-H and C26-$H_3$ and C5-H.

**Carbon numbering scheme and selected COSY, HMBC and TOCSY**

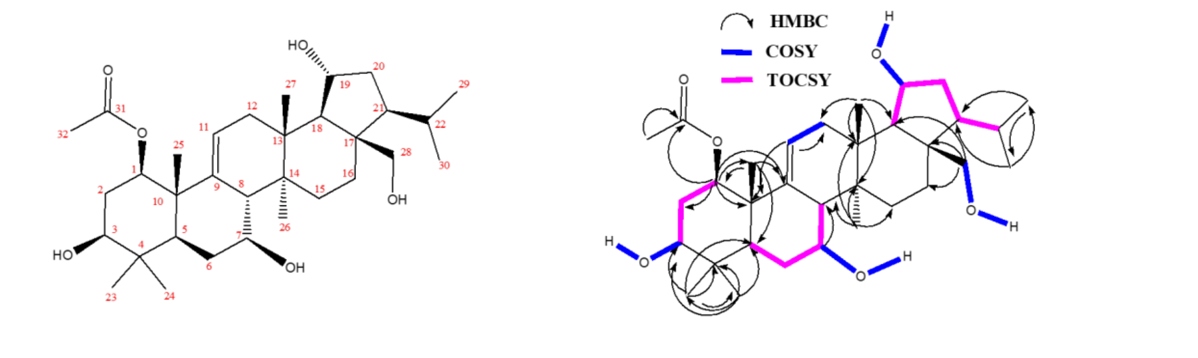


| Carbon # | $^{13}$C δ (150 MHz) | $^{1}$H δ (600 MHz) | Carbon # | $^{13}$C δ (150 MHz) | $^{1}$H δ (600 MHz) |
|---|---|---|---|---|---|
| **31** | 170.84 | / | **4** | 40.09 | / |
| **9** | 143.87 | / | **13** | 38.66 | / |
| **11** | 118.92 | 5.49 (1H, m) | **12** | 38.47 | 2.62 (2H, m) |
| **1** | 76.75 | 5.37 (1H, dd *J* = 11.5, 4.3) | **2** | 34.67 | 2.52 (1H, m)<br>2.16 (1H, m) |
| **3** | 74.75 | 3.67 (1H, m) | **6** | 33.56 | 2.33 (1H, m)<br>2.18 (1H, m) |
| **7** | 72.26 | 4.09 (1H, m) | **15** | 33.50 | 2.03 (1H, m)<br>1.64 (1H, m) |
| **19** | 71.07 | 5.11 (1H, m) | **16** | 33.43 | 2.88 (1H, m)<br>2.03 (1H, m) |
| **28** | 63.38 | 4.26 (1H, dd *J* = 11.1, 4.2)<br>4.12 (1H, dd *J* = 11.1, 4.2) | **22** | 31.18 | 2.17 (1H, m) |
| **18** | 60.50 | 2.40 (1H, d *J* = 9.9) | **23** | 28.56 | 1.22 (3H, s) |
| **21** | 58.50 | 1.61 (1H, m) | **30** | 24.03 | 0.99 (3H, d *J* = 6.5) |
| **8** | 49.47 | 2.50 (1H, m) | **29** | 23.81 | 1.11 (3H, d *J* = 6.5) |
| **17** | 49.38 | / | **32** | 21.59 | 1.96 (3H, s) |
| **5** | 48.00 | 1.20 (1H, m) | **26** | 17.67 | 1.34 (3H, s) |
| **10** | 45.51 | / | **27** | 16.99 | 1.53 (3H, s) |
| **20** | 43.80 | 2.65 (1H, m)<br>2.19 (1H, m) | **25** | 16.84 | 1.43 (3H, s) |
| **14** | 40.62 | / | **24** | 16.29 | 1.13 (3H, s) |

**Exchangable Protons (Assigned by COSY)**

**C3-OH:** δ 6.19 (1H, d, *J* = 5.2); **C7-OH:** δ 5.75 (1H, d, *J* = 6.2)**; C28-OH:** δ 5.66 (1H, brt, *J* = 3.9), **C19-OH:** δ 5.42 (1H, d, J= 5.8).

**Supplementary Table 7.** *B. distachyon* putative terpene cluster homologous to wheat BGC 2(2B)

| Gene ID | Location | Description |
|---|---|---|
| Bradi5g21383 | Bd5:24179552..24182576 (+ strand) | farnesyl diphosphatase/FPP phosphatase |
| Bradi5g21387 | Bd5:24182863..24188013 (+ strand) | ent-kaurene synthase |
| Bradi5g21400 | Bd5:24188931..24190869 (- strand) | cytochrome P450 |
| Bradi5g21410 | Bd5:24194057..24196081 (- strand) | cytochrome P450 |
| Bradi5g21420 | Bd5:24200947..24203100 (- strand) | germacrene A alcohol dehydrogenase |
| Bradi5g21430 | Bd5:24205464..24207731 (- strand) | germacrene A alcohol dehydrogenase |
| Bradi5g21440 | Bd5:24210985..24218157 (+ strand) | ent-kaurene synthase |
| Bradi5g21447 | Bd5:24218971..24221276 (- strand) | cytochrome P450 |
| Bradi5g21460 | Bd5:24222480..24225443 (- strand) | cytochrome P450 |
| Bradi5g21465 | Bd5:24227072..24227779 (+ strand) | unknown protein |
| Bradi5g21470 | Bd5:24229095..24230866 (- strand) | cytochrome P450 |
| Bradi5g21480 | Bd5:24238153..24242304 (+ strand) | ent-kaurene synthase |
| Bradi5g21488 | Bd5:24244530..24246038 (+ strand) | cytochrome P450 |
| Bradi5g21492 | Bd5:24246237..24247028 (+ strand) | unknown protein |
| Bradi5g21497 | Bd5:24270996..24274843 (+ strand) | ent-kaurene synthase |

**Supplementary Table 8.** Oligonucleotides used in this study

| | |
|---|---|
| **Full CDS cloning** | |
| TaCYP51H35_5D F | GCGCCGTCTCGCTCGAATGGACTTAGCAAGTCTC |
| TaCYP51H35_5D R | GCGCCGTCTCGCTCGAAGCCTACAAAATGCCATTCCT |
| TaCYP51H37_5D F | GCGCCGTCTCGCTCGAATGGAGATGGCAAGTAGCGC |
| TaCYP51H37_5D R | GCGCCGTCTCGCTCGAAGCCTAGCCTAGCAGCTGGCGCCTCTTGTAG |
| TaCYP51H13_5A F | GCGCCGTCTCGCTCGAATGGACTTGACAAGTCTCACTACG |
| TaCYP51H13_5A R | GCGCCGTCTCGCTCGAAGCTTAAATTCCATTTCTCGTATATCTCA |
| A. tauschii IAH F | GGGGACAAGTTTGTACAAAAAAGCAGGCTATGGACTTAGCAAGTCTCAC |
| A. tauschii IAH R | GGGGACCACTTTGTACAAGAAAGCTGGGTCTAATTAAGTGACTGCAAAATGC |
| BdOSC2 F | GCGCCGTCTCGCTCGAATGTGGAAGCTAAAGATCGCA |
| BdOSC2 R | GCGCCGTCTCGCTCGAAGCTTATGCCTTTTGTGCTTGCAG |
| BdCYP51H14 F | GCGCCGTCTCGCTCGAATGTTCATGACAAGTAGCGCC |
| BdCYP51H14 R | GCGCCGTCTCGCTCGAAGCCTAGCCCAACAGCCGATGTC |
| BdCYP51H15 F | GCGCCGTCTCGCTCGAATGGACTTGGCAAGCACAGC |
| BdCYP51H15 R | GCGCCGTCTCGCTCGAAGCCTAAGTGCTAGCACGGCAGC |
| BdCYP51H16 F | GCGCCGTCTCGCTCGAATGGAATTTACAAGTGGCGAC |
| BdCYP51H16 R | GCGCCGTCTCGCTCGAAGCCTAGGCTGACATCCTCGATC |
| TaCYP71C164_5D F | GGGGACAAGTTTGTACAAAAAAGCAGGCTATGGAAGATCTCGTGAAGAAACC |
| TaCYP71C164_5D R | GGGGACCACTTTGTACAAGAAAGCTGGGTCTACATCCAGGATTTTGGAATTAACAATAG |
| TaCYP71F53_5D F | GGGGACAAGTTTGTACAAAAAAGCAGGCTATGGAGGGTTGGTTAACCTTATGTTTC |
| TaCYP71F53_5D R | GGGGACCACTTTGTACAAGAAAGCTGGGTCTATATAGTGGAGCGTACATATGGAATAGC |
| TaOMT6 F | GGGGACAAGTTTGTACAAAAAAGCAGGCTATGGCGCCCACCCAAGGCAAGCAGAGTTCTCA |
| TaOMT6 R | GGGGACCACTTTGTACAAGAAAGCTGGGTTCAAGGGTAGAGCTCAATAACAGATCTAACTC |
| TaOMT3 F | GGGGACAAGTTTGTACAAAAAAGCAGGCTATGGGCTCCACTGCCGTGGAGAAGGTC |
| TaOMT3 R | GGGGACCACTTTGTACAAGAAAGCTGGGTCTATTTGACGAACTCAATGACCCATGC |
| TaOMT8 F | GGGGACAAGTTTGTACAAAAAAGCAGGCTATGGGTTCTATCTCCGACGACGAGGCG |
| TaOMT8 R | GGGGACCACTTTGTACAAGAAAGCTGGGTCTATTTGTTCAACTCAATGACCCATATG |
| ***T. aestivum* RT-PCR** | |
| TaCYP51H35_5D F | GGTGGGCTGTGGCTCTT |
| TaCYP51H35_5D R | CTACAAAATGCCATTCCTCTT |
| TaCYP51H13_5A F | ACCAACATTGATCCACAGC |
| TaCYP51H13_5A R | TTAAATTCCATTTCTCGTATATCTC |
| TaCYP51H37_5D F | CCGCTGCTCCAGCCAGTGA |
| TaCYP51H37_5D R | CTAGCCTAGCAGCTGGCGTCTC |
| TaOSC_5D F | GGGAGTTCGATCCTGCC |
| TaOSC_5D R | CTATCTCTTCCAAGCGAATGTG |
| TaOSC_5A F | TGGAACAACATGGGTATCACATA |
| TaOSC_5A R | CAGGGCAGTCCGCACG |
| ***T. aestivum* qRT-PCR** | |
| TaOSC_5D qRT-PCR F | GGGACTGCATATCGAGGGAA |
| TaOSC_5D qRT-PCR R | CCCCAAGCAATCTCAAAGCA |
| TaHSD qRT-PCR F | GCCTACTTCTCTGTCGACGT |
| TaHSD qRT-PCR R | GTTGTTGGTATGATCCGCCG |
| TaCYP51H35_5D qRT-PCR F | TGGAAGACACATTTGCACCG |
| TaCYP51H35_5D qRT-PCR R | CGAGCTCAAAGTTCCTCAGC |
| TaCYP51H37_5D qRT-PCR F | GCTGAACCCACCAACAACAA |
| TaCYP51H37_5D qRT-PCR R | GTCTTGTCCGCACTGTGAAA |
| TUBB qPCR F | CAAGGAGGTGGACGAGCAGATG |
| TUBB qPCR R | GACTTGACGTTGTTGGGGATCCA |
| ***B. distachyon* RT-PCR** | |
| BdOSC1 F | CATCAGAAGGAGATTCGGAGATA |
| BdOSC1 R | CCATCGTCATTCATCAGTGATAA |
| BdOSC2 F | CATCAGAAAGAGATGCGGAGATA |
| BdOSC2 R | CCATCCTCGTTCATCAGAGATAAC |
| BdCYP51H14 F | ATGCATACTTCAACAAGGATCTAT |
| BdCYP51H14 R | CCTTGATGCAACTATGGAGTGTA |
| BdACT F | ACAAGAATCACATGTGCATTCC |
| BdACT R | GAATCAAGTAGTGCTGGCGT |
| BdCYP51H15 F | CTCTTCTTCTCATCACTGCTTTAG |
| BdCYP51H15 R | TTGACTACAATAGGACTCGCTAC |
| BdCYP51H16 F | ATGGAATTTACAAGTGGCGAC |
| BdCYP51H16 R | CTGCCAAGGTATTGTAGTCGA |
| BdGAPDH F | ATGGGCAAGATTAAGATCGGAA |
| BdGAPDH R | TTACTGAGTCTTGGCCATGT |
| ***B. distachyon* qRT-PCR** | |
| BdOSC1 F | CAGGGGCTGGTGTTATTCAA |
| BdOSC1 R | GTAATCTGCAGCCTTTCGGA |
| BdOSC2 F | CTCCTGAATTGGCTGGTGAG |
| BdOSC2 R | GCCATCCTCGTTCATCAGAG |
| BdCYP51H14 F | GCTGCTCTCTGAAATCGTGA |
| BdCYP51H14 R | GGCCGTCTTTGTACTTGGAA |
| BdACT F | AATCACATGTGCATTCCGGT |
| BdACT R | GCACTCTTTATCGTCTCGGC |
| BdCYP51H15 F | TCGGTCTCCTATTTGCTGGA |
| BdCYP51H15 R | ACATTGGGTGGCTAAGCAAA |
| BdCYP51H16 F | CTCCTTGGACTTCTACACGC |
| BdCYP51H16 R | CACCTTTTGTCCAAGCAAGC |
| BdGAPDH F | TTGCTCTCCAGAGCGATGAC |
| BdGAPDH R | CTCCACGACATAATCGGCAC |
| **Site-directed mutagenesis** | |
| TaOSC_5D(I581S) F | GTACCCCAAACACAGTCGCTTGGAAGAG |
| TaOSC_5D(I581S) R | CTCTTCCAAGCGACTGTGTTTGGGGTAC |

**Supplementary Table 9.** Isolera Prime gradient conditions

| Compound | Column | Solvents | Gradient |
|---|---|---|---|
| Ellarinacin | SNAP Ultra 50 gr | A: hexane<br>B: ethyl acetate | 0-100% B (10 CV)<br>100-100% B (1 CV) |
| | KP- sil 25 gr | A: hexane<br>B: ethyl acetate | 50-70% B (60 CV) |
| | SNAP Ultra 10 gr | A: hexane<br>B: ethyl acetate | 0-70% B (117 CV) |
| Brachynacin | SNAP Ultra 50 gr | A: hexane<br>B: ethyl acetate | 0-100% B (10 CV)<br>100-100% B (1 CV) |
| | Sfar silica D 30 gr | A: hexane<br>B: ethyl acetate | 10-100% B (43 CV)<br>100-100% B (5 CV) |
